# The tissue-resident regulatory T cell pool is shaped by transient multi-tissue migration and a conserved residency program

**DOI:** 10.1101/2023.08.14.553196

**Authors:** Oliver Burton, Orian Bricard, Samar Tareen, Vaclav Gergelits, Simon Andrews, Carlos P. Roca, Carly Whyte, Steffie Junius, Aleksandra Brajic, Emanuela Pasciuto, Magda Ali, Pierre Lemaitre, Susan M. Schlenner, Harumichi Ishigame, Brian D. Brown, James Dooley, Adrian Liston

## Abstract

The tissues are the site of many of the most important immunological reactions, yet the immunology of the tissues has remained relatively opaque. Recent studies have identified Foxp3^+^ regulatory T cells (Tregs) in several non-lymphoid tissues. These tissue-resident populations have been ascribed unique characteristics based on comparisons to lymphoid Tregs. Here we performed a systematic analysis of the Treg population residing in non-lymphoid organs throughout the body, revealing shared phenotypes, transient residency and common molecular dependencies. Further, tissue Tregs from different non-lymphoid organs shared T cell receptor (TCR) sequences, with functional capacity to drive multi-tissue Treg entry. Finally, tissue Tregs extracted from non-lymphoid organs were tissue-agnostic on re-entry, without homing preference for their tissue of origin. Together these results demonstrate that the tissue-resident Treg pool in most non-lymphoid organs, other than the gut, is largely constituted by broadly self-reactive Tregs, characterised by transient multi-tissue migration and a common residency program.

## Introduction

The tissues are the key site for pathological reactions across a range of immunological challenges, from infections to autoimmunity and inflammation. Our understanding of the immune component of non-lymphoid tissues is, however, relatively limited, and often extrapolated from functions studied in the lymphoid tissues. An early paradigm for tissue immunology emerged from the biology of tissue macrophages, where generic precursor cells seed tissues at early developmental stages and undergo extensive specialization during indefinite residency ^1^. Fetal γδ T cells provide a lymphocyte counterpart to the tissue residency of myeloid cells ^2^, however αβ T cells were largely exempt from the seeding and specialization model, being defined by their circular migration through the lymphatic system.

Tissue resident memory (Trm) CD8 T cells, and, more recently, CD4 T cells, have broken the circular migration paradigm for αβ T cells. Trm cells are antigen-experienced memory T cells with prolonged or even indefinite residency within the tissue of original antigen-experience ^3^. In infections models in the skin, lung, brain, gut and liver, long-lasting pathogen-reactive clonal populations of T cells are found in the tissue following the resolution of infection, and contribute to protection during secondary infection ^4-11^. In the human context, Trm accumulate at the barrier sites from an early age ^12^, and undergo site-specific clonal expansion and transcriptional adaptations ^13^. In liver transplantation and recovery experiments, Trm from the original transplant can even be recovered in the transplanted organ 10 years later ^14^. Murine Trm express distinct phenotypic markers, including CD69, CD103, CXCR6, CD11a and PD-1 ^15^, thought to enable tissue adaptation and prolonged tissue residency. The residential transcriptional program is initiated and maintained within Trm by the transcription factors Hobit, Blimp1 and Runx3 ^16-19^, providing a molecular basis for this unusual cellular behaviour.

Tissue Tregs are an attractive analogue to the conventional Trm populations. Tregs have been isolated from multiple non-lymphoid tissues, and initiate similar transcriptional modules to those found in Trm. Tissue Treg express at high rates the canonical Trm markers of CD69, CD103, CD11a and PD-1, in addition to markers more commonly associated with circulating memory CD8, such as KLRG1 ^20^. These local Tregs are thought to contribute to immunological tolerance in the tissues; more intriguingly, there is an emerging consensus that they have additional, non-immunological, roles in physiological homeostasis. Adipose Tregs aid insulin sensitivity ^21^ and lipolysis ^22^, muscle Tregs can aid muscle repair after injury ^23^, brain Tregs drive oligodendrocyte precursor cell differentiation into oligodendrocytes ^24^, skin Tregs prevent fibrosis and aid regeneration ^25,26^, and diverse additional physiological responses have been attributed to tissue-resident Tregs across other tissues ^20^. The parallels between Trm and tissue Treg have supported the conceptualisation of tissue Tregs under the seeding and specialisation model of leukocyte tissue residency, although the generation of direct data on migratory behaviour and cross-tissue comparison of tissue Tregs has lagged behind that of their non-regulatory counterparts. Few studies have looked in parallel at Tregs from multiple non-lymphoid tissues, so the degree to which these cells are unique per tissue, or only distinct from the lymphoid tissue comparator group, remains largely undefined.

Here we tested the seed and specialisation model of tissue Tregs through a systematic analysis of the Treg population residing in lymphoid, non-lymphoid and gut-associated tissues throughout the body. Using high parameter flow cytometry and transcriptomics we have found unified phenotypes for tissue Tregs in the non-lymphoid tissues, coupled with common molecular dependencies. Tissue Tregs across multiple non-lymphoid organs were dependent upon KLRG1, BATF and CD11a, while being independent of other canonical tissue-residency markers, including CD69, CD103 and ST2. In contrast to the seed and specialisation model, tissue Tregs from different tissues shared TCR clonality and transcriptional phenotypes. Our data further showed that tissue residency was generally short, on the order of ∼3 weeks, other than in the white adipose and gut-associated tissues. Tissue Treg TCR sequences imparted multi-tissue homing, and extracted non-lymphoid tissue Tregs were tissue agnostic on re-entry. Together, these results suggest that tissue Tregs operate under a different residency paradigm from Trm or tissue-resident macrophages, being characterised by slow percolation through multiple tissues in a pan-tissue adapted state.

## Results

### Tissue Tregs share common phenotypes across the non-lymphoid tissues

To assess the uniqueness of Tregs residing within organs, we profiled the infiltrating population in 48 murine tissues, covering a comprehensive set of lymphoid, non-lymphoid and gut-associated tissues (**Supplementary Figure 1A**). Looking at the 28 major tissue sources, Tregs were largely represented as 10-20% of CD4^+^ T cells across each tissue, with frequencies elevated to ∼50% in the bone-marrow, skin and tongue (**Figure 1A**). In absolute numbers, ∼10^4^-10^7^ Tregs were found in each lymphoid tissue type, ∼10^3^-10^5^ Tregs found in each gut-associated tissue type, and ∼10-10^4^ Tregs found in each non-lymphoid tissue type (**Figure 1B**). Cumulatively, lymphoid tissues contained >98% of Tregs identified across the body, with another 0.3% of Tregs found in the blood, 0.8% of Tregs in the gut-associated tissues, and 0.3% of Tregs in the combined set of all non-lymphoid non-gut tissues (**Figure 1C**).

**Figure 1.**
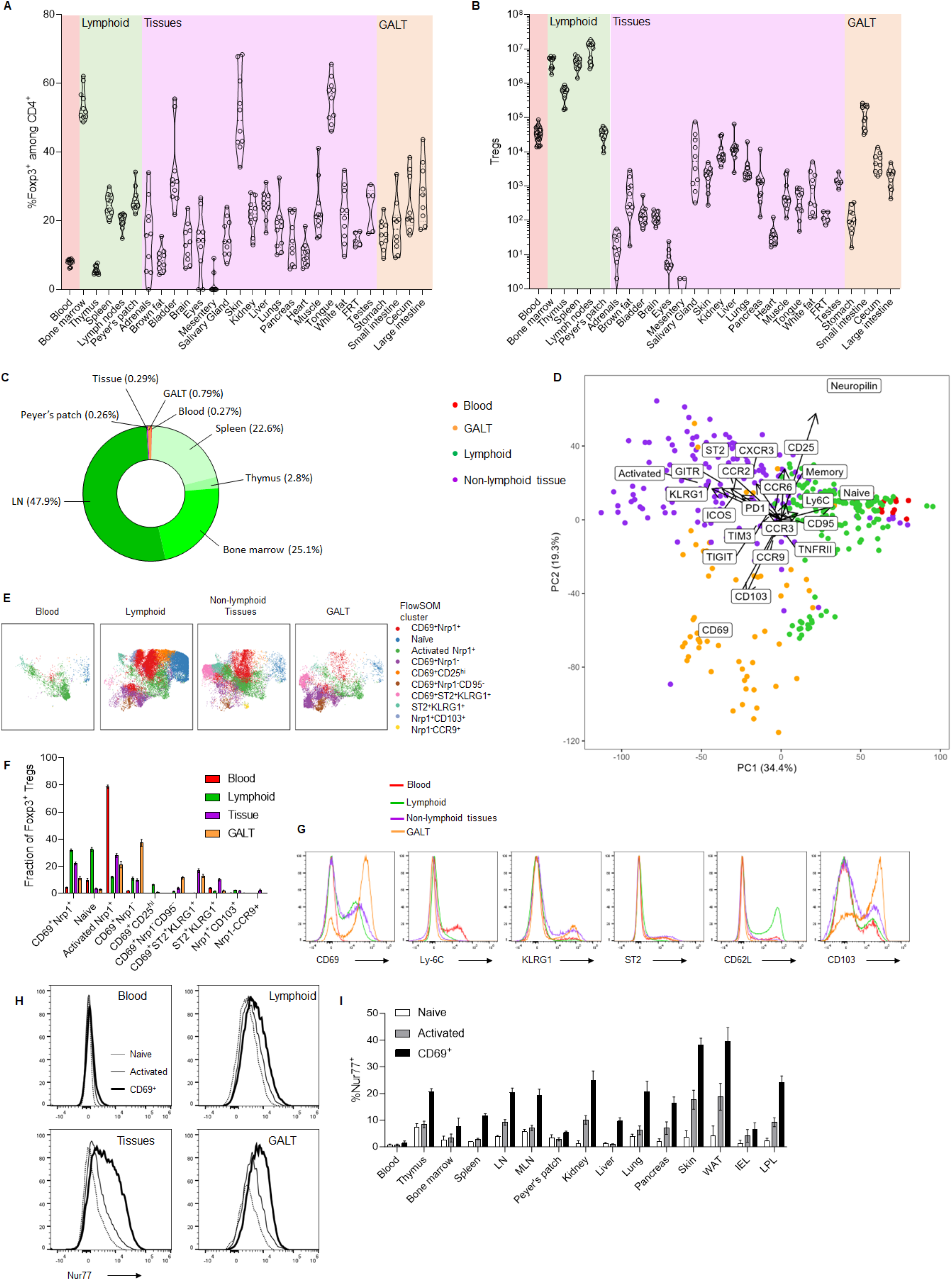
A common phenotypic program unites tissue-residents Tregs across all major organs. Foxp3-Thy1.1 mice, aged 12-20 weeks, were injected with intravenous anti-CD45 antibody label. Major tissues were dissected and lymphocytes purified for flow cytometric analysis of Treg populations. **A)** Frequency of Tregs among CD4^+^ TCRβ^+^ T cells, per organ (n = 9-10), with **B)** absolute number of Tregs recorded per tissue. **C)** Cumulative absolute number of Tregs within major tissues sources and tissue types. **D)** Tregs were phenotyped from 48 organs (n=9-10, classed as blood, lymphoid organs (thymus, spleen, bone marrow, cervical lymph nodes (LN), submandibular LN, axillary LN, inguinal LN, popliteal LN, pancreatic LN, mediastinal LN, renal LN, aortic LN, jejunal LN, colonic LN, duodenal Peyer’s Patches (PP), jejunal Peyer’s Patches, ileal Peyer’s Patches), non-lymphoid organs (skin, lung, pancreas, salivary gland, liver, kidney, tongue, heart, muscle, white fat, brown fat, brain, female reproductive tract (FRT), urethra, testes, prostate, bladder, peritoneum) and gut-associated tissues (stomach IEL, duodenal IEL, Jejunal IEL, ileal IEL, cecal IEL, colonic IEL, stomach LPL, duodenal LPL, jejunal LPL, ileal LPL, cecal LPL, colonic LPL). PCA display of Treg phenotype in each organ, with vectors indicating main phenotypic drivers of variance. **E)** UMAP representation with FlowSOM cluster overlay of high parameter flow cytometry data on Tregs grouped into four main source categories. **F)** Bar chart of FlowSOM cluster frequency in each category. **G)** Flow cytometry histograms showing key features of concatenated data from each Treg category. **H)** Flow cytometry histograms of Nur77 expression with naïve, activated and CD69^+^ Treg from the major tissue classes, with **I)** quantification across multiple tissues.

We next compared the phenotype of Tregs across each of 48 source tissues using a high dimensional flow cytometry panel allowing gating of Tregs and assessment of markers associated with Treg activation and residency: CD62L, CD44, CD25, Neuropilin-1, CD69, KLRG1, ST2, CD103, GITR, Tim-3, CCR2, CCR3, CCR6, CCR9, CXCR3, TIGIT, Ly-6c, CD95, TNFRII, ICOS and PD-1 (**Supplementary Figure 1**). When Treg phenotypes were compared between tissues, strong similarities were observed, with phenotypes aligning along lymphoid, non-lymphoid and gut-associated axes (**Figure 1D**). Grouping individual tissues into these larger tissue types provided resolution of Treg phenotypes into clusters (**Figure 1E**), where the blood Treg population was dominated by naïve and activated Tregs, lymphoid Tregs were largely naïve, activated and CD69^+^ Tregs, non-lymphoid Tregs were populated by activated and CD69^+^ Tregs, including enriched ST2^+^ and KLRG1^+^ populations, and gut-associated Tregs were populated by CD69^+^ CD103^+^ Nrp1^-^ Tregs (**Figure 1F**). None of these markers or populations were unique to a particular tissue type, with each tissue class including a diverse mixture of phenotypes, with only the relative intensity of markers (**Figure 1G**) and cluster frequency (**Figure 1F**) changing, the latter without distinct separation between the clusters. Shared across tissue classes, Tregs expressed the Nur77 reporter of TCR engagement at a high rate (**Figure 1H,I**), consistent with a high frequency of local antigen stimulation for Tregs across the different settings of lymphoid, non-lymphoid and gut-associated tissues, while trafficking between the sites in the blood occurred in the absence of TCR engagement. Together, these results demonstrate that Tregs are found in nearly all tissues, with a high phenotypic similarity spanning the populations found across non-lymphoid non-gut tissues.

A key requirement for comparative tissue phenotyping is the ability to discriminate between cells dwelling in the tissue and those captured due to vascular contamination. Here we used perfusion followed by anti-CD45 antibody vascular labelling to identify and gate out the vascular Treg population, which can be substantial in some non-lymphoid tissues (in particular liver and lungs; **Supplementary Figure 2A**). While the residual (post-perfusion) i.v. anti-CD45^+^ cells have previously been interpreted as blood contamination, and are here gated out of the tissue Treg population, the residual vascular component of Tregs in these tissue preparations was, however, phenotypically distinct from blood Tregs. For example, the i.v. CD45^hi^ Treg population in the liver included a CCR9^+^Nrp1^-^ population, normally absent from the blood but common in the liver tissue (**Supplementary Figure 2B**). Across the tissue groups, the residual vascular component was phenotypically closer to the i.v. CD45^-^ tissue Treg compartment than it was to the blood (**Supplementary Figure 2C,D**). Further, an i.v. CD45^int^ Treg population was observed, with phenotypes intermediate between the residual vascular (i.v. CD45^+^) and tissue-based (i.v. CD45^-^) populations, consistent with a perivascular or trans-vascular Treg population with partial vascular shielding (**Supplementary Figure 2B-D**). This result was unlikely to be driven by isolation effects, as the optimized tissue preparation approach used changed yield but not phenotype (**Supplementary Figure 3**). The residual vascular and peri/trans-vascular Treg populations are therefore more likely to represent cells in the process of in situ differentiation than random vascular contamination.

### Tissue Tregs are shaped by age and microbiome

We next sought to determine the degree to which sex, age and microbiome influence the tissue Treg niche and phenotype. For both tissue Treg frequency and absolute number, there were few differences between male and female mice, other than increased numbers of Tregs in the salivary glands of female mice and a trend towards increased Tregs in the white adipose tissue of male mice (**Supplementary Figure 4A,B**). At a phenotypic level, tissue Tregs in the non-lymphoid tissues were phenotypically indistinguishable between male and female mice in most organs, with the strongest differences being observed in the white adipose tissue and salivary glands, where the residency profile was intensified in male mice (**Supplementary Figure 4C,D**). This is consistent with the finding by Vasanthakumar et al, where white adipose Tregs show sexual dimorphism ^27^, but demonstrates that this is not a general property of tissue Tregs, as it is not replicated elsewhere, other than the salivary gland.

To assess the effect of age, we typed the tissue Treg population from mice aged 8, 12, 30, 52 and 100 weeks of age. Compared to the blood and lymphoid organs, where Treg number had a minor increase with age, tissue Tregs increased 5-20-fold across the non-lymphoid organs (**Figure 2A**), with the largest increases observed in white adipose tissue and liver (**Supplementary Figure 5A**). The largest average increase was in gut-associated tissues, although this was driven almost exclusively by a ∼100-fold expansion in the LPL Treg population at 2 years of age (**Supplementary Figure 5A**). In lymphoid tissues, the increase in Treg was associated with increased frequency of Tregs within the CD4 population (**Supplementary Figure 5B**), while in the non-lymphoid tissues the frequency was largely stable with age, with the exception of muscle and white adipose tissue, indicating that the large fold-increase was driven by an (inflammatory) expansion in the total CD4 T cell niche (**Supplementary Figure 5B**). At a phenotypic level, across the lymphoid and non-lymphoid tissues, phenotypic adaption intensified with age (**Supplementary Figure 5C**), with relative enrichment of the CD69^+^ gut-resident populations over time (**Figure 2B-D**). This result is consistent with both models of gradual accumulation of residential Tregs and with age-dependent expansion of the cellular niche for residential Tregs within the tissue.

**Figure 2.**
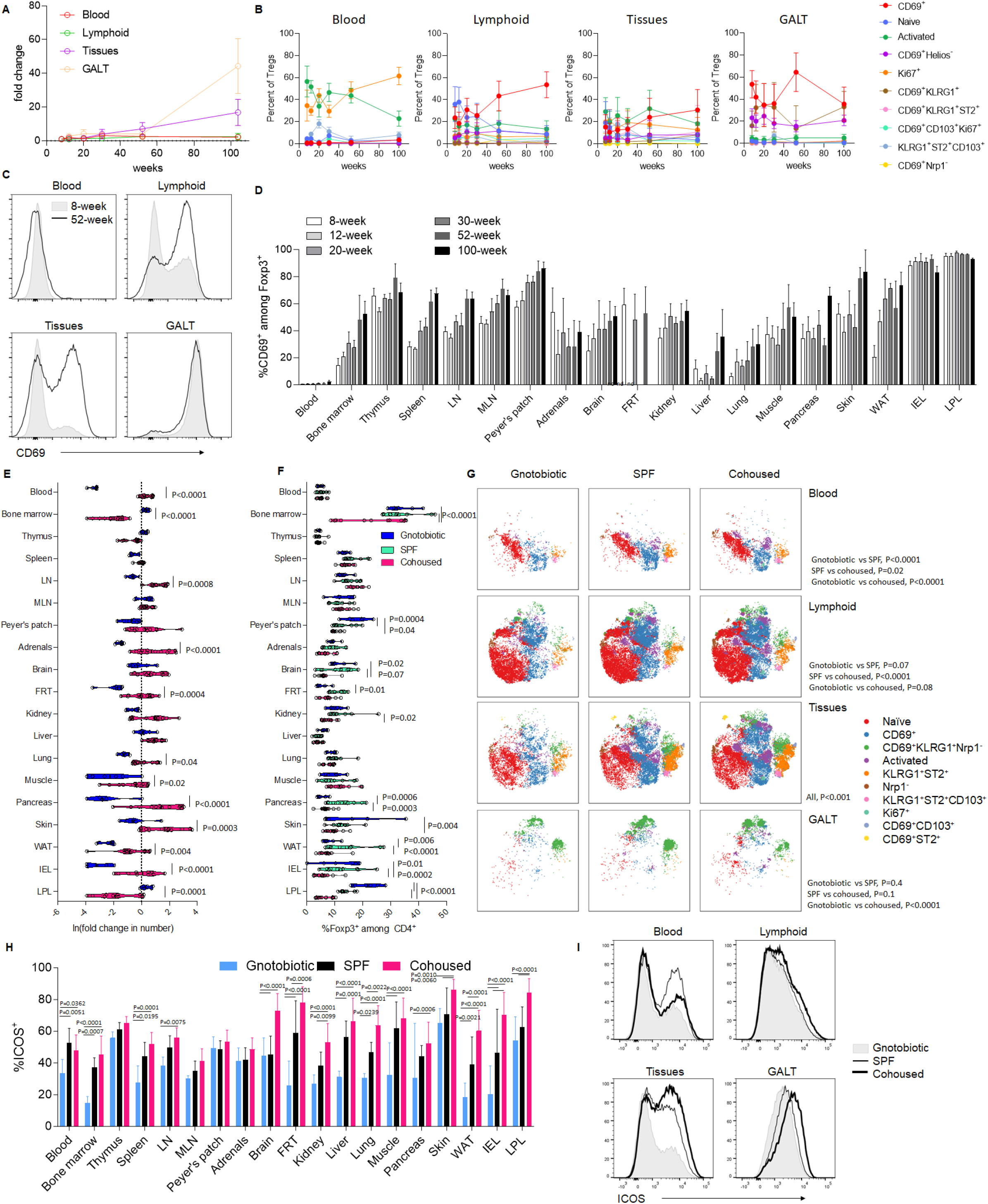
The tissue-resident Treg niche is shaped by aging and microbiome. Mice were perfused and assessed by flow cytometry for tissue Treg number and phenotype at 8, 12, 20, 30, 52 and 100 weeks of age (n=5-12. **A)** Fold change in Treg cell number in the blood, lymphoid tissues (thymus, spleen, LN, bone-marrow, mLN, Peyer’s patches), non-lymphoid tissues (skin, muscle, female reproductive tract, lungs, pancreas, brain, white adipose tissue, liver, kidney, adrenals) and gut-associated tissues (IEL and LPL). **B)** Quantitative change in phenotypic clusters of tissue Tregs by tissue class and age, based on annotated tSNE clusters. **C)** Histogram of CD69 expression at 8 and 100 weeks of age in the blood, lymphoid tissues, non-lymphoid tissues and gut-associated tissues. **D)** CD69 expression over age and tissue, within the Treg population. **E)** SPF-housed mice, gnotobiotic (germ-free) mice and wild-exposed co-housed mice (n=9, 6, 12) assessed by flow cytometry for tissue Treg number and phenotype at 8-12 weeks of age. Fold-change in tissue Treg number per tissue, and **F)** frequency of Tregs within CD4 T cell population per tissue. **G)** Phenotype of tissue Tregs in blood, lymphoid tissues (thymus, spleen, LN, bone-marrow, mLN, Peyer’s patches), non-lymphoid tissues (skin, muscle, female reproductive tract, lungs, pancreas, brain, white adipose tissue, liver, kidney, adrenals) and gut-associated tissues (IEL and LPL), displayed as tSNE of flow cytometry markers. **H)** ICOS expression over microbial exposure and tissues, within the Treg population, with **I)** amalgamated histograms in tissue classes.

Finally, we measured the impact of the microbiome on tissue Tregs, by comparing the standard SPF-housed mice to both gnotobiotic (germ-free) and microbial “re-wilded” mice. While effect sizes were lower than observed in CD8 T cells ^28^, a moderate increase in Treg number was associated with increased microbial complexity across the body, with the exception of the bone-marrow and the LPL population of the gut, where the opposite occurred (**Figure 2E**). In particular, the Treg numbers in the blood, pancreas, muscle, white adipose tissue and IEL compartment of the gut were reduced in size in the absence of a microbiome, while rewilding the microbiome substantially increased the population in the pancreas, skin, kidney and brain (**Figure 2E**). These effects were largely driven by changes in the total CD4 T cell infiltrate, as the frequency of Tregs within the CD4 T cell compartment was stable with microbiome changes. The exception to this was the bone-marrow and LPL compartments, where both the frequency of Tregs within CD4 T cells and the absolute number of Tregs fell markedly in re-wilded mice (**Figure 2E,F**), potentially reflecting displacement by effector CD8 T cells. Unlike that of CD8 T cells, where phenotypes became more residential with rewilding ^28^, Treg phenotype was relatively stable with the changing microbial complexity (**Figure 2G**, **Supplementary Figure 5D**), with the phenotypic cluster differences being driven by changes in ICOS expression, which was elevated with increased microbial complexity across most tissues (**Figure 2H,I**). Together, these results suggest that the tissue Treg niche is numerically expanded, but phenotypically preserved, by both age and microbial challenge.

### Overlapping molecular determinants of tissue Treg residency across the non-lymphoid tissues

To take a detailed unbiased analysis of the tissue Treg transcriptome, we performed a high depth bulk transcriptomics analysis of purified Tregs from the blood, lymphoid tissues (spleen, lymph nodes, Peyer’s Patches), gut tissues (IEL and LPL) and a representative selection of non-lymphoid non-gut tissues (liver, lung, pancreas, kidney). Tissue-residency and Treg purity was achieved by the use of extensive perfusion, on the *Foxp3^Thy1.1^*mice, allowing sort purification of Foxp3^Thy1.1+^ Tregs and the control Tconv group. An interactive analysis resource based on this data, providing cross-tissue comparisons and pathway analysis, is available at https://www.bioinformatics.babraham.ac.uk/shiny/expressionViewer/. Global analysis of the Treg populations found similar transcriptional profiles for the gut tissue and non-lymphoid tissue Treg populations (**Figure 3A**). Using pairwise comparisons, the Treg populations purified from non-lymphoid tissues (liver, lung, pancreas, kidney) demonstrated highly similar transcriptional profiles, with ∼95% transcriptional correlation (**Figure 3B**). IEL and LPL Tregs showed similar transcriptional similarity to each other. The largest differences observed, between the blood Treg and LPL Treg, still demonstrated >85% transcriptional correlation (**Figure 3B**). The small transcriptional differences observed in pairwise comparisons of tissue Tregs were largely non-significant, with only blood Tregs and gut tissue Tregs showing substantial numbers of significant transcriptional differences (**Figure 3C**). The largest set of transcriptional differences were genes downregulated in blood Tregs, a gene set which drove separation between the lymphoid, gut and non-lymphoid tissues (**Figure 3D**). Blood Tregs expressed lower levels of multiple migration-associated genes and effector cytokines (**Figure 3E**). Fewer transcriptional changes were observed when comparing other tissue classes (**Figure 3E**), with the strongest differences clustered within cytokine-cytokine receptor interactions, extracellular matrix-receptor interaction and cell adhesion molecules (**Figure 3F**). Integrins and chemokine receptors were prominent among these changes (**Figure 3G**), although the only class of tissue Tregs with a unique signature were gut Tregs, with elevated *Ccr9* and *Ccr5* and lower *Itgb1* and *Sell*.

**Figure 3.**
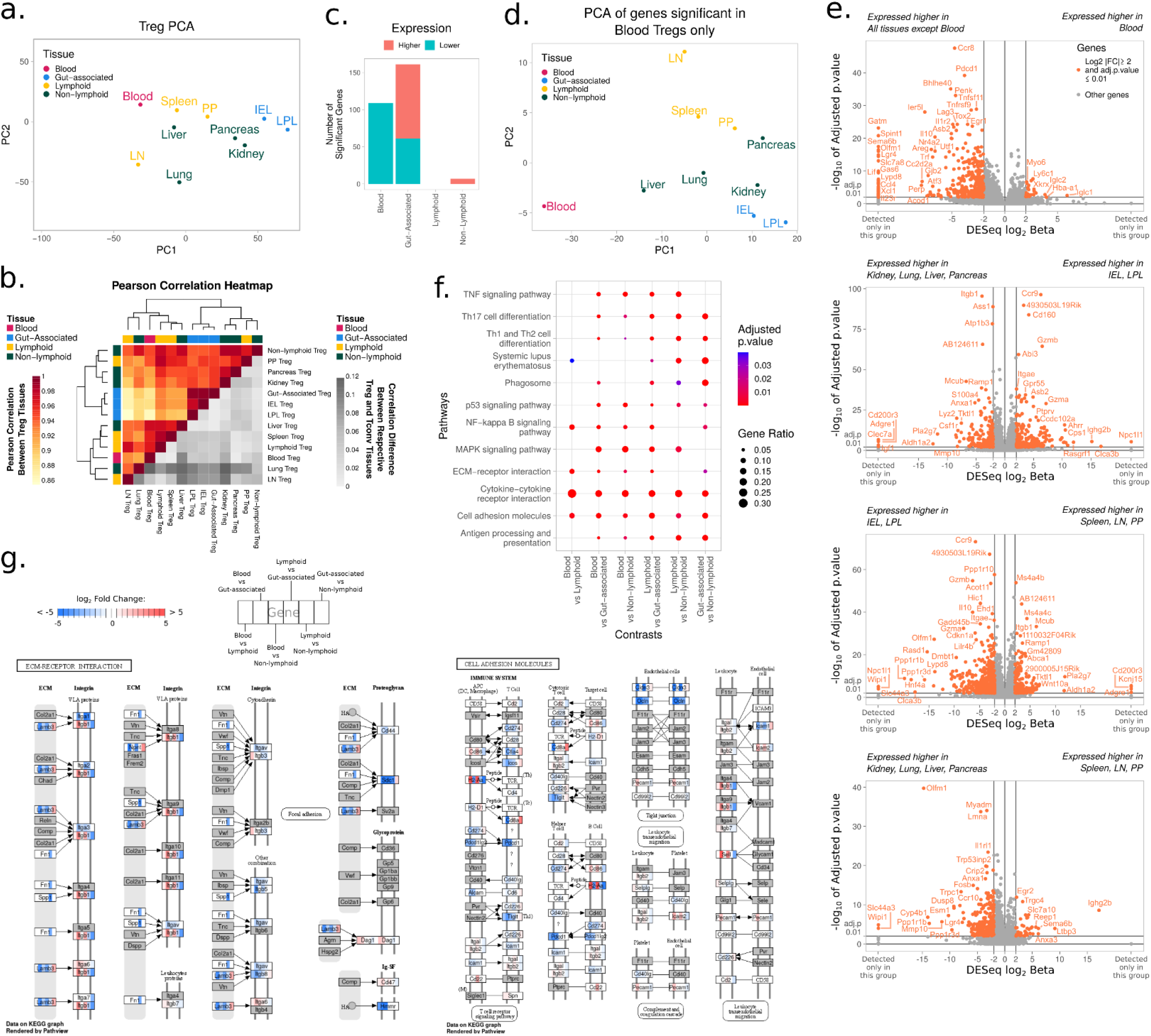
Transcriptional unity among tissue Tregs. Tregs were purified from the tissues of perfused *Foxp3^Thy1.1^*mice for RNAseq analysis. **A)** PCA of the Treg tissue mean expressions illustrating the broad clustering of the tissues in their respective tissue groups (n=3). **B)** Pearson correlation heatmap showing the similarity of the different Treg tissue mean expressions as well as the overall means of the four tissue groups. Additionally, the respective Pearson correlations in Tconv tissues and tissue groups are contrasted with the Treg correlations to show the similarity of the overall pattern of expression of genes. **C)** The number of genes persistently upregulated or downregulated in the respective tissue group when contrasted with each remaining tissue group pairwise. **D)** PCA of the Treg tissue mean expression using only the 109 genes persistently downregulated in blood Treg in panel C. **E)** Volcano plots of the differential expression analyses. Blood Tregs are contrasted with the means of all other Treg samples, followed by pairwise contrasts of the remaining three tissue groups. **F)** Dot plot showing the comparative overview of some of the various KEGG pathways enriched in the gene set enrichment analysis. **G)** The TNF Signalling pathway from KEGG overlaid with the log2 fold changes for genes found significant at adjusted p-value < 0.01 in the differential expression analysis showing the expression differences between the tissue groups for the coloured genes.

While tissue-resident Tregs were highly homologous across the different tissues, the generic tissue residency signature genes may still drive distinct functions in a tissue-specific manner. We therefore sought to functionally test the canonical tissue residency proteins of CD69, ST2, KLRG1, CD103, CD11a, amphiregulin (Areg), S1PR2, Hif1α, Blimp1 and BATF. While the relative expression of each of these signature proteins varies slightly from tissue to tissue, each of these proteins share a common expression profile of being low in the blood and lymphoid tissues, and upregulated across a broad set of non-lymphoid non-gut tissues, at both the protein (**Figure 4A**) and RNA (**Figure 4B**) level. We assessed mixed bone-marrow chimeras with 50%:50% wildtype and knockout bone-marrow reconstitution for *Cd69* KO (**Supplementary Figure 6**), *Il1rl1* (ST2) KO mice (**Supplementary Figure 7**), *Itgal* (CD11a) KO (**Supplementary Figure 8**), KLRG1 KO mice (**Supplementary Figure 9**), and CD4-Cre *Prdm1*(Blimp1)^flox^ mice (**Supplementary Figure 10**), *S1pr2* KO (**Supplementary Figure 11**), *Batf* Ko (**Supplementary Figure 12**), and *Hif1a* (HIF1α) KO mice (**Supplementary Figure 13**). We also generated and assessed *Areg* KO mice (**Supplementary Figure 14**) and *Itgae* (CD103) KO mice (**Supplementary Figure 15**). We used a summary matrix to assess the change in tissue Treg frequency across each of 15 different tissues in each of these 10 knockout mouse strains, considering both Treg as a proportion of CD4 T cells (**Figure 4C**) and the absolute number of Tregs within each tissue (**Figure 4D**). Despite their signature status, we found most of these residency genes had little impact on Treg number – ST2, CD103, S1PR2 and Hif1α knockouts had normal numbers of Tregs across all tissues surveyed. CD69-deficiency resulted in a mild decrease in Treg numbers across most tissues (**Supplementary Figure 6**). KLRG1-deficiency and BATF-deficiency drove an elevation and a reduction, respectively, in the number of Tregs across most non-lymphoid tissues (both gut and non-gut) (**Supplementary Figure 9** and **Supplementary Figure 12**), CD11a-deficiency reduced Tregs entering all tissues surveyed (**Supplementary Figure 8**), and Blimp1-deficiency enhanced tissue Treg numbers in the gut tissues (**Supplementary Figure 10**). No unique gene-tissue associations were observed to control tissue Treg numbers, other than an increase in Tregs in the skin of Areg-deficient mice, which could be due to local inflammation from Treg-extrinsic functions (**Supplementary Figure 8**), and an increase of S1PR2-deficient Tregs in the kidney (**Supplementary Figure 11**). As the CD69 KO result was partially confounded by reduced output of CD69 KO Tregs from the thymus (**Supplementary Figure 6**), we generated *CD69^flox^* mice to breed and assess *Foxp3^CreERT2^ CD69^flox^* mice. Following tamoxifen addition, Tregs from these mice were equally competitive at populating the tissues (**Supplementary Figure 16**). We further assessed the effect of genetic deficiency on Treg phenotype in each KO (**Supplementary Figures 6-16**). Using cross-entropy as a measure of global phenotypic difference ^29^, as measured in each tissue-resident population, only minor shifts were observed in Treg phenotype across the matrix of tissue types and genetic deficiencies (**Figure 4E**), with the exception of CD4-Cre Blimp1^flox^ mice demonstrating loss of the Helios+ Treg population within IEL and loss of the CD69^+^KLRG1^+^CD103^+^ Treg population within the skin (**Supplementary Figure 10**). Thus, while tissue-resident Tregs upregulate this signature gene set, expression of each member is largely independent of the other members. Together, these results suggest that not only do tissue Tregs show a unified phenotype across the non-lymphoid non-gut tissues, but they also appear to share the same key molecular determinants for residency.

**Figure 4.**
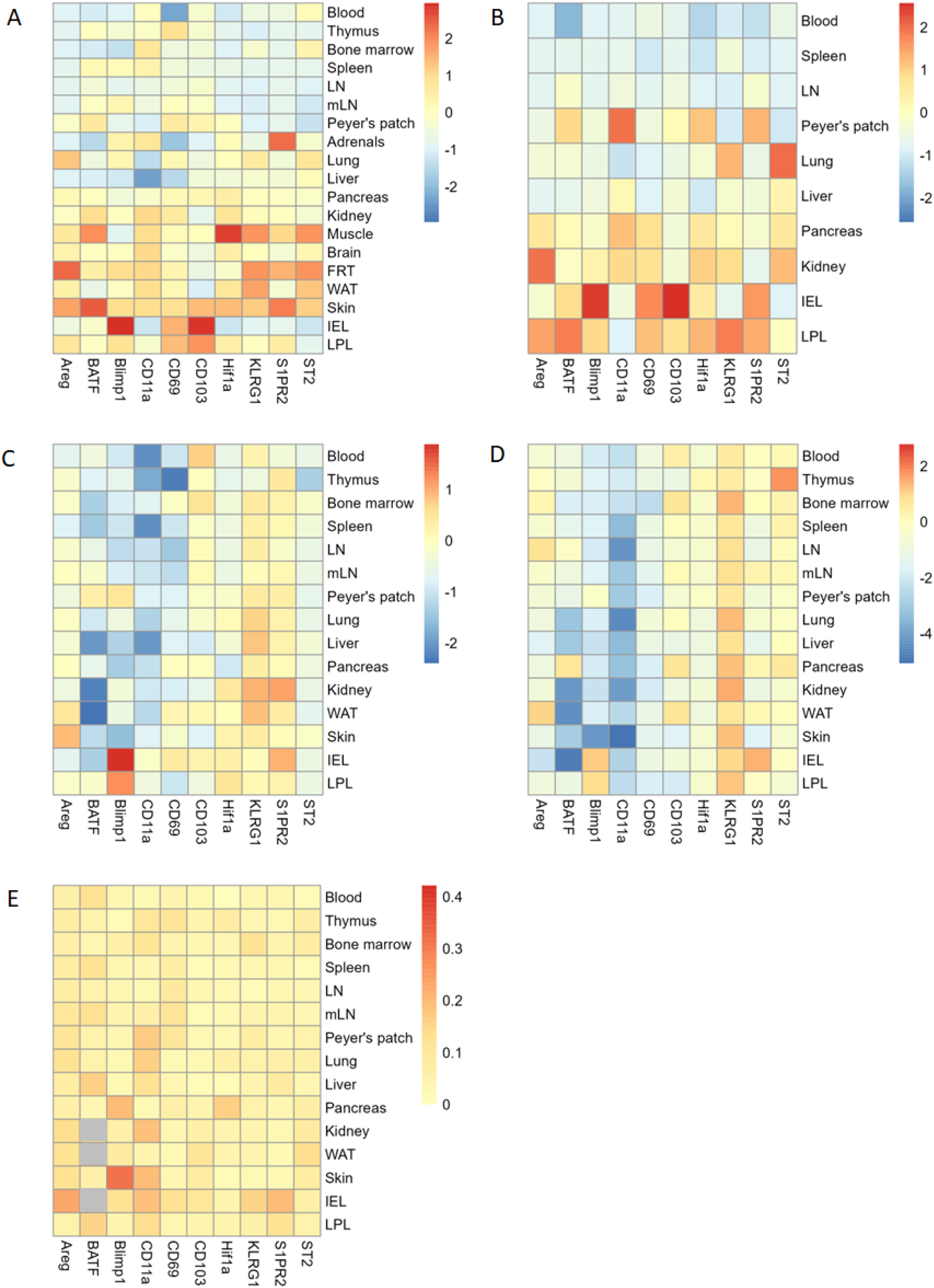
Conserved molecular determinants for Treg tissue residency across different tissues. **A)** Relative expression of Areg, BATF, Blimp1, CD11a, CD69, CD103, HIF1a, KLRG1, S1PR2 and ST2, as measured by flow cytometry, within tissue-resident Tregs from the blood, thymus, bone marrow, spleen, lymph nodes, mesenteric lymph nodes, Peyer’s patches, adrenals, lung, liver, pancreas, kidney, muscle, brain, female reproductive tract (FRT), white adipose tissue (WAT), skin, intraepithelial lymphocytes (IEL) of the gut and lamina propria lymphocytes (LPLs) of the gut (n=4). **B)** Relative mRNA expression of *Areg*, *Batf*, *Prdm1*, *Itgal*, *Cd69*, *Itgae*, *Hif1a*, *Klrg1*, *S1pr2* and *Il1rl1*, as measured by RNAseq, within tissue-resident Tregs from the blood, spleen, lymph nodes, Peyer’s patches, lung, liver, pancreas, kidney, intraepithelial lymphocytes (IEL) of the gut and lamina propria lymphocytes (LPLs) of the gut (n=3). **C)** In separate experiments, the number and phenotype of tissue Tregs was assessed in mixed bone-marrow chimeras with 50%:50% KO:wt bone-marrow, containing bone-marrow from BATF, Blimp1, CD11a, CD69, Hif1a, KLRG1, S1PR2 or ST2 KO bone-marrow (n=5, 6, 6, 4, 5, 4, 6, 4) or from comparison of wt and KO mice for Areg and CD103 (n=4, 4). Treg frequency among CD4 T cells and **D)** absolute number in the blood, thymus, bone marrow, spleen, LN, mLN, Peyer’s patches, lung, liver, pancreas, kidney, white adipose tissue, skin, IEL and LPL in KO relative to wt. **E)** Treg phenotype change, based on tSNE cross entropy distance between KO and wt cells from the same tissue. Gray indicates insufficient events for phenotypic assessment.

### Tissue Tregs are short-term residents in non-lymphoid tissues

The similarity in transcriptional programs and molecular dependency exhibited by tissue-resident Tregs across multiple different non-lymphoid tissues suggests that either the phenotypic plasticity of Tregs is sharply curtailed, preventing niche-driven specialization, or that the dwell time for residency is of a limited duration, limiting divergence. To directly test the homeostatic population flows for tissue Tregs we performed a large multi-timepoint parabiotic experiment, consisting of CD45.1 mice parabiosed to CD45.2 mice, providing a common circulatory system, and assessed at 2, 4, 8 and 12 weeks post-surgery for displacement of tissue-resident cells with circulatory cells. At each timepoint, the host/donor composition was assessed in the blood, lymphoid tissues (bone-marrow, spleen, lymph nodes, Peyer’s patches and mesenteric lymph nodes), non-lymphoid tissues (adrenals, brain, kidney, lung, liver, white adipose tissue and skin) and the gut-associated tissues (intraepithelial, lamina propria) by high dimensional flow cytometry. The use of multiple time-points in this experimental design allows for the estimate of dwell times, rather than a simple assignation of residency at a single time-point. Using the division of tissue Tregs into the states of resting CD44^-^CD62L^hi^ cells, activated CD44^+^CD62L^low^ cells and CD69^+^ cells, the CD69^+^ population showed the slowest displacement kinetics, consistent with the “residential” status of these cells (**Figure 5A**). To calculate the residency dwell time of the tissue Tregs, we developed a probabilistic Markov chain model of the cellular kinetics using Bayesian analysis, where each individual tissue was modelled in an interconnected population set consisting of the tissue, the blood, and a combined pool of all other tissues, with the population sets of the parabiotic pairs being linked only through the blood (**Supplementary Figure 17**). A feature of the probabilistic Markov approach is the ability to predict transition rates of best fit to the observed data, with the primary differences across tissues being the modelling entry rate (**Figure 5B-D**). The best fit Markov models exhibited large differences in entry rates, with the highest rates modelled from lymphoid tissue data, being around 100-fold higher than the entry rates into the gut tissues, which were in turn 10-fold higher than the entry rate into the average non-lymphoid non-gut tissues. There was, however, diversity within the non-lymphoid tissue models, with some tissues (liver, pancreas) modelled to have entry rates equivalent to gut tissues, while other tissues (adrenals, brain) were 10-fold lower than the average non-lymphoid tissue (**Figure 5B-D**). Tissue entry rates modelled for CD69^+^ Tregs (**Figure 5D**) were ∼10-fold higher than those modelled for resting or activated Tregs (**Figure 5B,C**), across all tissue types. The aggregate models, fitting the empirical data collected (**Supplementary Figure 17**), demonstrate limited residency for Tregs across all tissues. Modelled dwell-times for resting Tregs and activated Tregs were short across all tested tissues, on the order of ∼5 days, with most of the residency exhibited by the CD69^+^ population (**Figure 5E**). Within lymphoid tissues, even the residency of the CD69^+^ fraction was short: 3 days for lymph nodes, 17 days for spleen,21 days for bone-marrow, and 4 weeks for Peyer’s Patches and mesenteric lymph nodes (**Figure 5E**). The average dwell time for CD69^+^ Tregs within non-lymphoid tissues was slightly longer, at 23 days on average, with most tissues showing tissue-residency dwell times of around 3 weeks, with white adipose tissue extending out to 8 weeks (**Figure 5E**). Gut-tissue CD69^+^ Treg dwell time was at the upper end of this range, with 7-8 weeks for IELs and LPLs (**Figure 5E**). These dwell times were probabilistic, with individual cells modelled to fall along a distribution centered on these values (**Supplementary Figure 18**). Building a cellular population model featuring the blood, median lymphoid tissue and median non-lymphoid tissues (**Figure 5F**), lymphoid tissue kinetics are driven by the greatly elevated entry rates, with non-lymphoid tissues being seeded through direct entry of CD69^+^ Tregs from the blood, and by indirect seeding through highly transient resting/activated Tregs, of which a small fraction upregulate CD69 (**Figure 5F**). After the median dwell time of 3 weeks, the majority of these CD69^+^ Tregs are modelled to directly leave the tissue, with smaller numbers dedifferentiating or dying (**Figure 5F**). Consistent with this model of reversible residency, *KLRG1^Cre^ RosaA14* mice, with fate-mapped KLRG1 Tregs, have substantial populations of ex-KLRG1-expressing Tregs across the tissues, with activated but not naïve/resting phenotypes (**Supplementary Figure 19**). Together this data formally demonstrates a short dwell time for tissue-resident Tregs, across the full set of tissues analyzed, and supports a circular migration model, where activated Tregs enter, differentiate, transiently reside and leave the tissues.

**Figure 5.**
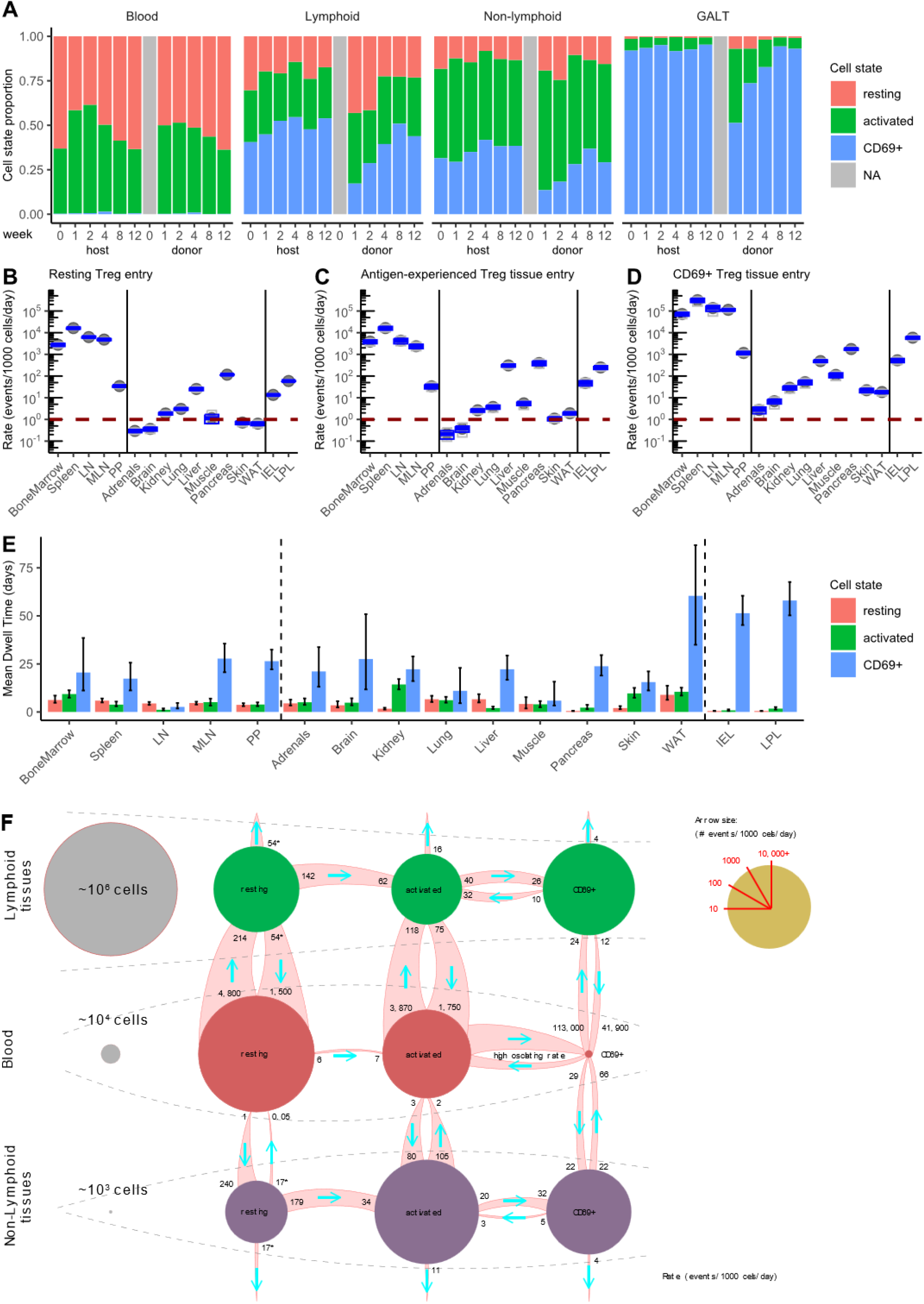
Tregs rapidly adopt a residency phenotype for limited duration tissue dwell time. CD45.1 mice were parabiosed to CD45.2 mice. Pairs of parabiotic animals were perfused and sacrificed at weeks 1, 2, 4, 8, and 12 by flow cytometry across the indicated tissues (n=11,12,18,16,14). **A)** Frequency of resting, activated and resident Tregs within the blood and tissues, for host cells and donor cells. **B)** Markov chain results for entry rates for resting Tregs, **C)** activated Tregs, and **D)** CD69^+^ Tregs. **E)** Markov chain modelling was used on the parabiotic data to estimate Treg dwell times among the CD44^-^CD62L^hi^, CD44^+^CD62L^low^ and CD69^+^ Treg subsets, for blood, lymphoid (bone marrow, spleen, LN, mLN, PP), non-lymphoid (adrenals, brain, kidney, lung, liver, WAT, skin) tissues and gut-associated (IEL, LPL) tissues. Model estimates with 80% credibility interval. **F)** Based on the Markov chain models for Tregs, the primary population flows between blood and tissues, indicating population size and the frequency of key events. * For tissue-dwelling resting Tregs the model does not distinguish between cell death and return to the blood.

### A common pool of tissue Treg cells seed multiple distinct non-lymphoid tissues

Our data demonstrates that tissue Tregs, across a large range of non-lymphoid tissues, initiate a common molecular program upon seeding tissues, with a highly transient residency. While incompatible with a long-term residency model, these data are compatible with two distinct models: clonally-restricted circular trafficking vs pan-tissue trafficking. Under the first model, Tregs from each tissue would show circular migration between one non-lymphoid tissue and the lymphoid tissues, providing for clonal-restriction while giving short dwell times. Under the second model, Tregs would show the capacity to seed multiple non-lymphoid tissues, with circular flow resulting in the distribution of tissue Treg clones across multiple tissues. To distinguish between these two models, we assessed both the clonality of tissue Tregs in a subset of non-lymphoid tissues, and the capacity for tissue Tregs to cross-seed alternative tissues.

We first investigated clonal sharing between non-lymphoid tissues, using the recombined TCR as a clonal barcode. We performed single cell sequencing of purified tissue-resident Tregs from four independent mice, focusing on the blood, as a reference population, and kidney, pancreas, liver and LPL as representative non-lymphoid tissue populations. TCRβ chain clones were used as markers of clonality, with the protein-level sequence used to enable cross-mouse identification. Despite being unmanipulated mice, with fully polyclonal repertoires, tissue-resident Tregs showed a high degree of clonal sharing within mice, suggestive of the presence of widely distributed clones within the tissue-resident populations (**Figure 6A**). TCR clones from Tregs purified from the liver, pancreas and kidney were highly represented within the blood and alternative non-gut tissues tested (**Figure 6A**). By contrast, the TCR clones identified in Tregs purified from the LPL population were only rarely observed in the non-gut tissues, despite high levels of sharing within mice (**Figure 6A-C**). To quantify the relative degree of sharing between tissues we assessed the frequency of shared clones between tissue pairs, using an extracted half-sample as the comparator. Between liver, pancreas and kidney comparisons, an average of 7.5% of TCR clones were shared, approaching the average of 12.6% sharing between half-samples (**Figure 6B**). By contrast, LPL Tregs only shared, on average, 1.3% of TCR clones, well below the half sample reference of 6.7% (**Figure 6B**). This data suggests that while the gut Treg population may largely follow the clonally-restricted circular trafficking model, the preponderance of clones in the non-gut non-lymphoid tissues are of a pan-tissue clonality.

**Figure 6.**
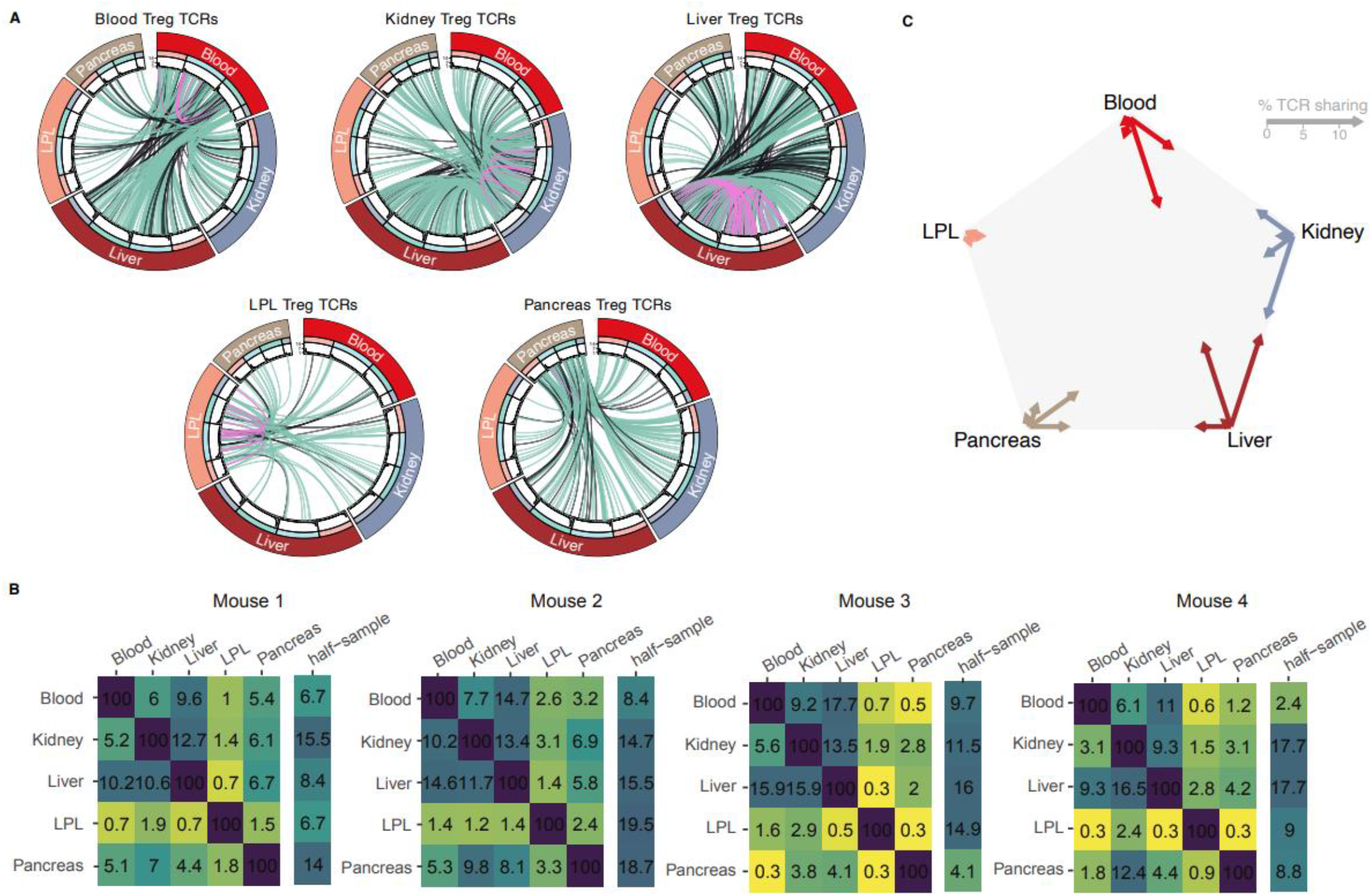
Extensive clonal sharing among Tregs from non-lymphoid organs. *Foxp3^Thy1.1^* mice at 16 weeks of age were injected with intravenous anti-CD45 antibody label prior to FACS sorting of Tregs from blood, kidney, liver, pancreas and LPL (n=4) for analysis by scTCRseq. **A)** Chord diagrams illustrating for each tissue origin the Treg TCR found to be shared between different tissues. For each tissue (main sector) of each mouse (sub-sector), the frequency ranked TCR repertoire is shown as a bar plot. Lines joining 2 repertoires indicate a clonotype sharing between 2 tissues of the same mouse (green), the same tissue type between different mice (pink) or different tissue types in different mice (black) **B)** Heat maps showing for each mouse the percentage of cells from x-tissue having a TCR also observed in y-tissue. Since sharing between small diverse repertoires is not expected to result in high sharing percentage even if the same sample would be compared to itself, for each x-tissue, the sharing half-sample subpool compared to the other half of x-tissue sample in indicated on the right (average value of 100 sub pools). **C)** Percentage of TCR sharing between tissue within each mouse, averaged for the 4 analysed mice, illustrated the length of an arrow pointing toward the tissue, where the distance between tissues represents 100% sharing.

The strong sharing of TCR clones among tissue Tregs across multiple tissues, in combination with our data indicating that tissue Tregs have TCR engagement within the tissues, is suggestive of tissue Treg clones recognizing pan-tissue antigens. While tissues are usually conceptualized by the distinct or unique expression patterns of constituent cells, this is not reflected in the available self-antigen pool. Indeed, reanalysis of a multi-tissue MHCI immunopeptidome dataset ^30^ suggests that only 1.4% of peptides are tissue-restricted (**Supplementary Figure 20A**), and, furthermore, the tissue-restricted peptides are generally found in lower abundance than the shared peptides (**Supplementary Figure 20B**). While Tregs recognize antigen presented on MHCII, this data demonstrates that the tissue peptidome available for presentation is heavily overlapping between tissues. Thus, unless an as-yet-to-be-determined mechanism is available to actively skew Treg recognition away from shared tissue antigens, an antigen-agnostic model would predict the majority of tissue Tregs to be reactive against shared antigens, consistent with our TCR clonotype data.

To formally test the functional outcome conferred upon tissue Tregs by these clonotypic TCRs, we sought to biologically challenge the identified pan-tissue Treg TCRs via a retrogenic system. In order to multiplex the retrogenic assay, and create a more representative perspective on tissue Treg TCRs, we adapted the Pro-Code epitope-based barcoding system ^31,32^, which enables cell barcode detection at the protein level. The original Pro-Code system utilised a triplet combination of 14 linear epitopes, to create 364 unique Pro-Codes. When incorporated into the retrogenic vector, the membrane-bound Pro-Codes were incompatible with *in vivo* lymphocyte-tracing, so we generated a Pro-Code variant, Flow Cytometry-optimised Histone-fused Pro-Codes (named here FlowCodes), with enhanced detection capacity by flow cytometry. This was achieved by cloning the Pro-Codes epitopes into a fusion with Histone 2B in a retrovirus backbone (**Figure 7A**). The long half-life and chromatin localisation of H2B allows a fixation-resistant Pro-Code with sufficient sensitivity and lack of immunoreactivity, allowing the Pro-Codes to be detected *in vivo* in lymphocytes (**Figure 7B**). Into each of 21 Flow-Code retrovirus backbones we inserted a paired TCRα/TCRβ chain. Twenty of the pairs were cloned from tissue Tregs (**Figure 6**) and one was cloned from the OTII TCR. The co-expression of a retrogenic TCR with a unique FlowCode allows cells expressing a specific TCR to be tracked by flow cytometry. Retrogenic FlowCode retroviruses were individually used to transduce RagKO donor bone marrow stem cells, which were then pooled and mixed with WT bone marrow stem cells before injection into irradiated wildtype mice, to allow the differentiation of 21 different transgenic TCR progenitors in a polyclonal environment. The 20 tissue Treg TCRs were selected from among the most populous tissue Treg clonotypes, including 10 clones selected from the 20 most highly distributed Treg TCRs (**Supplementary Figure 21A**), and 10 clones selected based on the highest detection in individual tissues, without consideration of cross-tissue distribution (**Supplementary Figure 21B**). The dual selection approach created a TCR set representative of tissue Treg TCRs as a whole, with representation of the most frequently observed tissue Treg clonotypes (**Supplementary Figure 21C**), and a similar distribution of multi-tissue vs single-tissue detection to that observed in the public clone repertoire (**Supplementary Figure 21D**). Ten weeks after transplantation, differentiated CD4 T cells expressing the different TCRs were assessed for tissue representation and Treg fate by staining for the 7 Pro-Code epitopes along with different cell markers, analysing by flow cytometry, and deconvoluting using the custom automated gating pipeline FlowCodeDecoder.

**Figure 7.**
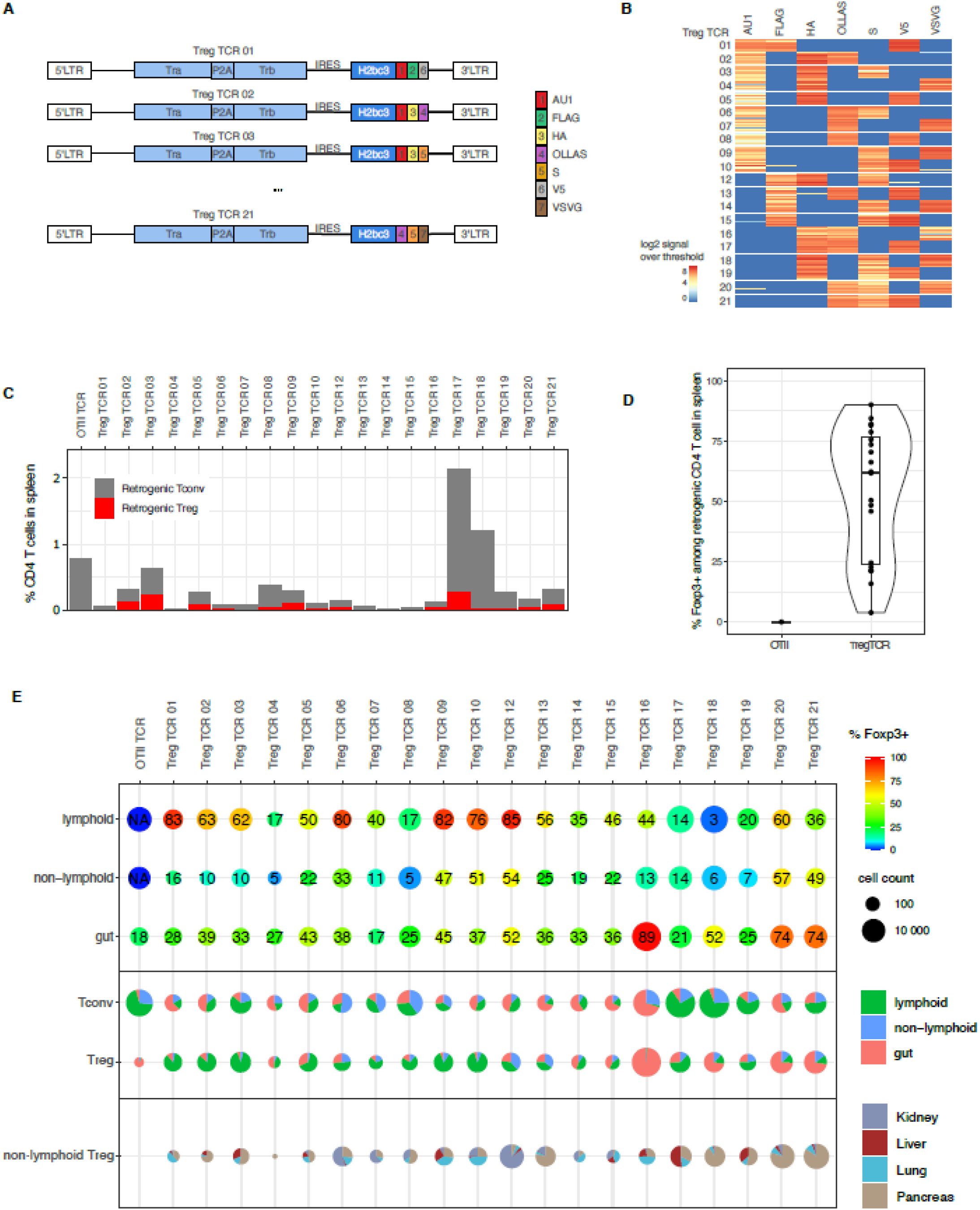
Multiplexed tissue Treg TCR retrogenics demonstrate pan-tissue functionality. 20 tissue Treg TCRs were selected from scSeq data, comprising 10 TCRs selected on the basis of distribution width and 10 TCRs selected on the basis of tissue frequency. **A.** The selected TCR panel, representative of the TCRs cloned from the tissue Treg population as a whole, were cloned into FlowCode retroviruses, combining the TCRα and TCRβ sequences with a unique triplicate ProCode epitope combination. **B.** Rag-deficient bone-marrow stem cells were individually transduced with the 20 retroviruses and pooled for reconstitution of irradiated mice. After 10 weeks, tissue samples were prepared for analysis by flow cytometry. FlowCodeDecoder was used to assign each T cell to the appropriate TCR clone, based on expression of the 7 epitopes. **C.** Quantification of Tconv (CD4^+^Foxp3^-^) and Treg (CD4^+^Foxp3^-^) T cells from each retrogenic TCR clone, based on splenic data, with **D.** average (and standard deviation) frequency of Treg fate within each clone (each data point corresponding to an individual retrogenic clone). **E.** Individual results from T cells derived from each of the FlowCode retrogenic TCRs, across the assessed tissues of spleen, LN, kidney, lung, liver, pancreas and LPLs. Top, absolute cell count of detected CD4 T cells for each clone (size) together with the frequency of Tregs within the population (indicate by colour, with percentage listed on each sample). Tissues groups are combinations of lymphoid (spleen, LN), non-lymphoid (kidney, lung, liver, pancreas) and gut (LPL) tissues. Middle, for Tconv (CD4^+^Foxp3^-^) and Treg (CD4^+^Foxp3^-^) T cells from each retrogenic TCR clone, a pie-chart visualising the distribution of cells within the lymphoid, non-lymphoid and gut compartments (pie chart size representative of clonal population). Bottom, for non-lymphoid tissue Tregs from each retrogenic TCR clone, a pie-chart visualising the distribution of cells within the non-lymphoid tissues of kidney, lung, liver, pancreas (pie chart size representative of clonal population).

While control retrogenic clones, bearing the OTII TCR only, remained conventional Foxp3^-^ cells, the retrogenic cells bearing each of the 20 tissue Treg TCRs spontaneously entered the Treg lineage, with Foxp3 expression detected at an average of >60% across the different clones (**Figure 7C,D**). Analysis of retrogenic tissue Treg clones at the tissue level revealed broad ability of the T cells to enter lymphoid, non-lymphoid and gut-associated tissues (**Figure 7E, Supplementary Figure 21E**). Of the ten retrogenics cloned from Tregs found in widely dispersed tissues, nine were found as tissue-Tregs in multiple non-lymphoid tissues, with the exception (TCR04) found only in the pancreas, but at the lowest frequency of all the clones making it a potential under-sampling detection issue (**Figure 7E, Supplementary Figure 21E**). Of the ten retrogenics cloned from Tregs based on high frequency detection within a single tissue, all were found as tissue-Tregs in multiple non-lymphoid tissues (**Figure 7E, Supplementary Figure 21E**). Five of these clones (TCR13/14/15/17/19) showed broadly balanced non-lymphoid tissue representation, indicating that the initial detection as tissue-restricted may have been driven by stochastic sampling. By contrast, 5 clones showed a strong, but not exclusive, tissue bias, aligning with the source of initial identification: TCR12 in the kidney, TCR20/TCR21 in the pancreas, and TCR16 and TCR18 in the LPL of the gut (**Figure 7E, Supplementary Figure 21E**). Together, these results formally demonstrate that a representative set of tissue Treg TCRs confer upon CD4 T cells a Treg fate, with the majority of TCRs driving a pan-tissue migration and residency profile and only a minority of TCRs driving a tissue-biased distribution.

As an independent test of the pan-tissue Treg model, we assessed the capacity of tissue Tregs to migrate into different organs. First, we returned to the parabiosis system. The equilibration of tissue Treg frequency following parabiosis (**Figure 5**) is consistent with both single-tissue or multi-tissue recirculation patterns. We posited, however, that under a single-tissue recirculation model, Tregs entering the system from a donor mouse that had never been exposed to a particular tissue would be at a competitive disadvantage in repopulation of that tissue. By contrast, under a multi-tissue recirculation model, the pan-tissue Treg population would show a similar capacity to populate a novel organ, even under competitive conditions. To create this novel organ competitive test, we compared the repopulation of the female reproductive tract in female:female parabionts (where the donor population would be experienced at female reproductive tract residency) to male:female parabionts (where the male donor population would be naïve for female reproductive tract residency). Comparing the frequency of host cell replacement by donor cells at 4 weeks post-parabiosis, at which point tissue Tregs were not completely normalized (i.e., with competitive replacement), the female reproductive tract was equally populated by donor cells from male mice (male:female parabionts) as by donor cells from female mice (female:female parabionts) (**Figure 8A**). The male donor and female donor Tregs were not only able to repopulate the female reproductive tract with equal efficiency, they were also phenotypically indistinguishable (**Figure 8B**).

**Figure 8.**
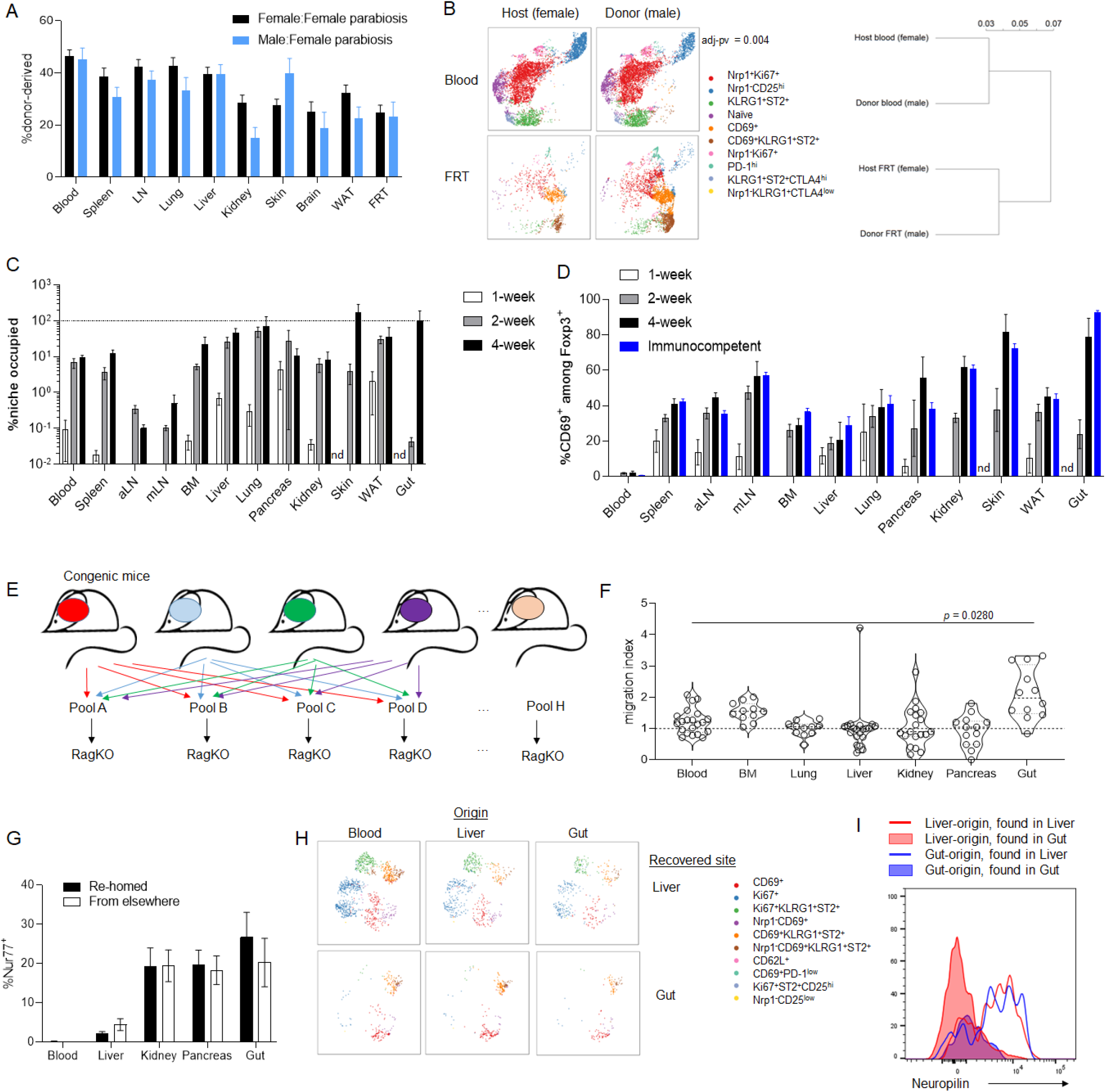
Tissue Tregs retain multi-tissue homing capacity and phenotypic plasticity. **A)** Proportion of incoming (donor-derived) Tregs in female mice when parabiotically paired to female (n=18) vs male (n=8) mice. All male:female comparisons for tissues resulted in p>0.05 by Sidak’s multiple comparison test. Mice were perfused prior to analysis. **B)** UMAP plots and dendrogram of high parameter flow phenotyping data on Tregs from male:female parabiotic pairs. No statistically significant difference between male and female cells in the female reproductive tract. **C)** Splenocytes were transferred into RagKO mice, with the Treg compartment measured quantitatively within each organ after 1, 2 or 4 weeks of reconstitution, in reference to unmanipulated numbers (n=6,6,3,9). Mice were injected with intravenous anti-CD45 label prior to sacrifice. **D)** Expression of CD69 on Tregs within each organ, 1, 2 or 4 weeks post-reconstitution, with comparison to unmanipulated mice. **E)** Eight pools of mixed tissue lymphocytes were generated from eight mouse strains with distinct markers. Each pool contained only one tissue from each mouse strain, creating a composite pool where the mixed tissue lymphocytes could all be traced for tissue origin based on the distinct markers of the donor strains. Pools were injected into RagKO mice. Mice were injected with intravenous anti-CD45 label prior to sacrifice. **F)** 4 weeks after the transfer of donor-tracked tissue pools, recipient mice were assessed for tissue Treg infiltration into blood, bone marrow, lung, liver, kidney, pancreas and gut (LPL). Rehoming migration index was calculated for each donor tissue type (n=7-11), where 1 was normalised to no preference for rehoming, and values above one indicate the fold-increase in preference for tissue of origin. Statistical testing by one-way ANOVA with Dunnett’s post-test. **G)** Expression of Nur77 in Tregs recovered post-transfer (n=6-10). All intra-tissue comparisons resulted in p>0.05 by Šídák’s multiple comparison test. **H)** tSNE plots showing phenotypic profiles of Tregs recovered from the liver and gut, sub-divided into cells coming from blood, liver or gut donors. **I)** Neuropilin histogram of liver and gut origin Tregs, recovered in the liver or gut.

Second, to assess the speed at which the tissue Treg compartment could be reconstituted, we transferred whole splenocytes into Rag-deficient mice. By measuring the quantitative (**Figure 8C**) and qualitative (**Figure 8D**) reconstitution of the compartment within different tissues, we observed that the tissue Treg population could be rapidly reconstituted from lymphoid Tregs. Within 2-4 weeks, the population of tissue Tregs were at 10-100% of the homeostatic level, across the non-lymphoid tissues (**Figure 8C**), and even at the 2-week time-point those Tregs that had reentered the tissues had upregulated CD69 to levels similar to that of homeostasis (**Figure 8D**). This data is consistent with a tissue Treg niche, with high capacity for rapid entry from the periphery.

Third, we performed a direct recirculation experiment. By extracting tissue-resident lymphocytes from 7 tissues from each of 8 antigenically-marked strains, we could make mixed ex-tissue-resident lymphocyte pools, where the tissue-resident Tregs extracted from each tissue had a unique antigenic marker combination (**Figure 8E**). These mixed pools were then injected into Rag-deficient hosts, and four weeks later the migration of tissue Tregs into the blood, bone-marrow, liver, lung, kidney, pancreas and gut was assessed by flow cytometry. A migration index was assessed for tissue-resident Tregs from each original donor organ, calculating the relative frequency by which the tissue Tregs were recovered from the same tissue as originally extracted from. While gut-purified Tregs were biased towards reentry into the gut, and bone-marrow Tregs showed a slight non-significant bias towards reentry into bone-marrow, Tregs originally isolated from the kidney, liver, lung and pancreas showed no preference for reentry into the tissues-of-origin (**Figure 8F**). Using Nur77 as a marker for TCR engagement, on a smaller scale repeat experiment, similar levels of antigen recognition were observed in the Tregs isolated from the different organs regardless of whether they had rehomed from the original source tissue or had cross-populated the tissue from a different source (**Figure 8G**). Comparing gut-associated Tregs to a non-lymphoid tissue such as the liver, providing for phenotypic differences (**Supplementary Figure 1**), the cross-populating Tregs exhibited a phenotype closer to that of the new host tissue than that of the originating host tissue (**Figure 8H,I**). Together, the parabiosis and tissue-transfer experiments strongly support a pan-tissue model for tissue Treg residency, whereby the same population of Tregs are capable of repopulating multiple tissues, with only slight preferences for target tissues other than the gut.

## Discussion

The shared phenotype of tissue Tregs and the evidence for a circular migration capacity leads to a parsimonious model of a pan-tissue Treg population. Under the pan-tissue model, activated Tregs have the capacity to enter, in a non-specialized manner, non-lymphoid tissues at a low rate. These activated Tregs would only be retained in the tissue for a short duration if their antigen was not present; if antigen recognition occurs, the stimulation would lead to retention, initiation of the generic tissue residency transcriptional program, and a dwell-time of weeks to months before death or egress. As most Tregs within the tissue pool would bear a TCR capable of recognizing antigen within multiple tissues, the aggregate effect of this model would involve the slow percolation of tissue Tregs, bearing shared TCR clones, throughout multiple tissues over the lifetime of the cell. We do not exclude the possibility of rare clones within a tissue pool with a more specialized circular pattern, where dwell time is largely restricted to a single tissue. Indeed, this model of antigen-mediated retention would necessarily lead to such migration patterns when the source antigen is tissue-restricted. This is consistent with the isolation of an adipose tissue TCR clone (clone 53), where TCR transgenesis drives high enrichment of Tregs within the adipose, although notably enrichment was also observed within muscle, and few other non-lymphoid tissues were assessed ^33^. However both theoretical considerations (the small fraction of peptides presented per organ that are restricted to a single tissue) and empirical data (agnostic rehoming capacity and shared TCR clones) necessitate such tissue-restricted clones being among the minority of tissue Tregs within the tissue. No *a priori* mechanism exists for how a Treg expressing a TCR with multi-tissue specificity could be differentially activated or retained within a tissue, when compared to a Treg expressing a TCR with mono-tissue specificity. Indeed, as Treg proliferation is driven by antigen-recognition, one would expect that any Tregs with a multi-tissue specificity would bear a competitive advantage over Tregs with a mono-tissue specificity and would, over time, grow to dominate the tissue repertoire. Through that prism it may be interesting to reflect on the Aire-dependent neonatal Tregs, where evidence for distinctiveness was higher than that observed here in tissue Tregs ^34^. During early development, rare clones with mono-tissue specificity could concentrate and expand within the tissue of antigen-recognition, while in later life the inexorable consequence of antigen-driven proliferation should dilute out these clones with multi-tissue specificity, especially when shared with lymphoid tissues.

The pan-tissue Treg model stands in contrast to the preceding seeding and specialization model. While additional data has driven a divergence of the model used in previous papers, the data of those prior studies is consistent with the new pan-tissue Treg model. For adipose Tregs, which are at the more distinct end of our multi-tissue analysis, phenotypic ^35^ and TCR clonality ^36^ distinctiveness were originally identified in comparison to Tregs from lymphoid organs. Likewise, the indefinite residency of adipose Tregs was derived from parabiosis experiments at 4-6 weeks ^36^, at which point spleen Tregs had normalized but adipose Tregs had not. It is only with the comparison to multiple additional non-lymphoid non-gut organs, and with the addition of a parabiotic time-course, that the seed and specialization model needs to be refined or replaced (of note, a parabiosis study at 12 weeks found no maintained residency for adipose Tregs ^27^, also consistent with our calculated dwell time). The pan-tissue model also allows us to reinterpret the complex data on tissue Treg “precursors” ^37^. PPARγ, originally identified as an adipose-specific Treg signature gene ^35^ but here observed more broadly, is also expressed in a smaller number of Tregs in lymphoid tissues. These cells are less diverse at the TCR level than other Tregs, and share clonality with Tregs from the liver, skin and adipose Tregs ^37^. Further, sorted PPARγ^+^ cells could “seed” all three non-lymphoid tissues tested ^37^. Similar findings are reported for KLRG1^+^NFIL3^+^ splenic Tregs, with a restricted TCR repertoire, overlapping with tissue Tregs, and an ability to seed tissues upon transfer ^38^. Under the seed and specialization model, the interpretation of this data required the postulation of tissue Treg “precursors” prior to permanent seeding, which problematically requires TCR specificity assortment preceding tissue entry. Under the pan-tissue Treg model, the lymphoid PPARγ^+^ Tregs can be reinterpreted as recirculating tissue Tregs, partially de-differentiated and captured during the lymphoid stage of a circular pan-tissue migration. While this idea had been considered and discarded in prior studies ^38^, this dismissal was based on KLRG1 being considered an irreversible marker, while our fate-mapper data demonstrates KLRG1 expression is reversible. The pan-tissue Treg model is therefore more consistent with the existing literature, even prior to the additional data generated here.

These data point to both conserved and disparate features between tissue Tregs and the conventional Trm populations. Beyond tissue-residency, the striking feature conserved between tissue Tregs and Trm is the transcriptional residency program, with markers such as CD69 and CD103 conserved across both populations. This Trm expression profile is unified across site and infection system ^15^. In the case of tissue Tregs, signature genes were initially identified in single non-lymphoid tissues and were labelled as unique in comparison to lymphoid Tregs. Thus markers such as KLRG1, ST2 and PPARγ were labelled as adipose-Treg specific ^35,39^, and Areg as muscle Treg-specific ^23^. The same markers, however, are expressed by Tregs from other non-lymphoid tissues, and when assessed in a pan-tissue manner, as here, they are near-ubiquitous. The tissue residency program in Tregs thus appears, like Trm, to be unified across tissues. We do not exclude “tilting” of the generic tissue-residency program by factors enriched in particular tissues, as observed in Trm ^40^, although this potential is reduced by the limited dwell-time of Tregs in most tissue. Further enrichment of these same genes in particular tissues, or sex-based comparisons ^27^, is more consistent with a relative enrichment of the generic tissue subset, rather than the presence of a unique population. This also holds true for DNA methylation studies and chromatin accessibility studies, where the largest differences are qualitative and observed between lymphoid and non-lymphoid Tregs, while differences between tissues are smaller and qualitative, with the same genes affected to different degrees across different tissues ^38,41^. The conservation of this signature, both across tissues and between tissue Tregs and Trm, suggests the importance of the constituent genes for the adaptation of lymphocytes to non-lymphoid tissues. We note, however, that it is plausible that a synchronous module is used across cell types, where not every gene within the module is necessary for each cell type that turns on the module. The phenotypic weakness of certain knockouts in tissue Tregs (e.g. CD69, CD103) may reflect higher importance in other cell types (i.e. Trm) expressing the same module.

The key difference between tissue Tregs and Trm is likely to be the nature of antigenic stimulation that turns on the residency module within the tissues. As conventional T cells, primed by infections, the antigen exposure of Trm is temporally- and spatially-restricted. Of necessity this requires the residency signature to be “locked in” without continual antigenic stimuli, a program which may utilise Hobit, Runx and Blimp ^16-18^. In the absence of such a transcriptional lock, conventional Trm would be diluted across tissues over time without a molecular signal present during recirculation to reinitiate residency, removing the advantage of site-memory. By contrast, homeostatic tissue Tregs recognise antigens without temporal restriction (although inflammation-responsive tissue Tregs may show these features) and with broader spatial distribution, enabling re-initiation on a continual basis. Indeed, in tissue Tregs, the absence of such a transcriptional lock, and the consequent sequential circulation-residency cycles, would enable optimisation of tissue distribution. While tissue Tregs do not appear to use a self-sustaining transcriptional lock, the expression of BATF does emerge as a requirement for the transient expression of the residency module, with BATF-deficiencies identified here and elsewhere ^42^ as negatively impacting the number of Tregs present across a range of non-lymphoid organs. Indeed, analysis of the reoccurring tissue Treg transcriptional program identifies BATF as a common binding motif, both within the tissues and in the ST2^+^ population present within lymphoid tissues ^38^. BATF in tissue Tregs may therefore fulfil a similar function to Hobit in Trm, but with the driver being continual antigenic exposure rather than a single activation step.

An additional feature of the tissue Treg pool is likely to be an adaptation to tissue inflammation. While here we largely concentrated on the homeostatic state, based on published experiments it is likely that tissue inflammation leads to an expansion of the residency niche (i.e., the number of Tregs the tissue is capable of housing), an intensification of the residency markers, and a potential change in the TCR repertoire in response to changing antigen presentation. The intense tissue residency phenotype observed in adipose Tregs ^27,33^ likely reflects this inflammatory effect, with the male adipose tissue in some mouse colonies showing molecular characteristics of inflammation, driven by sex hormones ^27^. Likewise, Kohlgruber et al demonstrated that the elevated ST2^+^ Treg number in the adipose tissue was dependent on IL17 and γδ T cells, demonstrating that this is a *bona fide* inflammatory phenotype ^43^. In our mouse colonies, the homeostatic state of adipose Tregs was less striking (a colony-dependent effect has been noted ^44^), however the niche was expanded and the phenotype intensified by age, and altered by microbiome, consistent with an inflammation-dependent impact. Inflammation-driven phenotypic intensification is likely to require Blimp1 and Id2 ^45,46^, with niche expansion driven by the release of IL33 alarmin following inflammation, acting on the ST2 receptor on tissue Tregs ^36^. As the adipose tissue in our colony was not inflamed, no major phenotypes of Blimp1 or ST2 were observed in the adipose Tregs. In this regard, the inflamed adipose is similar to aged/injured muscle ^47,48^, the inflamed heart ^49^, and the damaged kidney ^50^, where IL33 is released and ST2 becomes a potent regulator. In the muscle, this inflammation is accompanied by a change in the tissue Treg TCR repertoire, with at least one TCR clone showing enhanced preference for inflamed over non-inflamed muscle ^51^. Likewise in the heart, the expanded tissue Treg niche is driven by external influx of new Tregs, with an altered TCR repertoire, rather than *in situ* proliferation of existing cells ^49^. Whether a dependency on Blimp1/Id2 and IL33-ST2 for tissue Treg expansion during inflammation is shared across tissues, and whether the same recirculating Treg clones are attracted to inflammation in a pan-tissue manner, remains to be seen. This intriguing possibility is, however, consistent with the pre-priming of tissue Tregs around the body to respond to IL33 (i.e., the expression of ST2) even in the absence of IL33 release. Likewise the universality of core inflammation transcriptional changes across tissues ^52^ at least raises the potential that some Treg clones will be capable of enhanced tissue-residency during inflammation in multiple tissues.

A shared residency program initiated in pan-tissue Tregs is not mutually exclusive with tissue-specific functional endpoints. From an evolutionary perspective, repurposing of homeostasis-restoring effector molecules on a tissue-by-tissue basis would enable a single regulatory module in tissue Tregs to drive disparate physiological responses. Thus, effector molecules for tissue homeostasis, such as Notch ligand expression by skin Tregs ^26^, could allow a single mediator to feed into diverse responses across multiple tissues. While largely hypothetical until all key mediators are identified, a clear example of this pleiotropy is the production of amphiregulin by Tregs. First proposed as a muscle Treg-specific effector molecule capable of enhancing injury repair processes ^23^, Areg is widely produced by Tregs. Reports suggest that Treg-derived Areg is capable of suppressing neurotoxic astrogliosis in the brain ^53^, driving alveolar regeneration in the lung ^54,55^, regulating angiogenesis and vascular remodeling in the limbs ^56^, suppressing renal and hepatic inflammation ^57,58^, and driving corneal healing in the eye ^59^. In each case, source and effector molecule are conserved, while the response cell and initiated reparative program differ markedly. We can thus predict further conservation of pan-tissue effector molecules while also expecting to unravel novel and unusual functional impacts of tissue Tregs in diverse tissues. Finally, it is worth noting that “cancer Tregs” are likely to be a maladaptation of tissue Tregs. A phenotypic capture and exploitation of an existing program is more likely than an evolutionary pressure creating a “cancer Treg” program preventing tumor clearance. Indeed, cancer Tregs are now recognized to acquire the features of fat Tregs ^60^, i.e., the common tissue-residency markers described here, and Areg is a known mediator of anti-tumour immune suppression by cancer Tregs ^61^. In this and other areas, tissue Treg biology will likely lead to surprising advances across physiology and pathology.

## Acknowledgements

This work was supported by the ERC Consolidator Grant TissueTreg (to A.L.), the Wellcome Trust (to A.L.), the Biotechnology and Biological Sciences Research Council through Institute Strategic Program Grant funding BBS/E/B/000C0427 and BBS/E/B/000C0428, and the Biotechnology and Biological Sciences Research Council Core Capability Grant to the Babraham Institute. The authors acknowledge the important contributions of Jeason Haughton (VIB) and the Babraham Institute BSU for mouse husbandry, the KU Leuven FACS Core and Babraham Institute Flow Cytometry Core for flow cytometry, and the Babraham Institute Sequencing and Bioinformatics Cores for RNAseq acquisition and analysis. We thank Amy Dashwood, Lidia Yshii, Ana Laura Calvanese, Wenson Karunakaran, Stephanie Lienart, Kailash Singh, Alena Moudra, Ntombizodwa Makuyana and Daria Vdovenko for technical assistance. For mouse strains, we acknowledge Lars Vereecke (VIB) for germfree mice, Andrew McKenzie (LMB) for ST2 KO and LFA1 KO mice, Alexander Rudensky (Sloan Kettering) for *Foxp3^Thy1.1^* reporter and Foxp3^CreERT2^ mice, Michelle Linterman (Babraham Institute) for UbGFP mice, Rahul Roychoudhuri (University of Cambridge) for CD4Cre Blimp1-flox mice, Pilar Lauzurica (Instituto de Salud Carlos III) for CD69 KO mice, Laurent Brossay (Brown University) for KLRG1 KO mice, Takaharu Okada (RIKEN) for KLRG1Cre mice, and Alex Gould (Francis Crick Institute) for Hif1α-deficient mice. Murine TCR OTII-2A.pMIG II was a gift from Dario Vignali (University of Pittsburg).

