## Supplementary material for "The tissue-resident regulatory T cell pool is shaped by transient multi-tissue migration and a conserved residency program"

**Supplementary methods**

*Mouse strains and strain generation*

All mice were housed in SPF conditions unless otherwise stated. All mice used were on the C57BL/6 background. *Foxp3$^{Thy1.1}$* and CD45.1 *Foxp3$^{Thy1.1}$* mice [1], *Klrg1$^{Cre}$ RosaAi14* mice [2], *CD69$^{-/-}$* mice [3], *Rag1$^{-/-}$* mice [4], *CD11a$^{-/-}$* (*Itgal*$^{tm1Bll}$) mice [5], CD4Cre *Hif1a$^{fl/fl}$* mice [6], Ubiquitin-GFP mice [7], *Klrg1$^{-/-}$* mice [8], *S1pr2$^{-/-}$* mice (*S1pr2$^{tm2a(EUCOMM)Hmgu}$*), CD4Cre *Blimp1$^{fl/fl}$* [9] and *ST2$^{-/-}$* (*Il1rl1$^{tm1Anjm}$*) mice [10] are as previously described. *BATF$^{-/-}$* bone-marrow (013758) was obtained from JAX for bone-marrow reconstitution.

*CD69$^{fl/fl}$* mice were generated using C57BL/6N ES cells, through Cyagen. Briefly, exons 2-4 were targeted for flox site insertion, using homologous arms generated using BAC clones RP23-11N22 and RP23-399O9 as templates. The targeting vector included a self-deleting Neo cassette site, leaving the *CD69$^{fl/fl}$* mice. Cre activity on this allele results in the deletion of 2382bp spanning exons 2-4, resulting in a frameshift after exon 1 (residue 21) and a stop codon after residue 24. Mice were then crossed to the *Foxp3$^{CreERT2}$* strain [11].

*CD103$^{-/-}$* mice were generated using direct microinjection of Cas9 protein (IDT, 1µg/µl) and *CD103* gRNA (5µl of 1µM, 5’ GTTGAGGCTGCGGAACTTCC-3’ targeting exon 5) into C57BL/6 zygotes, followed by transfer into CD1 surrogates. Offspring were screened, and mice bearing a 10bp deletion (aagttccgca) resulting in a frameshift at residue 118 were bred to homozygosity and confirmed for lack of expression by flow cytometry.

*Areg$^{-/-}$* mice were generated using direct microinjection of Cas9 protein (IDT, 1µg/µl) and *Areg* gRNA (5µl of 20µM mixed gRNA 5’-TCTGGGGACCACAGTGCCGGTGG-3’ and 5’-ATAATATAGCCGGATATTTGTGG-3’ targeting exon 2, plus 5µl of 20µM trRNA) into C57BL/6 zygotes, followed by transfer into CD1 surrogates. Offspring were screened by sequencing, and mice bearing a 118bp deletion in exon 2 resulting in a frameshift and early truncation were bred to homozygosity and confirmed for lack of expression by flow cytometry.

*Mouse procedures*

Gnotobiotic C57BL/6J mice were housed at the VIB gnotobiotic facility, Ghent. For microbiome enrichment, pet store female mice were wild-exposed prior to cohousing with SPF C57BL/6J mice (KUL L2 animal facility, Leuven). After 12 weeks of cohousing, a subset of mice was tested for 47 pathogens by serology and screened for microbiome diversity by fecal microbiome sequencing, as previously reported [12].

For parabiosis, Foxp3$^{Thy1.1}$ and CD45.1 Foxp3$^{Thy1.1}$ mice were intercrossed for at least 5 generations prior to starting parabiosis experiments. The congenic mice were then cohoused for one week prior to surgery. Parabiosis surgery was performed as previously described [13]. Mice in which the blood lymphocyte population was not between 40-60% of donor origin after 1 week were excluded from the study. For male:female parabiosis, the two mice were matched in weight rather than age.

*Foxp3*$^{ERT2Cre}$ *CD69*$^{fl/fl}$ mice were treated with tamoxifen (0.2mg/g) by oral gavage weekly for three weeks.

Competitive mixed bone marrow chimeras were created using equal mixtures of bone marrow. A total of 2x10$^{6}$ cells were injected i.v. into 11 Gy-irradiated CD45.1/CD45.2 heterozygous mice or 5.5Gy-irradiated RagKO mice. Mice were given water containing Baytril (50mg/ml) after irradiation. Chimeras were analyzed 8-12 weeks post-reconstitution.

For retrogenic TCR mice, hematopoietic stem cells (HSC) were isolated as lineage-depleted bone marrow cells using the EasySep Mouse Hematopoietic Progenitor Cell Isolation kit (Stem Cell Technologies), and transduced with retrovirus in the presence of Lentiblast (Oz Biosciences). HSC were cultured in TPO (R&D Systems), IL-3, IL-6, SCF and FLT3L (BioLegend) for 24hr in StemSpan SFEM II medium (Stem Cell Technologies). The transduced cells (5x10$^{5}$ per mouse) were transplanted into lethally irradiated C57BL/6 mice along with unmanipulated *Rag1*$^{-/-}$ bone marrow (1x10$^{6}$ cells). Mice were assessed 10 weeks post-transplant.

To test tissue re-homing, leukocytes were isolated from the tissues of congenic strains (CD45.1, CD45.2, CD45.1/CD45.2, Foxp3$^{Thy1.1}$, CD45.1 Foxp3$^{Thy1.1}$, UbGFP, CD45.1 UbGFP, RosaRFP) and combined such that each pool contained uniquely identifiable Tregs from each tissue source. The pools were injected i.p. into individual Rag-deficient mice. Cells were recovered and analyzed four weeks post-transfer.

*Tissue preparation*

Prior to isolating tissue leukocytes, mice were either injected intravenously with anti-CD45 antibody to label vascular leukocytes or they were lethally anesthetized with ketamine/xylazine and perfused with PBS containing 2.5% FCS. Tissues were harvested into HBSS with 2.5% FCS and 2mM EDTA, and stored on ice until further processing. The thymus, lymph nodes, Peyer's patches, spleen, adrenal glands were dispersed with frosted glass microscope slides, filtered through 100μm mesh. Where necessary, erythrocytes were lysed prior to counting using a Countess Automated Cell Counter (ThermoFisher). The liver was prepared similarly, except the resulting cell suspension was washed extensively and centrifuged (600g, 10min) through a 40% Percoll (Sigma Aldrich) solution prior to erythrocyte lysis. Bone marrow was extracted from a single femur by crushing with a mortar and pestle. Blood was treated to lyse red cells prior to staining.

The mesentery, salivary glands, lungs, pancreas, kidney, adipose tissue, reproductive tissues, eyes, back skin, tongue, hind limb muscle, heart, bladder and brain were digested as follows to extract tissue leukocytes: Tissues were chopped finely with razor blades and washed by centrifugation to remove debris. They were then resuspended in digest buffer (IMDM supplemented with 20% FCS, 10mM HEPES, 1mM sodium pyruvate, 10μg/ml gentamicin, 1mM $CaCl_2$ and 1mM $MgCl_2$) with enzymes (400μg/ml collagenase D, 100μg/ml hyaluronidase and 40μg/ml DNase I, Sigma Aldrich). In some experiments 2mg/ml collagenase IV was used in place of collagenase D. Tissues were shaken in an agitating mixer at 37°C for 15min, dispersed with a pipette, and shaken for another 15min. The solution was then filtered through 100μm

mesh and any remaining chunks were forced through the mesh. The cells were washed and passed through 40% Percoll to enrich for the leukocyte fraction.
The various parts of the gastrointestinal tract were separated into intraepithelial and lamina proprial compartments using a pre-digest step of agitation at 37°C in HBSS with 10mM HEPES, 10mM EDTA and 2.5% FCS. The released cells were collected and passed through 40% Percoll to isolate the intraepithelial leukocytes (IELs). The remaining tissue was diced finely, digested enzymatically and passed through Percoll to isolate the lamina propria leukocytes (LPLs).

*Murine tissue flow cytometry*

For murine samples, Fc receptor interactions were blocked using 2.4G2 hybridoma supernatant. Viability staining was performed in HBSS, using either ViaKrome 808 (Beckman Coulter) or Fixable Viability Dye eFluor780 (ThermoFisher). *In vivo* intravenous labeling of leukocytes with anti-CD45-biotin was developed using fluorescent streptavidin. Any epitopes requiring pre-fixation staining were stained for 1 hour at 4°C in the dark in FACS buffer (PBS with 2.5% FCS and 2mM EDTA). Cells were washed with FACS buffer and then fixed. For fluorescent protein retention, cells were fixed at room temperature for 45 minutes in the dark with neutral buffered formalin (VWR). For all other purposes, cells were fixed and permeabilized for 30 minutes using the Foxp3/Transcription Factor Staining Buffer Set (eBioscience). Following two washes with the eBioscience permeabilization buffer, cells were stained overnight (~16hrs) with all remaining antibodies in permeabilization buffer supplemented with 20% 2.4G2 supernatant, as described [14]. Cells were washed and acquired with 10,000 Precision Count beads (BioLegend) added per sample. The antibody cocktail used for the Nur77 stain set is given in **Supplementary Table 1**. The antibody cocktail used for the Aurora tissue Treg enumeration panel is given in **Supplementary Table 2**. The antibody cocktail used for the Aurora mouse bone-marrow chimera panel is given in **Supplementary Table 3**. The antibody cocktail information used for the Aurora mouse adoptive cell transfer panel is given in **Supplementary Table 4**. The antibody cocktail information used for the Symphony parabiosis panel is given in **Supplementary Table 5**. For ageing and cohousing data, the Symphony parabiosis panel was used, with the exception of including Alexa Fluor 488 anti-TCRβ (clone H57-597, BioLegend, 109215, 1:2500) in place of CD45.1.

Where indicated, antibodies were conjugated to fluorophores according to the manufacturer's instructions. Where possible, free fluorophore was removed using centrifugal filters with a 50kDa cut-off. The following antibody conjugation kits were used in this research: PE/Cy5.5® Conjugation Kit - Lightning-Link®, APC/Cy5.5® Conjugation Kit - Lightning-Link® (AbCam), Alexa Fluor™ 350 Antibody Labeling Kit, Alexa Fluor™ 532 Antibody Labeling Kit, and Alexa Fluor™ 790 Antibody Labeling Kit (ThermoFisher).

Cohousing, gnotobiotic, parabiosis, ageing and $CD103^{-/-}$ chimera data were acquired on a BD FACSymphony A5 flow cytometer (BD Biosciences). The remaining murine data were acquired on an Aurora Spectral Analyzer (Cytek).

Flow cytometry data are available for download on FlowRepository at the links below.

| **Dataset** | **Figure** | **FlowRepository ID** |
|---|---|---|
| Tissue screen | Figure 1A-G, | FR-FCM-Z6ME |

| | | |
|---|---|---|
| | Supplementary Figure 1, Supplementary Figure 4 | |
| Nur77 | Figure 1H,I | FR-FCM-Z6L8 |
| Ageing | Figure 2A-D, Supplementary Figure 5 | FR-FCM-Z6L9 |
| Microbiome | Figure 2E-I, Supplementary Figure 5 | FR-FCM-Z6LT |
| CD69 KO chimera | Figure 4, Supplementary Figure 6 | FR-FCM-Z6MF |
| ST2 KO chimera | Figure 4, Supplementary Figure 7 | FR-FCM-Z6MK |
| CD11a KO chimera | Figure 4, Supplementary Figure 8 | FR-FCM-Z6ML |
| KLRG1 KO chimera | Figure 4, Supplementary Figure 9 | FR-FCM-Z6MJ |
| CD4Cre *Blimp1*$^{fl/fl}$ chimera | Figure 4, Supplementary Figure 10 | FR-FCM-Z6MW |
| S1PR2 chimera | Figure 4, Supplementary Figure 11 | FR-FCM-Z6MH |
| BATF KO chimera | Figure 4, Supplementary Figure 12 | FR-FCM-Z6MM |
| CD4Cre *Hif1a*$^{fl/fl}$ chimera | Figure 4, Supplementary Figure 13 | FR-FCM-Z6MN |
| Areg KO | Figure 4, Supplementary Figure 14 | FR-FCM-Z6MS |
| CD103 KO | Figure 4, Supplementary Figure 15 | FR-FCM-Z6MR |
| Parabiosis | Figure 5, Supplementary Figures 17, 18 | FR-FCM-Z5UV |
| Female:male parabiosis | Figure 8A-B | FR-FCM-Z6MQ |
| i.v. CD45 labeling | Supplementary Figure 2 | FR-FCM-Z6MX |
| *Foxp3*$^{ERT2Cre}$ *CD69*$^{fl/fl}$ | Supplementary Figure 16 | FR-FCM-Z6MG |
| *Klrg1*$^{Cre}$ *RosaAi14* fate mapper | Supplementary Figure 19 | FR-FCM-Z6MP |

*FlowCode cloning*

TCR coding sequences from murine TCR OTII-2A.pMIG II [15] (Addgene plasmid # 52112 ; http://n2t.net/addgene:52112 ; RRID:Addgene_52112) were removed using EcoRI and XhoI and replaced by TCR of interest coding sequences using NEBuilder Hifi DNA assembly and IDT eblocks gene fragments encoding Travj, Trac_P2A, Trbvdj, Trbc1 or Trbc2. EGFP was removed using NcoI and NotI and replaced by H2bc3_AU1_Flag_HA coding sequence using NEBuilder Hifi DNA assembly and an IDT eblocks gene fragment. AU1_Flag_HA was removed using BamHI and NotI and replaced by other Procode tags using NEBuilder Hifi DNA assembly and PCR amplicons generated from NLS-mCherry Pro-Code vector library [16,17].

*RNAseq*

For bulk RNA sequencing, 2000 CD4$^+$ Foxp3Thy1.1$^+$ Tregs and Foxp3Thy1.1$^-$ Tconv were sorted on a BD FACSAria from each source from perfused mice. RNA was isolated using RNeasy Mini kit (Qiagen). RNA concentration and purity were determined using the Nanodrop ND-1000 (Nanodrop Technologies) and RNA integrity with a Bioanalyser 2100 (Agilent). 3′mRNA-seq library preparation and transcriptome analysis was performed by Lexogen (Austria) using the QuantSeq 3′mRNA-Seq Library Prep Kit for Illumina and QuantSeq data analysis workflow. scRNA-Seq was performed using 10x Genomics 5' VDJ Single Cell Immune Profiling. Tregs were flow sorted on a BD Influx, Aria, Jazz or Fusion on the basis of CD4$^+$Foxp3Thy1.1$^+$ i.v.CD45$^-$ CD19$^-$CD11b$^-$CD8$^-$F4/80$^-$. Cells were labeled with Hashtag TotalSeq regents and loaded onto the 10x Chromium Controller. Sequencing was performed on an Illumina HiSeq. Data was processed in R using scripts as detailed below.

Following sequencing of purified tissue Treg and Tconv populations, the RNAseq data was processed using FastQC [18] for assessing read quality pre-and post-trimming, Trim Galore! [19] for trimming adapter sequences, and STAR aligner [20] for aligning to GENCODE [21] mouse primary assembly vM23, with the raw counts being generated by featureCounts [22]. A combined report was generated using MultiQC [23]. Raw counts were normalised using DESeq2 [24], and global analysis using PCA, t-SNE [25,26] (perplexity parameter at '3') and UMAP [27,28] was performed on the genes post minimum transcript per cell filtering of 0.01 transcripts per cell. Heatmaps were produced using Pearson correlation. Differential expression analysis was performed using the default DESeq2 analysis pipeline, but without using zero-centered Normal priors, log2 fold change shrinkage, outlier replacement using Cook's distance or independent filtering. The selection criteria was absolute DESeq log2 beta $\geq 2$ and adjusted p.value $< 0.01$. Gene set enrichment analysis was performed using GAGE [29] with the pathway visualisation done by Pathview [30]. The script for the RNAseq analysis is available at https://github.com/AdrianListon/TissueTregs. The enriched profiles were compared using the clusterProfiler package [31]. A publicly accessible webtool was developed in R Shiny [32] which provides an interactive bulk RNA-seq data analysis module allowing users to perform custom contrasts to explore the data further for additional results and/or verification. In addition to this module, other modules are also available, such as the TCR module which explores the T cell repertoire in the data. The script for generating the webtool is available at https://github.com/AdrianListon/ExpressionViewer. The webtool is available at https://www.bioinformatics.babraham.ac.uk/shiny/expressionViewer/.

For TCR sequence analysis, scRNA data was used to allow for paired TCRαβ chain identification. Row sequencing data was processed with CellRanger 5.0.0 without barcode mismatches allowed and with multi-features analysis (gene expression, vdj-t, totalseq hashtags). Features were linked based on cell barcodes and tissue origin was retrieved based on hashtags signals when highest hashtag count was at least 30 times higher than the others using a custom R script. Gene expression and TCR repertoire analysis were generated in R using circlize, dplyr, ggplot2, ggsci, Matrix, pheatmap,scales, scater, Seurat, SingleCellExperiment, tidyr, viridis packages. For repertoire analysis, a TCR clonotype is associated to a unique set of Trav, tra_cdr3, trbv, trb_cdr3. The custom script is available at: https://github.com/AdrianListon/TissueTregTCR.

*Statistical analysis*

FlowJo version 10.8.1 (BD), GraphPad Prism 9, R version 4.1.2, RStudio, and CellRanger (10X Genomics) were used for analysis. Within the R environment, the following packages were employed: digest, dunn.test, flowCore, ggplot2, ggridges, RANN, RColorBrewer, reshape2, EmbedSOM, Rtsne, umap, ConsensusClusterPlus, FlowSOM, EmbedSOM, Seurat and dplyr. Flow cytometry spillover compensation was calculated using AutoSpill [33].

Statistical tests used are indicated in the figure legends, typically a two-way ANOVA with Šídák's multiple comparisons. Statistical comparisons of tSNE and UMAP plots were performed with the Cross Entropy test, using Kolmogorov-Smirnov tests with Holm correction [34]. The scripts used for flow cytometry data analysis and tSNE Cross Entropy comparisons are available on GitHub: https://github.com/AdrianListon/Cross-Entropy-test. Immunopeptidome reanalysis was based on the published $H2D^d$ immunopeptidome of 19 normal tissues from C57BL/6 mice [35].

Parabiosis data (proportions of cell states (resting, activated, CD69+ Tregs) in each of 16 tissues and blood measured at weeks 0, 1, 2, 4, 8, 12 in each parabiont) was modelled by continuous time Markov Chains using a Bayesian approach. The aim was to describe the observed data (flow cytometry data of T cell types and state proportions evolving in time), while directly estimating the biological flow/transition rates including their distributions, i.e., the biological process governing the observed data. Independently for each tissue, the Markov Chain models the whole body Treg dynamics using 9 model states (3 body compartments (blood, selected tissue, all other tissues pooled) x 3 cell states (resting, activated, CD69+)) and possible flows between them. The assumed prior distributions of flow rates were selected as uniform on the support 2-3 orders of magnitude larger than the final posterior estimates in order to provide the least information and robustness. The models were fitted using Markov Chain Monte Carlo (MCMC) algorithm, specifically Hamiltonian Monte Carlo (HMC) in the Stan programming language, while using the rstan 2.26 package, interface for R language. Each Markov Chain model was estimated by four Monte Carlo chains with 5,000 samples. The flow rate estimates were provided as mode posterior together with 80% highest density (credible) interval (HDI). The full pipeline is available at https://github.com/gergelits/markov_chain_tissue_treg.

Retrogenic TCR analysis via FlowCodes was assessed using FlowJo v10.8.1 to extract CD4 T cell-related events and then analyzed with the custom FlowCode Decoder shiny app to obtain count and expression levels per TCR clone. The FlowCode Decoder script is available on GitHub: https://github.com/obricard/FlowcodeDecoder. Graphs were generated in R 4.1.2 using ggplot2.

**Supplementary Table 1. Antibody information for Aurora Nur77 panel**.

| Vendor | Catalogue # | Antibody | Clone | Dilution |
|---|---|---|---|---|
| BD | 740238 | BUV395 anti-CD103 | M290 | 500 |
| BD | 612952 | BUV496 anti-CD4 | GK1.5 | 500 |
| BD | 612833 | BUV737 anti-CD62L | MEL-14 | 1000 |
| BD | 612971 | BUV661 anti-CD19 | 1D3 | 2000 |
| BD | Prototype | BUV615 anti-CTLA-4 | UC10-4F10-11 | 1000 |
| BD | 747971 | BV480 anti-CCR2 | 475301 | 200 |
| BD | 564722 | BV650 anti-RORγT | Q31-378 | 500 |
| BD | 747298 | BV750 anti-CXCR3 | CXCR3-173 | 500 |
| BD | Prototype | BB660-P2 anti-CD95 | Jo2 | 1000 |
| BD | Prototype | BB790 anti-Ly-6C | AL-21 | 2000 |
| BD | 565577 | BB515 anti-CCR9 | CW-1.2 | 200 |
| BioLegend | 117330 | BV421 anti-CD11c | N418 | 2000 |
| BioLegend | 103044 | BV510 anti-CD44 | IM7 | 1000 |
| BioLegend | 135231 | BV711 anti-PD-1 | 29F.1A12 | 5000 |
| BioLegend | 104510 | PE-Cy5 anti-CD69 | H1.2F3 | 200 |
| BioLegend | 100260 | Spark Blue 550 anti-CD3 | 17A2 | 5000 |
| BioLegend | 107624 | PerCP anti-MHCII | M5/114.15.2 | 200 |
| BioLegend | 138429 | BV785 anti-KLRG1 | 2F1/KLRG1 | 500 |
| BioLegend | 321224 | APC/Fire 750 anti-Integrin β7 | FIB504 | 200 |
| eBioscience | Q10091MP | Qdot545 Streptavidin | N/A | 1000 |
| eBioscience | 56-5698-82 | Alexa Fluor 700 anti-Ki67 | SolA15 | 1000 |
| eBioscience | 46-3041-82 | PerCP-eFluor710 anti-CD304 | 3DS304M | 5000 |
| eBioscience | 12-5965-82 | PE anti-Nur77 | 12.14 | 200 |
| eBioscience | 50-9858-82 | eFluor 660 anti-IRF4 | 3E4 | 5000 |
| eBioscience | 17-5773-82 | APC anti-Foxp3 | FJK-16s | 200 |
| eBioscience | 48-9883-41 | eFluor 450 anti-Helios | 22F6 | 500 |
| Beckman Coulter | C36628 | ViaKrome 808 | N/A | 300 |
| Miltenyi | 130-111-601 | APC anti-Foxp3 | REA788 | 200 |

| | | | | |
|---|---|---|---|---|
| BioRad | MCA1260SBV515 | SBV515 anti-CD25 | PC61 | 200 |
| BioRad | MCA1266SBV610 | SBV610 anti-NK1.1 | PK136 | 200 |

**Supplementary Table 2: Antibody information for Aurora mouse Treg enumeration panel.**

| Vendor | Catalogue # | Antibody | Clone | Dilution |
|---|---|---|---|---|
| BD | 564279 | BUV395 anti-CD45 | 30-F11 | 2000 |
| BioLegend | 14-0242-82 | Purified anti-CD24 | M1/69 | 250 |
| eBioscience | A20180 | Alexa Fluor 350 conjugation kit | N/A | N/A |
| BD | 612952 | BUV496 anti-CD4 | GK1.5 | 500 |
| BD | 741233 | BUV563 anti-NK1.1 | PK136 | 500 |
| BD | 624297 | BUV615 anti-Siglec F | E50-2440 | 1000 |
| BD | 612971 | BUV661 anti-CD19 | 1D3 | 2000 |
| BD | 612833 | BUV737 anti-CD62L | MEL-14 | 1000 |
| BD | 742061 | BUV805 anti-GITR | DTA-1 | 500 |
| BD | 612971 | BUV661 anti-CD19 | 2E7 | 500 |
| BioLegend | 121422 | BV421 anti-CD103 | 2E7 | 500 |
| Biotechne | FAB69501V-100UG | Alexa Fluor 405 anti-CD11c | N418 | 500 |
| eBioscience | 48-5871-82 | eFluor450 anti-TIM-3 | 8B.2C12 | 200 |
| BD | 747971 | BV480 anti-CD192 | 475301 | 1000 |
| BioLegend | 103044 | BV510 anti-CD44 | IM7 | 2000 |
| BioLegend | 105329 | BV570 anti-CD90.2 | 53-2.1 | 5000 |
| eBioscience | MF48030 | Pacific Orange anti-F4/80 | BM8 | 500 |
| eBioscience | 13-0451-85 | Biotin anti-CD45 | 30-F11 | 2µg/mouse |
| eBioscience | Q10091MP | Qdot545 Streptavidin | N/A | 300 |
| BioLegend | 313538 | BV605 anti-ICOS | C398.4A | 500 |
| BioLegend | 148220 | BV650 anti-XCR1 | ZET | 1000 |
| BioLegend | 135231 | BV711 anti-PD-1 | 29F.1A12 | 2000 |
| BD | 747298 | BV750 anti-CD183 | CXCR3-173 | 500 |
| eBioscience | 78-5893-80 | Super Bright 780 anti-KLRG1 | 2F1 | 500 |
| BD | 565577 | BB515 anti-CD199 | CW-1.2 | 500 |
| eBioscience | 11-5711-82 | FITC anti-TCRγδ | GL3 | 200 |
| BioLegend | 139302 | Purified anti-CD64 | X54-5/7.1 | 1000 |
| eBioscience | A20182 | Alexa Fluor 532 conjugation kit | N/A | N/A |
| BioLegend | 100260 | Spark Blue 550 anti-CD3 | 17A2 | 500 |

| | | | | |
|---|---|---|---|---|
| BD | 624295 | BB660-P2 anti-CD95 | Jo2 | 2000 |
| Biolegend | 107624 | PerCP anti-I-A/I-E | M5/114.15.2 | 200 |
| eBioscience | 46-3041-82 | PerCP-eFluor710 anti-CD304 | 3DS304M | 2000 |
| BD | Prototype | BB755-P anti-CD8a | 53-6.7 | 500 |
| BD | 624296 | BB790-P anti-Ly-6C | AL-21 | 2000 |
| BioLegend | 113406 | PE anti-CD120b | TR75-89 | 5000 |
| BioLegend | 129822 | PE-Dazzle 594 anti-CD196 | 29-2L17 | 500 |
| BioLegend | 104510 | PE-Cy5 anti-CD69 | H1.2F3 | 200 |
| eBioscience | 35-0251-82 | PE-Cy5.5 anti-CD25 | PC61 | 1000 |
| eBioscience | 25-9335-80 | PE-Cy7 anti-ST2 | RMST2-2 | 5000 |
| eBioscience | 17-0900-82 | APC anti-CD90.1 | HIS51 | 500 |
| BioLegend | 144508 | Alexa Fluor 647 anti-CD193 | J073E5 | 5000 |
| eBioscience | 14-0112-82 | Purified anti-CD11b | M1/70 | 10000 |
| AbCam | ab102855 | APC-Cy5.5 conjugation kit | N/A | N/A |
| Biotechne | FAB72671N-100UG | Alexa Fluor 700 anti-TIGIT | 2190A | 200 |
| eBioscience | 65-0865-18 | Fixable Viability Dye eFluor780 | N/A | 4000 |
| BioLegend | 127602 | Purified anti-Ly-6G | 1A8 | 5000 |
| eBioscience | A20189 | Alexa Fluor 790 conjugation kit | N/A | N/A |

**Supplementary Table 3**: **Antibody information for Aurora mouse bone marrow chimera panel.**

| Vendor | Catalogue # | Antibody | Clone | Dilution |
|---|---|---|---|---|
| BD | 740238 | BUV395 anti-CD103 | M290 | 500 |
| BD | 612952 | BUV496 anti-CD4 | GK1.5 | 500 |
| BD | 741233 | BUV563 anti-NK1.1 | PK136 | 500 |
| BD | 624297 | BUV615 anti-CD152 | UC10-4F1 | 1000 |
| BD | 612971 | BUV661 anti-CD19 | 1D3 | 2000 |
| BD | 612833 | BUV737 anti-CD62L | MEL-14 | 1000 |
| BD | 564920 | BUV805 anti-CD8 | 53-6.7 | 500 |
| BD | 750718 | BUV661 anti-CD103 | 2E7 | 500 |
| BioLegend | 110732 | BV421 anti-CD45.1 | A20 | 1000 |
| BD | 747971 | BV480 anti-CD192 | 475301 | 200 |
| BioLegend | 103044 | BV510 anti-CD44 | IM7 | 1000 |
| BioLegend | 137220 | Pacific Blue anti-Helios | 22F6 | 500 |
| eBioscience | 13-0451-85 | Biotin anti-CD45 | 30-F11 | 2µg/mouse |
| eBioscience | Q10091MP | Qdot545 Streptavidin | N/A | 300 |
| BioLegend | 313538 | BV605 anti-ICOS | C398.4A | 500 |
| BD | 564722 | BV650 anti-RORγT | Q31-378 | 500 |
| BioLegend | 135231 | BV711 anti-PD-1 | 29F.1A12 | 5000 |
| BD | 747298 | BV750 anti-CD183 | CXCR3-173 | 500 |
| eBioscience | 78-5893-80 | Super Bright 780 anti-KLRG1 | 2F1 | 1000 |
| BD | 565577 | BB515 anti-CD199 | CW-1.2 | 200 |
| BioLegend | 109862 | Spark Blue 550 anti-CD45.2 | 104 | 5000 |
| BD | 624295 | BB660-P2 anti-CD95 | Jo2 | 1000 |
| eBioscience | 46-3041-82 | PerCP-eFluor710 anti-CD304 | 3DS304M | 5000 |
| BD | 624296 | BB790-P anti-Ly-6C | AL-21 | 2000 |
| BioLegend | 113406 | PE anti-CD120b | TR75-89 | 5000 |
| BioLegend | 644833 | Alexa Fluor 594 anti-T-bet | 4B10 | 500 |
| eBioscience | 61-9966-41 | PE-eFluor610 anti-GATA-3 | TWAJ | 200 |
| BioLegend | 104510 | PE-Cy5 anti-CD69 | H1.2F3 | 200 |

| | | | | |
|---|---|---|---|---|
| eBioscience | 35-0251-82 | PE-Cy5.5 anti-CD25 | PC61 | 1000 |
| eBioscience | 25-9335-80 | PE-Cy7 anti-ST2 | RMST2-2 | 5000 |
| eBioscience | 17-5773-82 | APC anti-Foxp3 | FJK-16s | 200 |
| Miltenyi | 130-111-601 | APC anti-Foxp3 | REA788 | 200 |
| eBioscience | 50-9858-82 | eFluor660 anti-IRF4 | 3E4 | 5000 |
| eBioscience | 56-5698-82 | Alexa Fluor 700 anti-Ki67 | SolA15 | 1000 |
| eBioscience | 65-0865-18 | Fixable Viability Dye eFluor780 | N/A | 4000 |

**Supplementary Table 4: Antibody information for Aurora mouse adoptive transfer panel.**

| Vendor | Catalogue # | Antibody | Clone | Dilution |
|---|---|---|---|---|
| BioRad | Prototype | StarBright UV400 anti-F4/80 | Cl:A3-1 | 1000 |
| BioRad | Prototype | StarBright UV445 anti-CD45R | RA3-6B2 | 1000 |
| BioRad | MCA500SBV440 | StarBright Violet440 anti-CD3 | KT3 | 500 |
| BioRad | MCA1260SBV515 | StarBright Violet515 anti-CD25 | PC61 | 1000 |
| BioRad | MCA1266SBV610 | StarBright Violet610 anti-CD161 | PK136 | 200 |
| BioRad | STAR210SBV670 | StarBright Violet670 Streptavidin | N/A | 10 |
| BD | 612952 | BUV496 anti-CD4 | GK1.5 | 500 |
| BD | 612833 | BUV737 anti-CD62L | MEL-14 | 1000 |
| BD | 564920 | BUV805 anti-CD8 | 53-6.7 | 500 |
| BD | 624297 | BUV615 anti-CD152 | UC10-4F1 | 1000 |
| BD | 750718 | BUV661 anti-CD103 | 2E7 | 500 |
| BD | 747971 | BV480 anti-CD192 | 475301 | 200 |
| BD | 747298 | BV750 anti-CD183 | CXCR3-173 | 500 |
| BD | 624295 | BB660-P2 anti-CD95 | Jo2 | 1000 |
| BD | 742206 | BB700 anti-LAG3 | C9B7W | 100 |
| BD | 624296 | BB790-P anti-Ly-6C | AL-21 | 2000 |
| BioLegend | 313524 | BV421 anti-ICOS | C398.4A | 500 |
| BioLegend | 137220 | Pacific Blue anti-Helios | 22F6 | 250 |
| BioLegend | 103044 | BV510 anti-CD44 | IM7 | 1000 |
| BioLegend | 135231 | BV711 anti-PD-1 | 29F.1A12 | 5000 |
| BioLegend | 109862 | Spark Blue 550 anti-CD45.2 | 104 | 2000 |
| BioLegend | 104510 | PE-Cy5 anti-CD69 | H1.2F3 | 200 |
| eBioscience | 56-5698-82 | Alexa Fluor 700 anti-Ki67 | SolA15 | 1000 |
| BioLegend | 321224 | APC-Fire 750 anti-Integrin β7 | FIB504 | 200 |
| eBioscience | 78-5893-80 | Super Bright 780 anti-KLRG1 | 2F1 | 500 |
| BioLegend | 110718 | Alexa Fluor 488 anti-CD45.1 | A20 | 2000 |
| eBioscience | 46-3041-82 | PerCP-eFluor710 anti-CD304 | 3DS304M | 5000 |
| eBioscience | 12-5965-82 | PE anti-Nur77 | 12.14 | 250 |
| eBioscience | 17-0900-82 | APC anti-CD90.1 | HIS51 | 400 |

| | | | | |
|---|---|---|---|---|
| eBioscience | 606-5773-80 | Alexa Fluor 660 anti-Foxp3 | FJK-16s | 400 |
| eBioscience | 13-0451-85 | Biotin anti-CD45 | 30-F11 | 2μg/mouse |
| Beckman Coulter | C36628 | ViaKrome 808 | N/A | 1000 |

**Supplementary Table 5: Antibody information for Symphony mouse parabiosis panel.**

| Vendor | Catalogue # | Antibody | Clone | Dilution |
|---|---|---|---|---|
| BD | 740238 | BUV395 anti-CD103 | M290 | 1000 |
| BD | 612952 | BUV496 anti-CD4 | GK1.5 | 500 |
| BD | 564920 | BUV563 anti-CD45 | 30-F11 | 10000 |
| BD | 612833 | BUV737 anti-CD62L | MEL-14 | 1000 |
| BD | 564920 | BUV805 anti-CD8a | 53-6.7 | 1000 |
| BioLegend | 106312 | BV421 anti-CD152 | UC10-4B9 | 200 |
| BD | 566120 | BV480 anti-CD25 | PC61 | 1000 |
| BioLegend | 103037 | BV570 anti-CD44 | IM7 | 500 |
| BioLegend | 313538 | BV605 anti-ICOS | C398.4A | 500 |
| BioLegend | 100229 | BV650 anti-CD3 | 17A2 | 2000 |
| BioLegend | 135231 | BV711 anti-PD-1 | 29F.1A12 | 5000 |
| BD | 747332 | BV750 anti-CD19 | 1D3 | 2000 |
| BioLegend | 138429 | BV785 anti-KLRG1 | 2F1/KLRG1 | 500 |
| BioLegend | 110718 | Alexa Fluor 488 anti-CD45.1 | A20 | 2000 |
| eBioscience | 46-3041-82 | PerCP-eFluor710 anti-CD304 | 3DS304M | 5000 |
| Miltenyi | 130-098-596 | PE anti-T-bet | REA102 | 200 |
| Miltenyi | 130-112-636 | PE-Vio615 anti-Helios | REA829 | 500 |
| BioLegend | 104510 | PE-Cy5 anti-CD69 | H1.2F3 | 200 |
| BioLegend | 108702 | Purified anti-NK1.1 | PK136 | 10000 |
| AbCam | ab102899 | PE-Cy5.5 conjugation kit | N/A | N/A |
| eBioscience | 25-9335-80 | PE-Cy7 anti-ST2 | RMST2-2 | 200 |
| eBioscience | 17-5773-82 | APC anti-Foxp3 | FJK-16s | 200 |
| eBioscience | 56-5698-82 | Alexa Fluor 700 anti-Ki67 | SolA15 | 1000 |
| eBioscience | 65-0865-18 | Fixable Viability Dye eFluor780 | N/A | 2000 |

**Supplementary Figures**

**Supplementary Figure 1. Phenotypic unity for tissue-resident Tregs across multiple organs.** Wildtype mice, aged 12-20 weeks, were injected with intravenous anti-CD45 antibody label and assessed for flow cytometry. Tregs were purified from 48 organs: blood, thymus, spleen, bone marrow, cervical LN, submandibular LN, axillary LN, inguinal LN, popliteal LN, pancreatic LN, mediastinal LN, renal LN, aortic LN, jejunal LN, colonic LN, skin, lung, pancreas, salivary gland, liver, kidney, tongue, heart, muscle, white fat, brown fat, brain, female reproductive tract, urethra, testes, prostate, bladder, peritoneum, duodenal Peyer's Patches, jejunal Peyer's Patches, ileal Peyer's Patches, stomach IEL, duodenal IEL, Jejunal IEL, ileal IEL, cecal IEL, colonic IEL, stomach LPL, duodenal LPL, jejunal LPL, ileal LPL, cecal LPL, and colonic LPL. **A)** UMAP plots of high parameter flow analysis of gated tissue Tregs, built upon expression of GITR, TIM3, CCR3, TIGIT, Ly-6C, CCR9, CCR6, CXCR3, CCR2, KLRG1, ST2, CD62L, Neuropilin, CD69, CD44, CD25, CD95, TNFRII, ICOS, PD-1. **B)** Heatmap of phenotypic marker expression in Tregs from each tissue, clustered by similarity, for each marker.

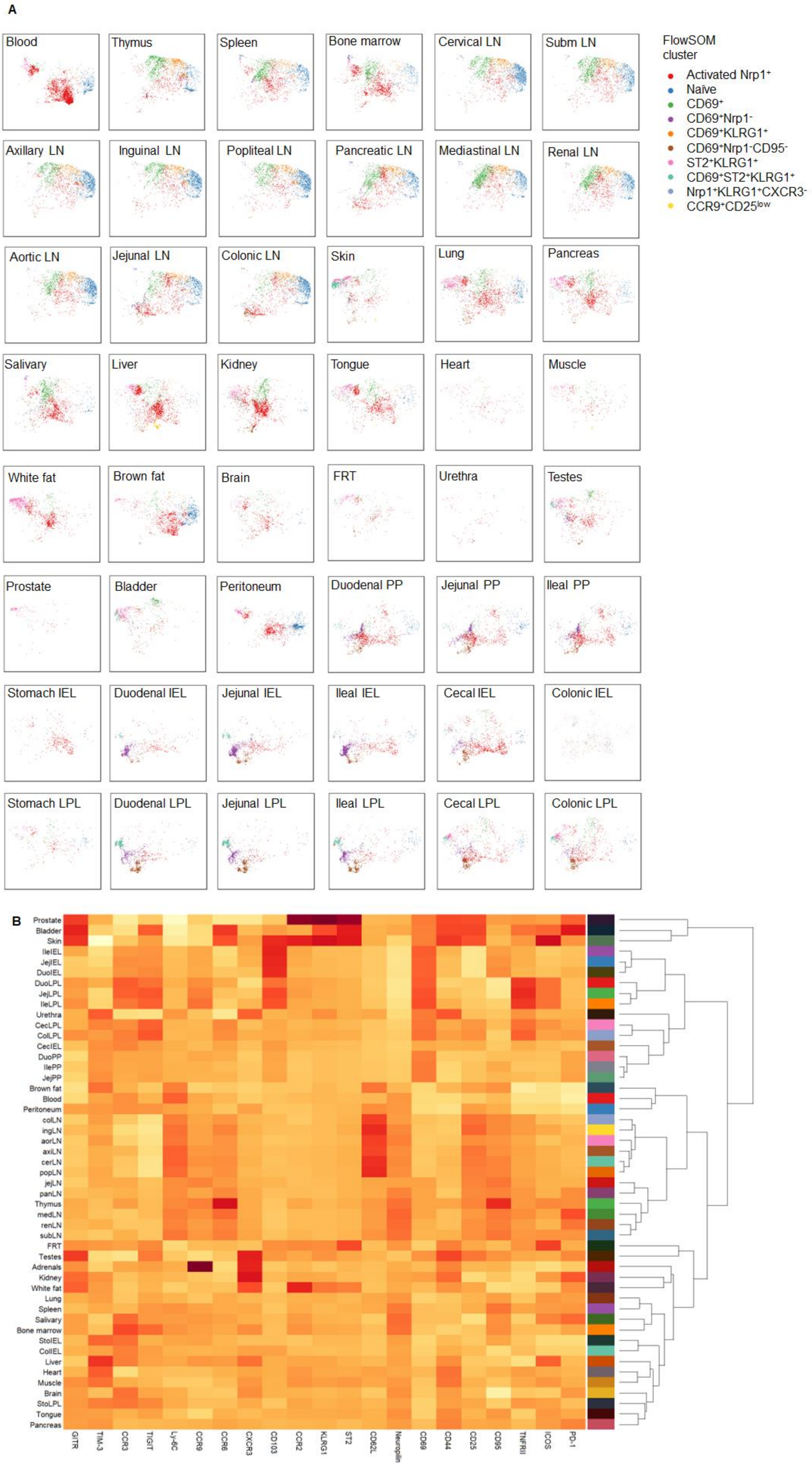
A
Blood
Thymus
Spleen
Bone marrow
Cervical LN
Subm LN
Axillary LN
Inguinal LN
Popliteal LN
Pancreatic LN
Mediastinal LN
Renal LN
Aortic LN
Jejunal LN
Colonic LN
Skin
Lung
Pancreas
Salivary
Liver
Kidney
Tongue
Heart
Muscle
White fat
Brown fat
Brain
FRT
Urethra
Testes
Prostate
Bladder
Peritoneum
Duodenal PP
Jejunal PP
Ileal PP
Stomach IEL
Duodenal IEL
Jejunal IEL
Ileal IEL
Cecal IEL
Colonic IEL
Stomach LPL
Duodenal LPL
Jejunal LPL
Ileal LPL
Cecal LPL
Colonic LPL
FlowSOM cluster
Activated Nrp1+
Naïve
CD69+
CD69+Nrp1-
CD69+KLRG1+
CD69+Nrp1-CD95-
ST2+KLRG1+
CD69+ST2+KLRG1+
Nrp1+KLRG1+CXCR3-
CCR9+CD25low
B
Prostate
Bladder
Skin
IleIEL
JejIEL
DuoIEL
DuoLPL
JejLPL
IleLPL
Urethra
CecLPL
ColLPL
CecIEL
DuoPP
IlePP
JejPP
Brown fat
Blood
Peritoneum
colLN
ingLN
aorLN
axiLN
cerLN
popLN
jejLN
panLN
Thymus
medLN
renLN
subLN
FRT
Testes
Adrenals
Kidney
White fat
Lung
Spleen
Salivary
Bone marrow
StoIEL
ColIEL
Liver
Heart
Muscle
Brain
StoLPL
Tongue
Pancreas
GITR
TIM-3
CCR3
TIGIT
Ly-6C
CCR9
CCR6
CXCR3
CD103
CCR2
KLRG1
ST2
CD62L
Neuropilin
CD69
CD44
CD25
CD95
TNFRII
ICOS
PD-1

**Supplementary Figure 2. Phenotypic signals of *in situ* differentiation defines tissue-Treg populations.** Wildtype mice, aged 16 weeks, were injected with intravenous anti-CD45-PE antibody label and perfused prior to tissue Treg isolation for flow cytometry analysis. **A)** Fraction of Tregs labeled by i.v. injection of anti-CD45-PE (n=4). **B)** Tissue Treg phenotypes were compared by tSNE and FlowSOM cluster overlay for blood vs liver Tregs, with liver Tregs separated into the vascular (CD45-PE$^{+}$) population, a potentially perivascular CD45-PE$^{mid}$ population and the tissue-embedded CD45-PE$^{-}$ populations. **C)** Heat map and **D)** UMAP for tissue Treg phenotypes were compared by tSNE and FlowSOM cluster overlay for blood vs combined lymphoid, non-lymphoid and gut-associated Tregs, with these Tregs separated into the vascular (CD45-PE$^{+}$) population, a potentially perivascular CD45-PE$^{low}$ population and the tissue-embedded CD45-PE$^{-}$ populations.

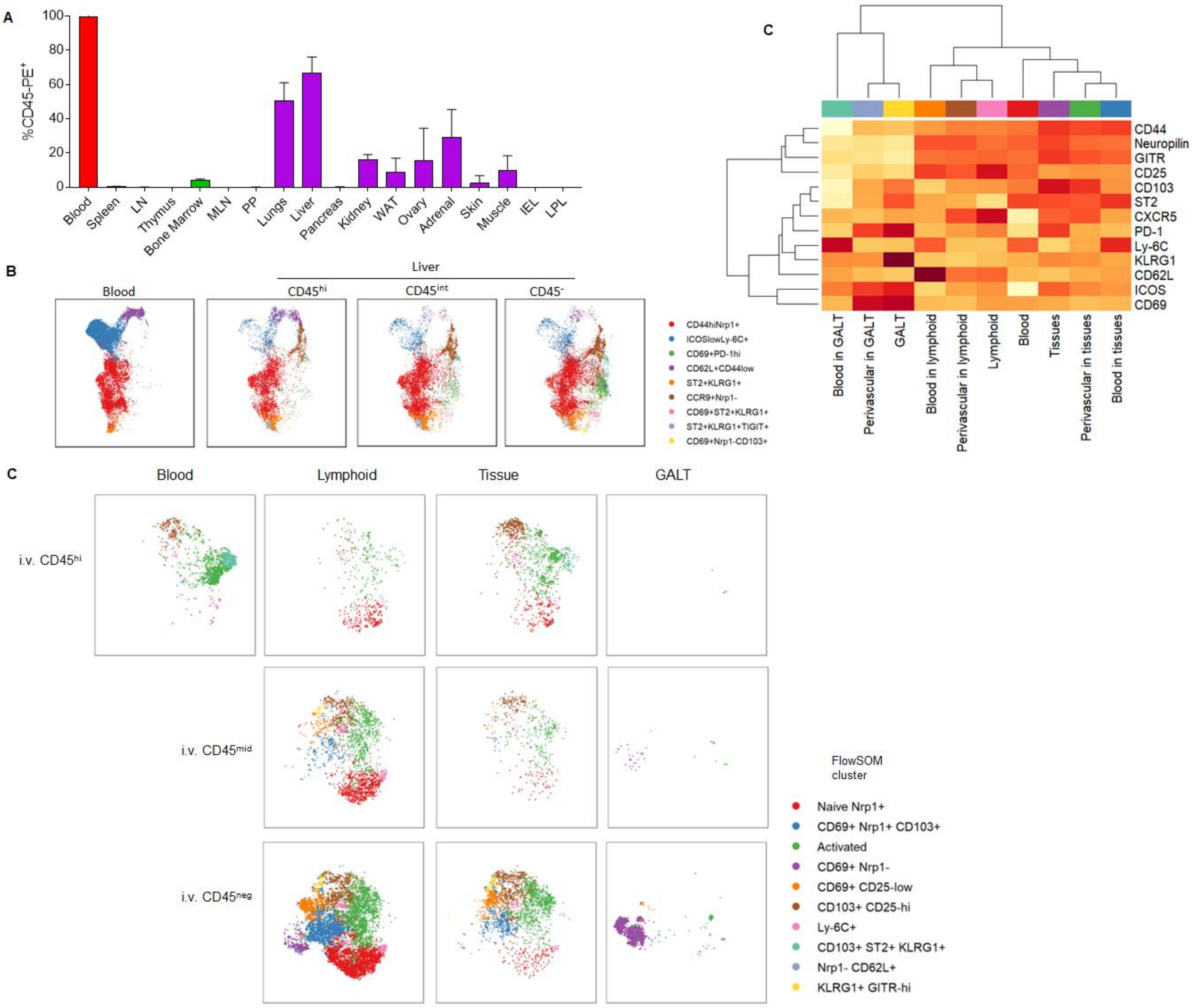

**Supplementary Figure 3. Tissue digestion protocols do not drive phenotypic divergence.** Blood, LN, skin and spleen were isolated for tissue Treg phenotyping by flow cytometry. Spleen samples were either given the standard (lymphoid tissue) mechanical disruption or (the non-lymphoid protocol of) enzymatic tissue digestion. **A)** Total leukocyte and **B)** Treg numbers for spleen samples prepared by mechanical or enzymatic digestion of the spleen. **C)** Frequency of Tregs among the isolated CD4 T cell population, for spleen samples prepared by mechanical or enzymatic digestion of the spleen. Statistical analysis by Šídák's multiple comparison test on 2-way ANOVA. **D)** Phenotypic comparison of mechanical and enzymatic splenic Tregs, using blood, lymphoid (cervical LN) and non-lymphoid (skin) out-groups. Displayed by PCA or **E)** UMAP, with **F)** quantified FlowSOM clusters. Statistical analysis by Tukey’s multiple comparison test on 2-way ANOVA. **G)** Heat map of the Treg expression of the markers used for PCA and UMAP analysis, GITR, TIM3, CCR3, TIGIT, Ly-6C, CCR9, CCR6, CXCR3, CCR2, KLRG1, ST2, CD62L, Neuropilin, CD69, CD44, CD25, CD95, TNFRII, ICOS, PD-1.

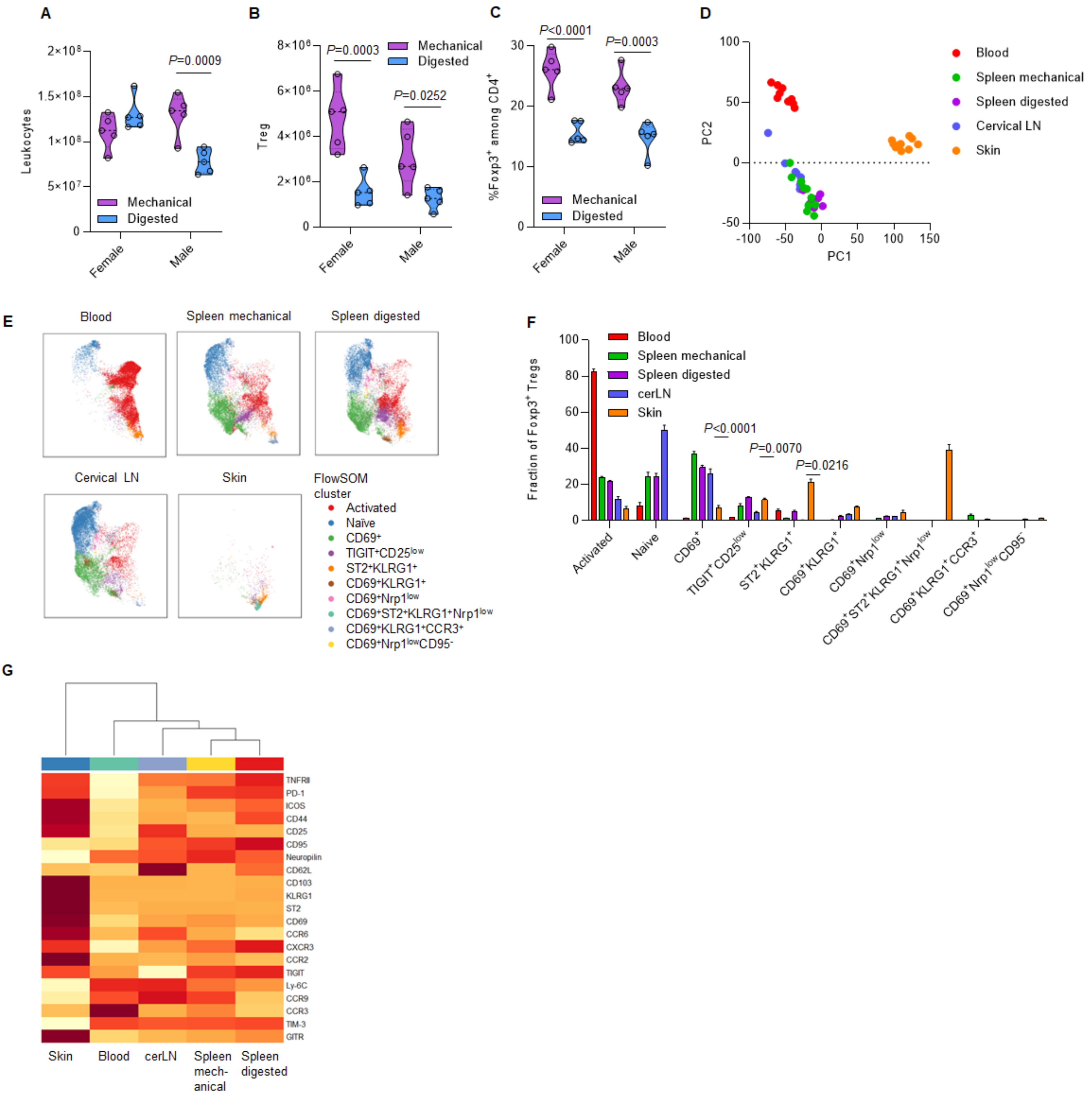
A
B
C
D
Leukocytes
2×10⁸
1.5×10⁸
1×10⁸
5×10⁷
0
P=0.0009
Mechanical
Digested
Female
Male
Treg
8×10⁶
6×10⁶
4×10⁶
2×10⁶
P=0.0003
P=0.0252
%Foxp3⁺ among CD4⁺
30
20
10
P<0.0001
P=0.0003
PC2
PC1
100
50
0
-50
-100
150
Blood
Spleen mechanical
Spleen digested
Cervical LN
Skin
E
Cervical LN
FlowSOM cluster
Activated
Naïve
CD69⁺
TIGIT⁺CD25^low
ST2⁺KLRG1⁺
CD69⁺KLRG1⁺
CD69⁺Nrp1^low
CD69⁺ST2⁺KLRG1⁺Nrp1^low
CD69⁺KLRG1⁺CCR3⁺
CD69⁺Nrp1^lowCD95⁻
F
Fraction of Foxp3⁺ Tregs
80
60
40
cerLN
P<0.0001
P=0.0070
P=0.0216
Activated
Naive
CD69⁺
TIGIT⁺CD25^low
ST2⁺KLRG1⁺
CD69⁺KLRG1⁺
CD69⁺Nrp1^low
CD69⁺ST2⁺KLRG1⁺Nrp1^low
CD69⁺KLRG1⁺CCR3⁺
CD69⁺Nrp1^lowCD95⁻
G
TNFRII
PD-1
ICOS
CD44
CD25
CD95
Neuropilin
CD62L
CD103
KLRG1
ST2
CD69
CCR6
CXCR3
CCR2
TIGIT
Ly-6C
CCR9
CCR3
TIM-3
GITR
Skin
Blood
cerLN
Spleen mech-anical
Spleen digested

**Supplementary Figure 4. Minor effects of sex on tissue Treg number and phenotype across multiple tissues.** Wildtype mice, aged 12-20 weeks, were injected with intravenous anti-CD45 antibody label. Major tissues were dissected and lymphocytes purified for flow cytometric analysis of Treg populations. **A)** Frequency of Tregs among $CD4^{+}$ $TCR\beta^{+}$ T cells, per organ, for female and male mice (n=5/group, representative of two independent experiments). No statistically significant differences by Šídák's multiple comparisons test on 2-way ANOVA. **B)** Absolute number of Tregs recorded per tissue, box-and-whisker plots showing 2.5-97.5% interval. Statistical analysis by by Šídák's multiple comparisons test on 2-way ANOVA. **C)** tSNE plots showing Treg phenotype in male or female Tregs, gated as viable $CD45^{+}CD3^{+}CD4^{+}Foxp3^{+}$ intravenously labeled $CD45^{-}$ and built on CCR2, CCR3, CCR6, CCR9, CD25, CD44, CD62L, CD69, CD95, CD103, CXCR3, GITR, ICOS, KLRG1, Ly-6c, Neuropilin, PD-1, ST2, TIGIT, Tim-3 and TNFRII for major tissue groups. **D)** tSNE plots showing comparisons of the most sexually dimorphic non-lymphoid tissues: reproductive tissue, white adipose tissue, kidney, salivary gland. P-values by KS test on crossentropy (tSNE-diff) for each pair.

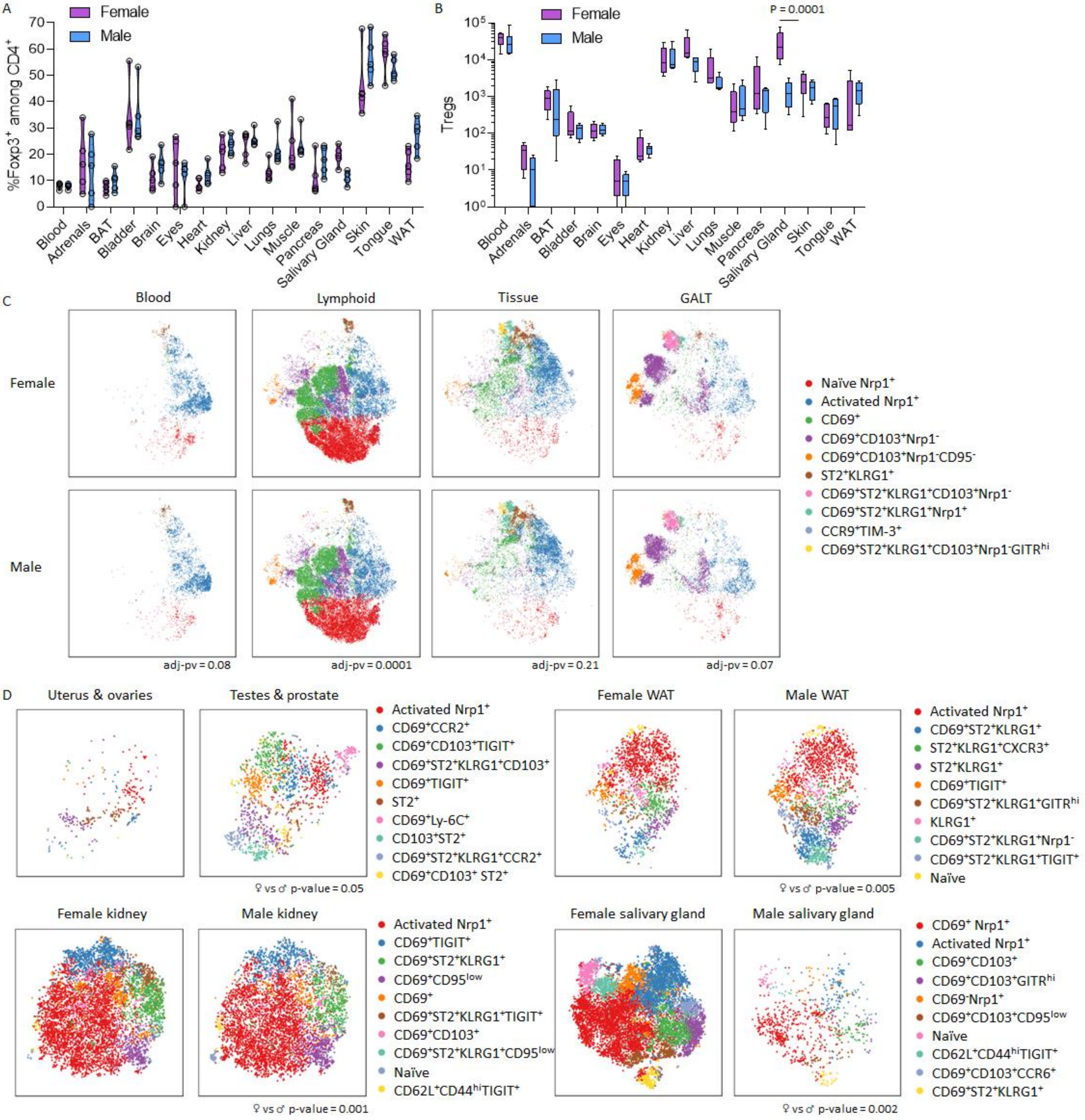
A
Female
Male
%Foxp3+ among CD4+
70
60
50
40
30
20
10
0
Blood
Adrenals
BAT
Bladder
Brain
Eyes
Heart
Kidney
Liver
Lungs
Muscle
Pancreas
Salivary Gland
Skin
Tongue
WAT
B
Female
Male
P = 0.0001
Tregs
10^5
10^4
10^3
10^2
10^1
10^0
Blood
Adrenals
BAT
Bladder
Brain
Eyes
Heart
Kidney
Liver
Lungs
Muscle
Pancreas
Salivary Gland
Skin
Tongue
WAT
C
Blood
Lymphoid
Tissue
GALT
Female
Male
adj-pv = 0.08
adj-pv = 0.0001
adj-pv = 0.21
adj-pv = 0.07
Naïve Nrp1+
Activated Nrp1+
CD69+
CD69+CD103+Nrp1-
CD69+CD103+Nrp1-CD95-
ST2+KLRG1+
CD69+ST2+KLRG1+CD103+Nrp1-
CD69+ST2+KLRG1+Nrp1+
CCR9+TIM-3+
CD69+ST2+KLRG1+CD103+Nrp1-GITRhi
D
Uterus & ovaries
Testes & prostate
♀ vs ♂ p-value = 0.05
Activated Nrp1+
CD69+CCR2+
CD69+CD103+TIGIT+
CD69+ST2+KLRG1+CD103+
CD69+TIGIT+
ST2+
CD69+Ly-6C+
CD103+ST2+
CD69+ST2+KLRG1+CCR2+
CD69+CD103+ ST2+
Female WAT
Male WAT
♀ vs ♂ p-value = 0.005
Activated Nrp1+
CD69+ST2+KLRG1+
ST2+KLRG1+CXCR3+
ST2+KLRG1+
CD69+TIGIT+
CD69+ST2+KLRG1+GITRhi
KLRG1+
CD69+ST2+KLRG1+Nrp1-
CD69+ST2+KLRG1+TIGIT+
Naïve
Female kidney
Male kidney
♀ vs ♂ p-value = 0.001
Activated Nrp1+
CD69+TIGIT+
CD69+ST2+KLRG1+
CD69+CD95low
CD69+
CD69+ST2+KLRG1+TIGIT+
CD69+CD103+
CD69+ST2+KLRG1+CD95low
Naïve
CD62L+CD44hiTIGIT+
Female salivary gland
Male salivary gland
♀ vs ♂ p-value = 0.002
CD69+ Nrp1+
Activated Nrp1+
CD69+CD103+
CD69+CD103+GITRhi
CD69-Nrp1+
CD69+CD103+CD95low
Naïve
CD62L+CD44hiTIGIT+
CD69+CD103+CCR6+
CD69+ST2+KLRG1+

**Supplementary Figure 5. Impact of age and microbiome on tissue-resident Tregs.** Mice were perfused and assessed by flow cytometry for tissue Treg number and phenotype at 8, 12, 20, 30, 52 and 100 weeks of age (n=5-8). **A)** Fold change in Treg cell number in the blood, lymphoid tissues (thymus, spleen, LN, bone-marrow, mLN, Peyer's patches), non-lymphoid tissues (skin, muscle, female reproductive tract, lungs, pancreas, brain, white adipose tissue, liver, kidney, adrenals) and gut-associated tissues (IEL and LPL). Tissues displayed on multiple graphs according to the scale of the fold-change. **B)** Frequency of Tregs within CD4 T cell population per tissue and age group, n.d. = not done. **C)** tSNE of Treg phenotype among blood, lymphoid tissues, non-lymphoid tissues and gut-associated tissues at each age, gated on viable $CD45^{+}CD3^{+}CD4^{+}Foxp3^{+}$ and built on CD103, CD62L, CTLA-4, CD25, CD44, ICOS, PD-1, KLRG1, Neuropilin, T-bet, Helios, CD69, ST2 and Ki67. **D)** SPF-housed mice, gnotobiotic (germ-free) mice and wild-exposed co-housed mice (n=9, 6, 12) perfused and assessed by flow cytometry for tissue Treg phenotype at 8-12 weeks of age, displayed as PCA plots of expression for the same markers as in C. Graphs separated into blood, lymphoid tissues (thymus, spleen, LN, bone-marrow, mLN, Peyer's patches), non-lymphoid tissues (skin, muscle, female reproductive tract, lungs, pancreas, brain, white adipose tissue, liver, kidney, adrenals) and gut-associated tissues (IEL and LPL).

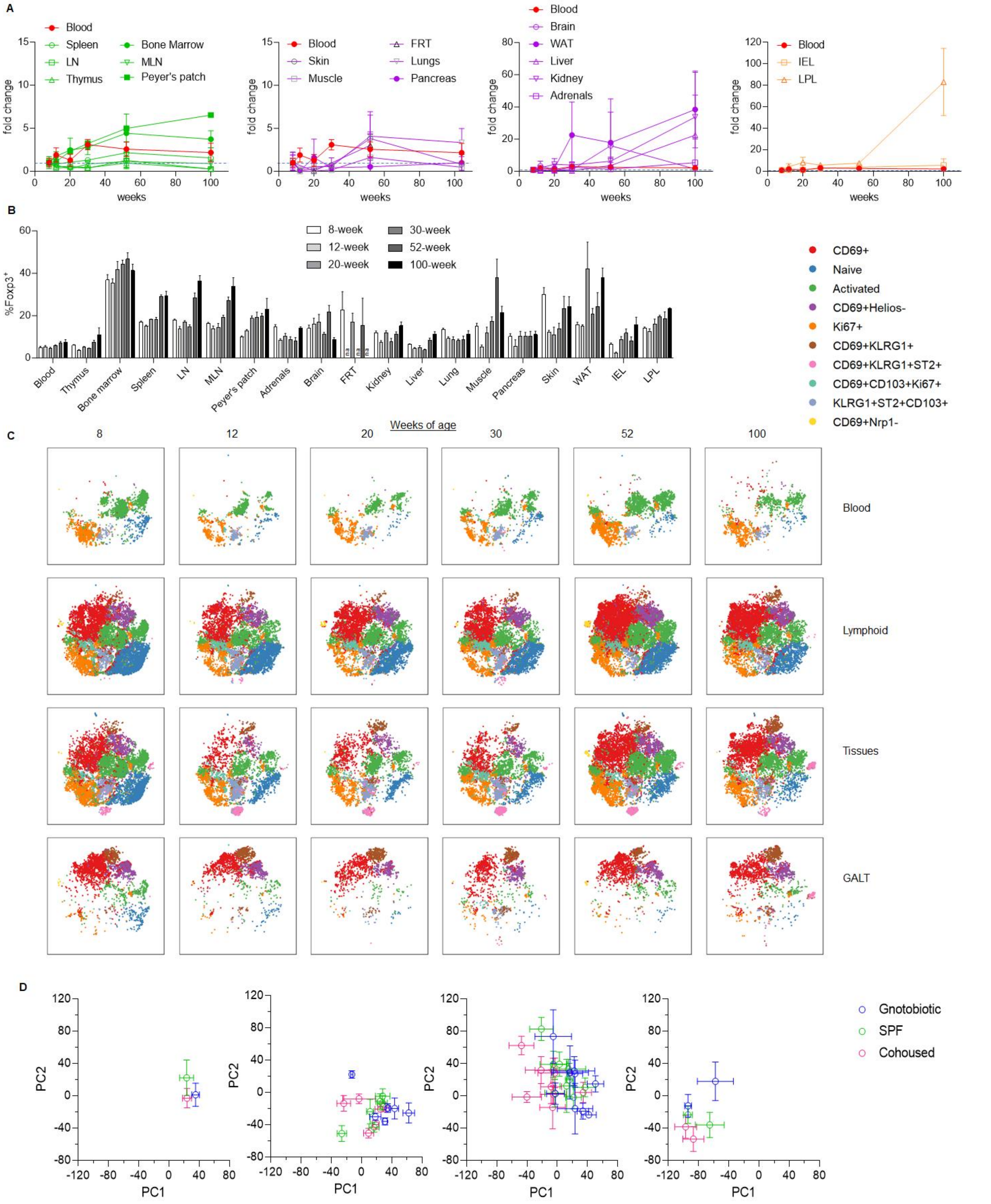
A
Blood
Spleen
LN
Thymus
Bone Marrow
MLN
Peyer's patch
fold change
weeks
Skin
Muscle
FRT
Lungs
Pancreas
Brain
WAT
Liver
Kidney
Adrenals
IEL
LPL
B
%Foxp3+
8-week
12-week
20-week
30-week
52-week
100-week
Blood
Thymus
Bone marrow
Spleen
LN
MLN
Peyer's patch
Adrenals
Brain
FRT
Kidney
Liver
Lung
Muscle
Pancreas
Skin
WAT
IEL
LPL
CD69+
Naive
Activated
CD69+Helios-
Ki67+
CD69+KLRG1+
CD69+KLRG1+ST2+
CD69+CD103+Ki67+
KLRG1+ST2+CD103+
CD69+Nrp1-
Weeks of age
C
8
12
20
30
52
100
Blood
Lymphoid
Tissues
GALT
D
PC1
PC2
Gnotobiotic
SPF
Cohoused

**Supplementary Figure 6**. **CD69 deficiency globally reduce Treg numbers without specific impact on tissue Treg differentiation.** Mixed bone-marrow chimeras were generated with 50% CD45.1 wildtype bone-marrow and 50% CD45.2 *Cd69*$^{-/-}$ bone-marrow, transplanted into Rag-deficient recipients. Recipient mice were injected with intravenous anti-CD45 antibody label and assessed at 10 weeks post-transplantation by flow cytometry, allowing comparative assessment of the CD45.1 wt and the CD45.2 KO Tregs. **A)** Frequency of Tregs among WT or *Cd69*$^{-/-}$ CD4$^{+}$ T cells, across the tissue set assessed. **B)** Absolute numbers of WT and *Cd69*$^{-/-}$ Tregs. Statistical analysis by Šídák's multiple comparisons test on 2-way ANOVA. **C)** Frequency of Tregs when normalized to blood (100 being equal). **D)** tSNE plot of flow cytometry phenotype, for WT and *Cd69*$^{-/-}$ Tregs, based on the tissue grouping of blood, lymphoid tissues, non-lymphoid tissues and gut-associated tissues, built on the markers T-bet, IRF4, Ki67, CCR9, CD96, Ly-6C, CD103, CTLA-4, CD62L, CCR2, CD44, ICOS, RORγT, PD-1, CXCR3, KLRG1, CD86, CD25, ST2, GATA-3, Neuropilin and Helios. Statistical analysis by Kolmogorov-Smirnov test (tSNE-diff) comparing WT vs. *Cd69*$^{-/-}$ Tregs. **E)** FlowSOM cluster distribution. Statistical analysis by Tukey's multiple comparisons test on 2-way ANOVA.

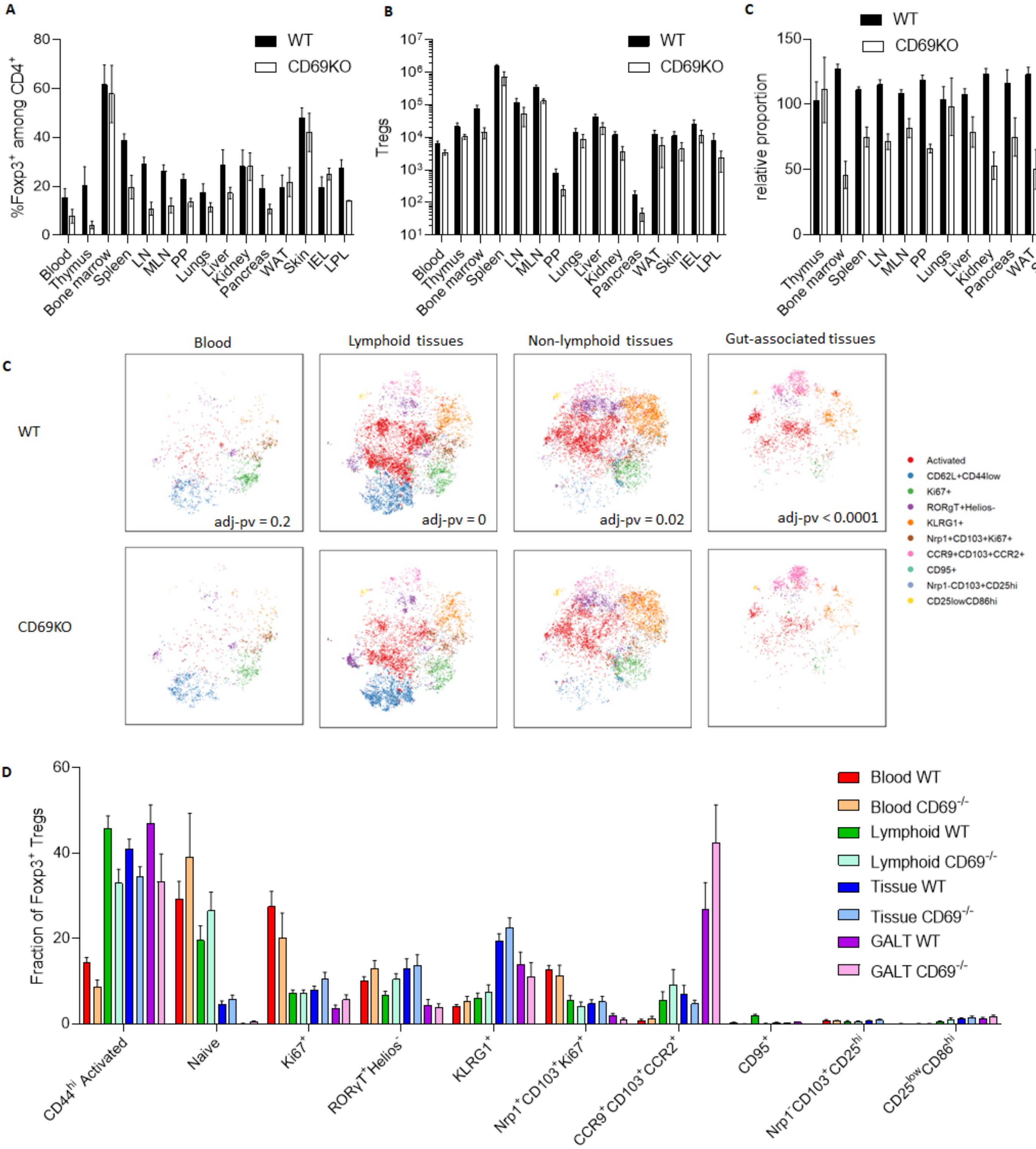
A
%Foxp3+ among CD4+
WT
CD69KO
Blood
Thymus
Bone marrow
Spleen
LN
MLN
PP
Lungs
Liver
Kidney
Pancreas
WAT
Skin
IEL
LPL
B
Tregs
C
relative proportion
Blood
Lymphoid tissues
Non-lymphoid tissues
Gut-associated tissues
adj-pv = 0.2
adj-pv = 0
adj-pv = 0.02
adj-pv < 0.0001
Activated
CD62L+CD44low
Ki67+
RORgT+Helios-
KLRG1+
Nrp1+CD103+Ki67+
CCR9+CD103+CCR2+
CD95+
Nrp1-CD103+CD25hi
CD25lowCD86hi
D
Fraction of Foxp3+ Tregs
Blood WT
Blood CD69-/-
Lymphoid WT
Lymphoid CD69-/-
Tissue WT
Tissue CD69-/-
GALT WT
GALT CD69-/-
CD44hi Activated
Naive
Ki67+
RORγT+Helios-
KLRG1+
Nrp1+CD103+Ki67+
CCR9+CD103+CCR2+
CD95+
Nrp1-CD103+CD25hi
CD25lowCD86hi

**Supplementary Figure 7**. **ST2 deficiency does not alter Treg fitness or differentiation in tissues.** Mixed bone-marrow chimeras were generated with 50% CD45.1 wildtype bone-marrow and 50% CD45.2 *Il1rl1* $^{-/-}$ bone-marrow, transplanted into CD45.1/2 recipients. Recipient mice were assessed at 8 weeks post-transplantation by flow cytometry with intravenous anti-CD45 antibody label, allowing comparative assessment of the CD45.1 wt and the CD45.2 KO Tregs. **A)** Frequency of Tregs among WT or *Il1rl1* $^{-/-}$ CD4$^{+}$ T cells, across the tissue set assessed. **B)** Absolute numbers of WT and *Il1rl1* $^{-/-}$ Tregs. Statistical analysis by Šídák's multiple comparisons test on 2-way ANOVA. **C)** tSNE plot of flow cytometry phenotype, for WT and *Il1rl1* $^{-/-}$ Tregs, based on the tissue grouping of blood, lymphoid tissues, non-lymphoid tissues and gut-associated tissues, built on the markers FR4, CTLA-4, CD103, CD62L, GITR, Helios, CXCR3, CD44, CD25, PD-1, RORγT, CD127, ICOS, CCR6, CCR9, CCR7, Neuropilin, Ly-6c, Blimp-1, GATA-3, CD69, KLRG1, T-bet, CD38, IRF4 and Ki67. P-value by KS test with Holm correction on crossentropy (tSNE-diff) comparing WT vs. *Il1rl1* $^{-/-}$ Tregs. **D)** FlowSOM cluster distribution. Statistical analysis by Tukey's multiple comparisons test on 2-way ANOVA.

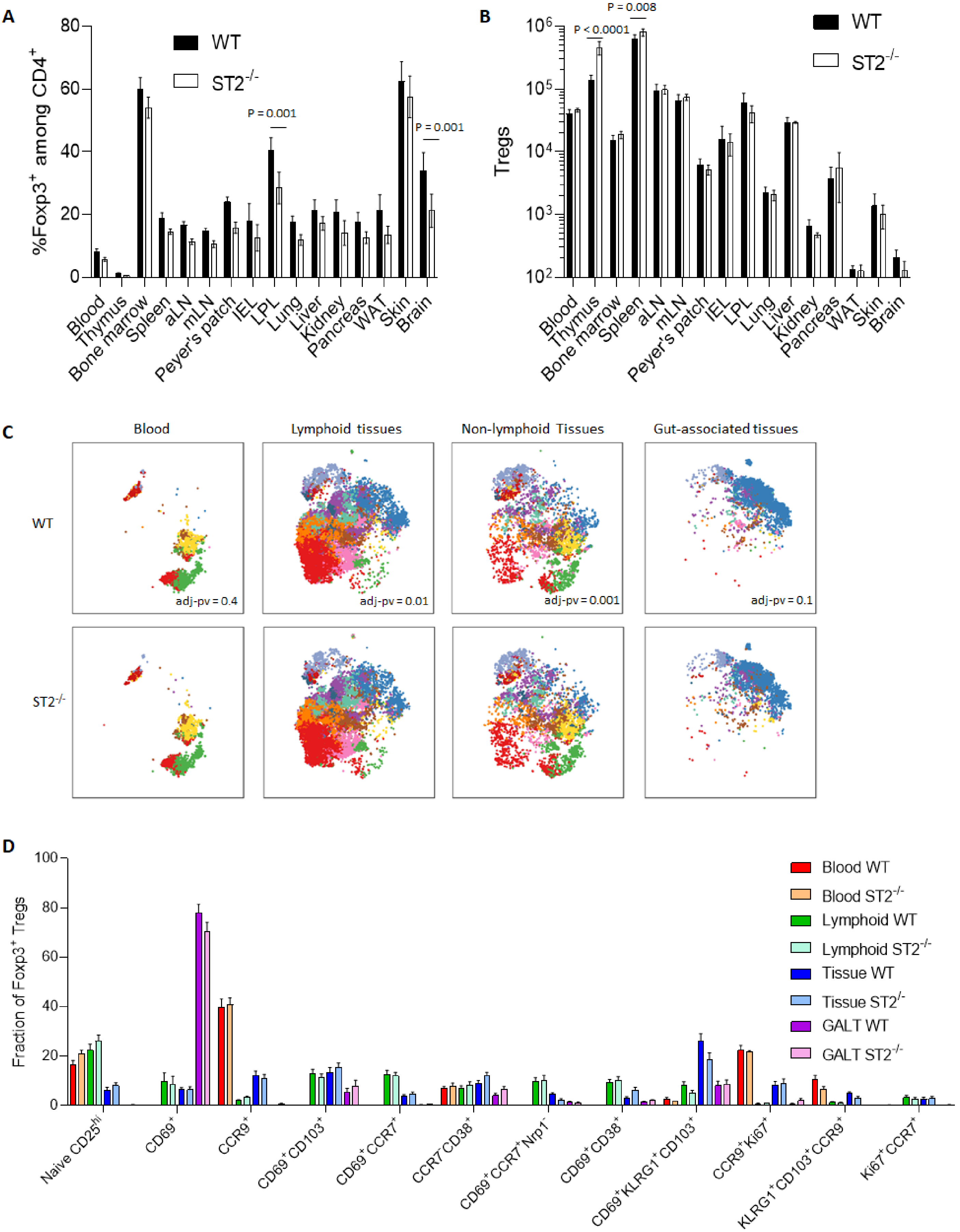
A
%Foxp3+ among CD4+
WT
ST2-/-
P = 0.001
P = 0.001
Blood
Thymus
Bone marrow
Spleen
aLN
mLN
Peyer's patch
IEL
LPL
Lung
Liver
Kidney
Pancreas
WAT
Skin
Brain
B
Tregs
P < 0.0001
P = 0.008
WT
ST2-/-
C
Blood
Lymphoid tissues
Non-lymphoid Tissues
Gut-associated tissues
WT
ST2-/-
adj-pv = 0.4
adj-pv = 0.01
adj-pv = 0.001
adj-pv = 0.1
D
Fraction of Foxp3+ Tregs
Blood WT
Blood ST2-/-
Lymphoid WT
Lymphoid ST2-/-
Tissue WT
Tissue ST2-/-
GALT WT
GALT ST2-/-
Naive CD25hi
CD69+
CCR9+
CD69+CD103+
CD69+CCR7+
CCR7-CD38+
CD69+CCR7+Nrp1-
CD69+CD38+
CD69+KLRG1+CD103+
CCR9+Ki67+
KLRG1+CD103+CCR9+
Ki67+CCR7+

**Supplementary Figure 8. CD11a-deficiency impedes Treg tissue entry in a pan-tissue manner.** Mixed bone-marrow chimeras were generated with 50% CD45.1 wildtype bone-marrow and 50% CD45.2 *Itgal*$^{-/-}$ bone-marrow, transplanted into Rag-deficient recipients. Recipient mice were assessed at 8 weeks post-transplantation by flow cytometry, allowing comparative assessment of the CD45.1 wt and the CD45.2 KO Tregs. **A)** Frequency of Tregs among WT or *Itgal*$^{-/-}$ CD4$^{+}$ T cells, across the tissue set assessed. **B)** Absolute numbers of WT and *Itgal*$^{-/-}$ Tregs. Statistical analysis by Šídák's multiple comparisons test on 2-way ANOVA. **C)** Frequency of Tregs when normalized to blood (100 being equal). **D)** UMAP plot of flow cytometry phenotype, for WT and *Itgal*$^{-/-}$ Tregs, based on the tissue grouping of blood, lymphoid tissues, non-lymphoid tissues and gut-associated tissues, built on the markers CD103, CTLA-4, CD62L, Helios, CCR2, CD44, ICOS, RORγT, PD-1, CXCR3, KLRG1, CCR9, CD95, Neuropilin, Ly-6c, TNFRII, T-bet, GATA-3, CD69, CD25, ST2, IRF4 and Ki67. Statistical analysis by Kolmogorov-Smirnov test (UMAP-diff) comparing WT vs. *Itgal*$^{/-}$ Tregs. **E)** FlowSOM cluster distribution. Statistical analysis by Tukey’s multiple comparisons test on 2-way ANOVA.

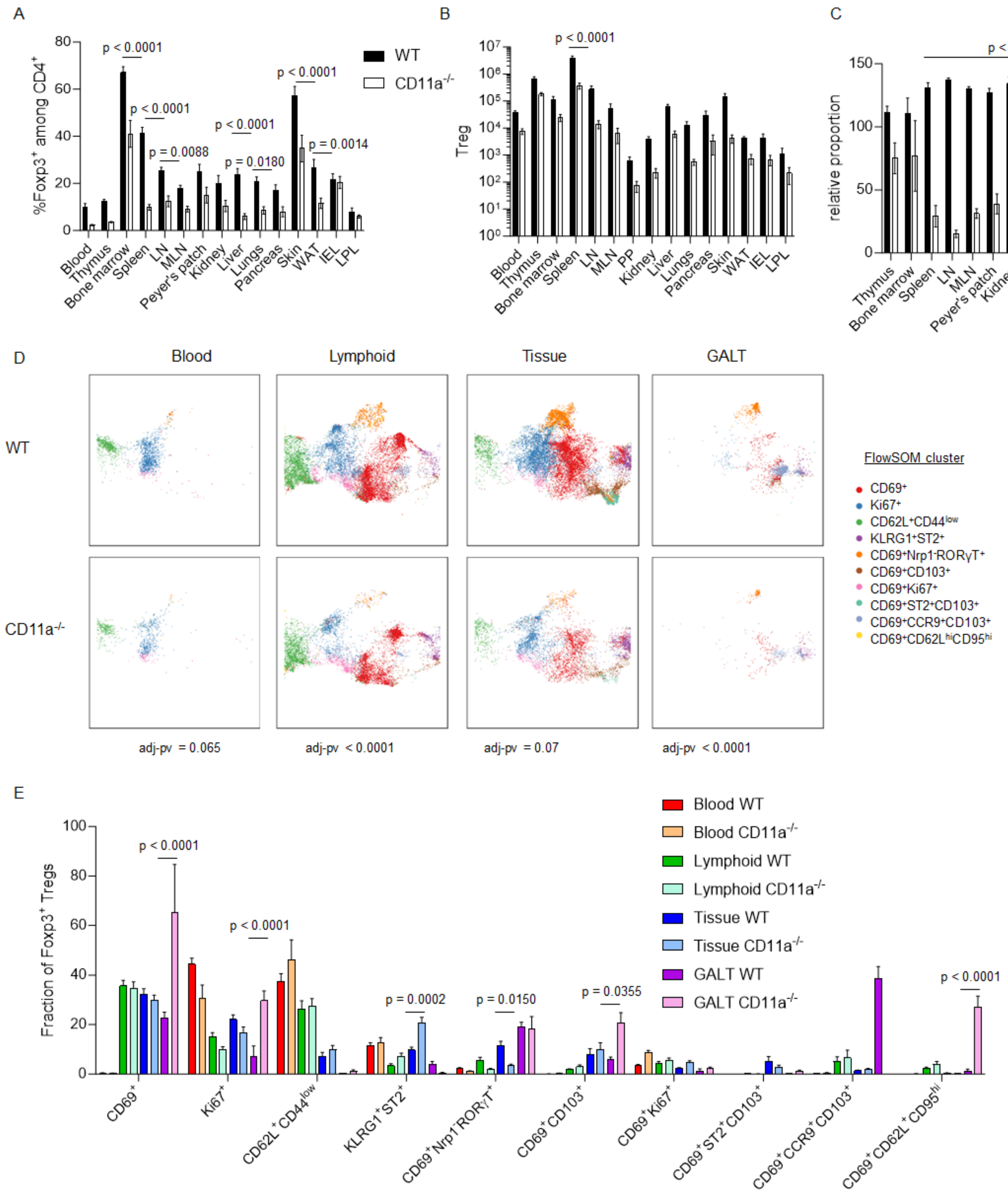

A
%Foxp3+ among CD4+
WT
CD11a-/-
p < 0.0001
p = 0.0088
p = 0.0180
p = 0.0014
Blood Thymus Bone marrow Spleen LN MLN Peyer's patch Kidney Liver Lungs Pancreas Skin WAT IEL LPL
B
Treg
p < 0.0001
Blood Thymus Bone marrow Spleen LN MLN PP Kidney Liver Lungs Pancreas Skin WAT IEL LPL
C
relative proportion
p < 0.0001
Thymus Bone marrow Spleen LN MLN Peyer's patch Kidney Liver Lungs Pancreas Skin WAT IEL LPL
D
Blood
Lymphoid
Tissue
GALT
WT
CD11a-/-
FlowSOM cluster
CD69+
Ki67+
CD62L+CD44low
KLRG1+ST2+
CD69+Nrp1-RORγT+
CD69+CD103+
CD69+Ki67+
CD69+ST2+CD103+
CD69+CCR9+CD103+
CD69+CD62LhiCD95hi
adj-pv = 0.065
adj-pv < 0.0001
adj-pv = 0.07
adj-pv < 0.0001
E
Fraction of Foxp3+ Tregs
Blood WT
Blood CD11a-/-
Lymphoid WT
Lymphoid CD11a-/-
Tissue WT
Tissue CD11a-/-
GALT WT
GALT CD11a-/-
p < 0.0001
p = 0.0002
p = 0.0150
p = 0.0355
CD69+
Ki67+
CD62L+CD44low
KLRG1+ST2+
CD69+Nrp1-RORγT+
CD69+CD103+
CD69+Ki67+
CD69+ST2+CD103+
CD69+CCR9+CD103+
CD69+CD62L+CD95hi

**Supplementary Figure 9**. **KLRG1 deficiency does not alter Treg fitness or differentiation in tissues.** Mixed bone-marrow chimeras were generated with 50% CD45.1 wildtype bone-marrow and 50% CD45.2 *Klrg1*$^{-/-}$ bone-marrow, transplanted into CD45.1/2 recipients. Recipient mice were assessed at 9 weeks post-transplantation by flow cytometry with intravenous CD45 labeling, allowing comparative assessment of the CD45.1 wt and the CD45.2 KO Tregs. **A)** Frequency of Tregs among WT or *Klrg1*$^{-/-}$ CD4$^{+}$ T cells, across the tissue set assessed. **B)** Absolute numbers of WT and *Klrg1*$^{-/-}$ Tregs. Statistical analysis by Šídák's multiple comparisons test on 2-way ANOVA. **C)** Frequency of Tregs when normalized to blood (100 being equal). **D)** tSNE plot of flow cytometry phenotype, for WT and *Klrg1*$^{-/-}$ Tregs, based on the tissue grouping of blood, lymphoid tissues, non-lymphoid tissues and gut-associated tissues, built on the markers CTLA-4, CD103, CD62L, GITR, Helios, CXCR3, CD44, PD-1, RORγT, CD127, LAG3, Tim-3, Neuropilin, Ly-6C, Blimp-1, ST2, CD25, CD69, CD137, CD38, T-bet, Ki67, CCR8 and CCR7. Statistical analysis by Kolmogorov-Smirnov test with Holm correction (tSNE-diff) comparing WT vs. *Klrg1*$^{-/-}$ Tregs. **E)** FlowSOM cluster distribution. Statistical analysis by Tukey’s multiple comparisons test on 2-way ANOVA.

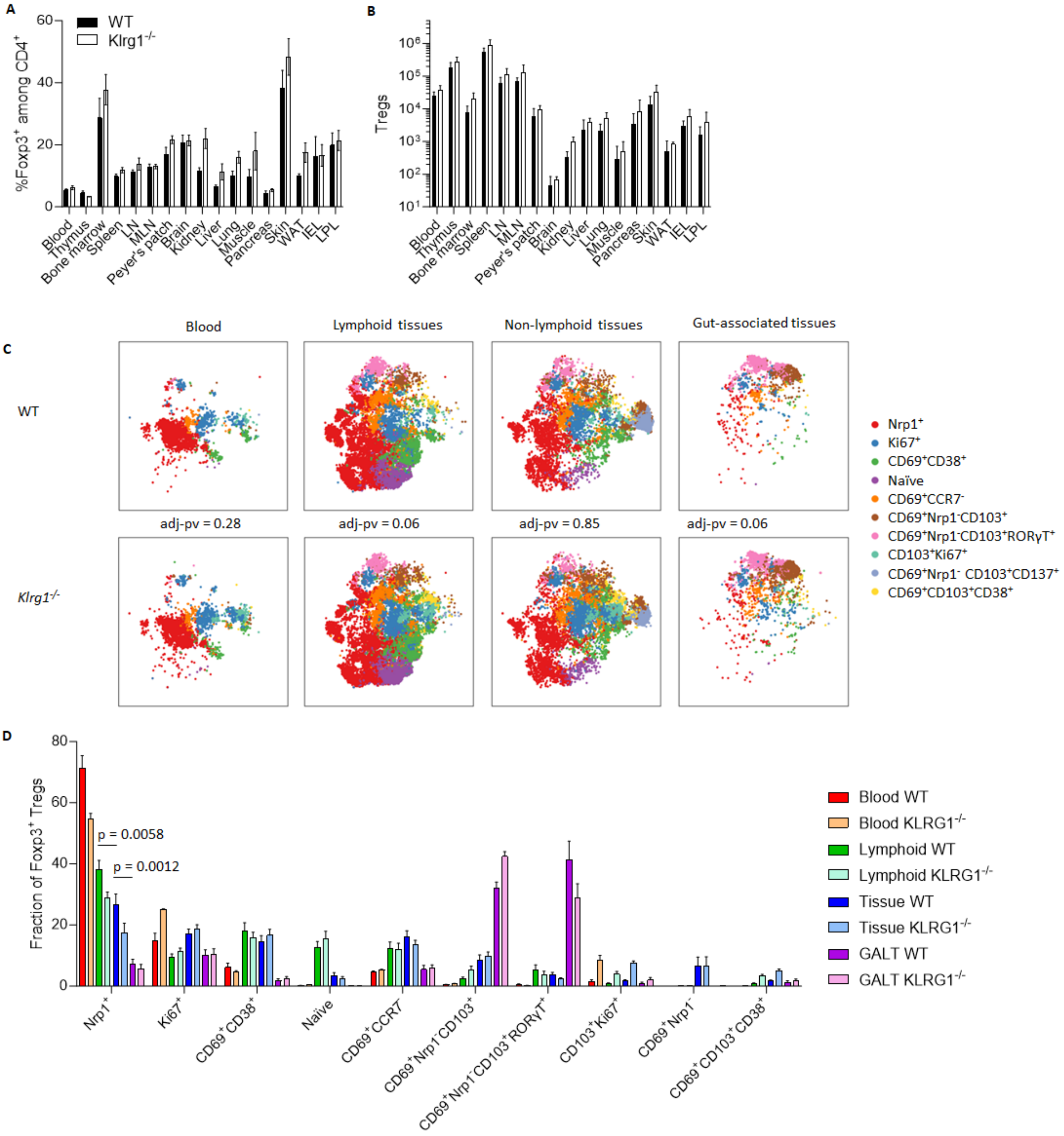
A
%Foxp3+ among CD4+
WT
Klrg1-/-
B
Tregs
Blood
Thymus
Bone marrow
Spleen
LN
MLN
Peyer's patch
Brain
Kidney
Liver
Lung
Muscle
Pancreas
Skin
WAT
IEL
LPL
C
Blood
Lymphoid tissues
Non-lymphoid tissues
Gut-associated tissues
WT
Klrg1-/-
adj-pv = 0.28
adj-pv = 0.06
adj-pv = 0.85
adj-pv = 0.06
Nrp1+
Ki67+
CD69+CD38+
Naïve
CD69+CCR7-
CD69+Nrp1-CD103+
CD69+Nrp1-CD103+RORγT+
CD103+Ki67+
CD69+Nrp1- CD103+CD137+
CD69+CD103+CD38+
D
Fraction of Foxp3+ Tregs
p = 0.0058
p = 0.0012
Blood WT
Blood KLRG1-/-
Lymphoid WT
Lymphoid KLRG1-/-
Tissue WT
Tissue KLRG1-/-
GALT WT
GALT KLRG1-/-
Nrp1+
Ki67+
CD69+CD38+
Naïve
CD69+CCR7-
CD69+Nrp1-CD103+
CD69+Nrp1-CD103+RORγT+
CD103+Ki67+
CD69+Nrp1-
CD69+CD103+CD38+

**Supplementary Figure 10. Loss of Blimp1 disfavors Tregs except in the gut.** Mixed bone-marrow chimeras were generated with 50% CD45.1 wildtype bone-marrow and 50% CD45.2 CD4-Cre *Prdm1*$^{fl/fl}$ (Blimp1-deficient) bone-marrow, transplanted into CD45.1/2 recipients. Recipient mice were assessed at 11 weeks post-transplantation by flow cytometry with intravenous CD45 labeling, allowing comparative assessment of the CD45.1 wt and the CD45.2 KO Tregs. **A)** Frequency of Tregs among WT or Blimp1-deficient CD4$^+$ T cells, across the tissue set assessed. **B)** Absolute numbers of WT and Blimp1-deficient Tregs. Statistical analysis by Šídák's multiple comparisons test on 2-way ANOVA. **C)** tSNE plot of flow cytometry phenotype, for WT and Blimp1-deficient Tregs, based on the tissue grouping of blood, lymphoid tissues, non-lymphoid tissues, gut-associated tissues, and the skin, built on the markers CTLA-4, CD103, CD62L, Helios, CCR2, CD44, CD25, RORγT, PD-1, CXCR3, KLRG1, CCR9, CD95, CCR7, Neuropilin, Ly-6c, Nur77, CD69, ST2, T-bet and Ki67. P-value by KS test with Holm correction on crossentropy (tSNE-diff) comparing WT vs. Blimp1-deficient Tregs. **E)** FlowSOM cluster distribution. P-values by Tukey’s multiple comparisons test on 2-way ANOVA.

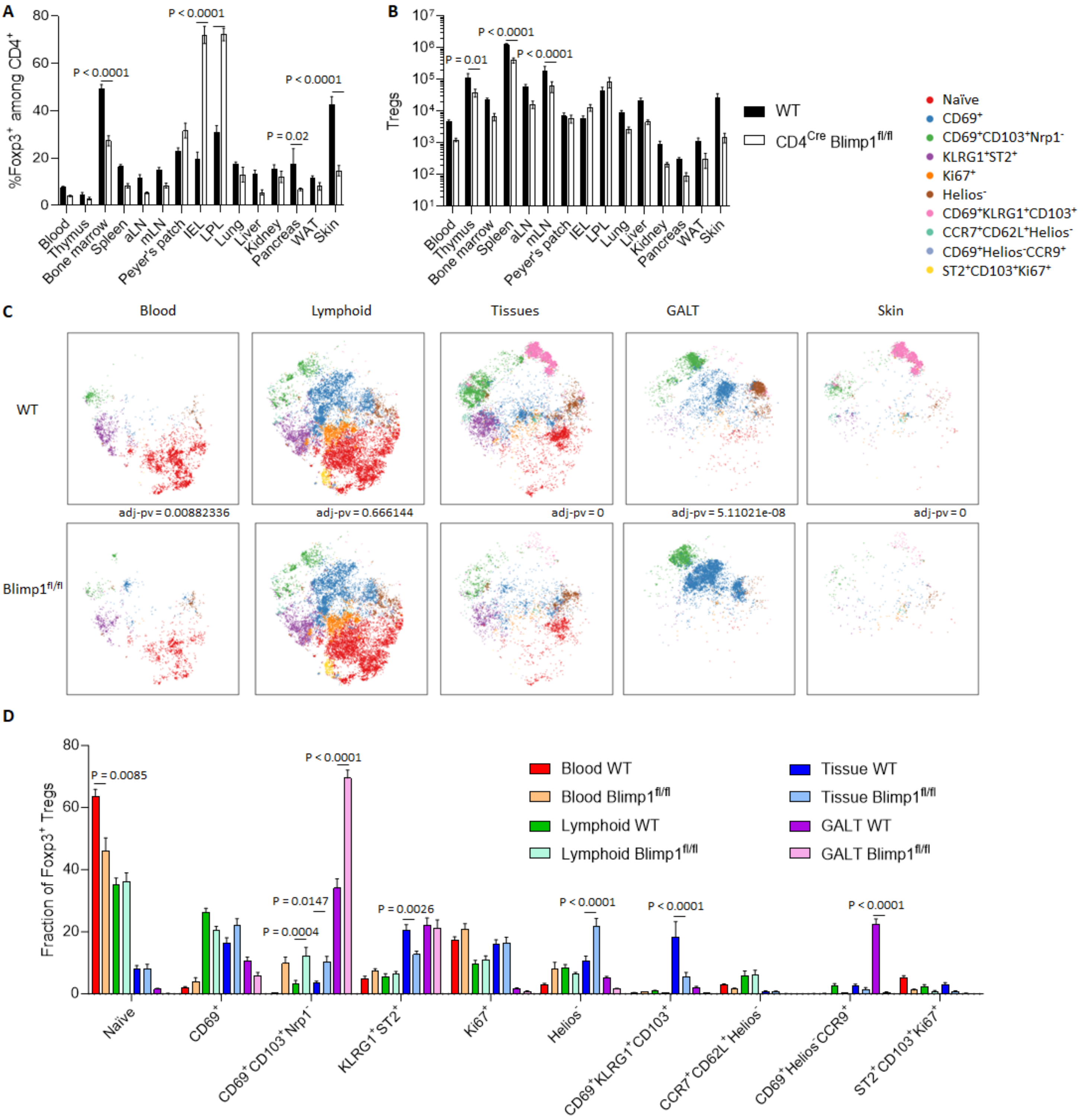
A
%Foxp3+ among CD4+
P < 0.0001
P < 0.0001
P = 0.02
P < 0.0001
Blood
Thymus
Bone marrow
Spleen
aLN
mLN
Peyer's patch
IEL
LPL
Lung
Liver
Kidney
Pancreas
WAT
Skin
B
Tregs
P = 0.01
P < 0.0001
P < 0.0001
WT
CD4Cre Blimp1fl/fl
Naïve
CD69+
CD69+CD103+Nrp1-
KLRG1+ST2+
Ki67+
Helios-
CD69+KLRG1+CD103+
CCR7+CD62L+Helios-
CD69+Helios-CCR9+
ST2+CD103+Ki67+
C
Blood
Lymphoid
Tissues
GALT
Skin
WT
adj-pv = 0.00882336
adj-pv = 0.666144
adj-pv = 0
adj-pv = 5.11021e-08
adj-pv = 0
Blimp1fl/fl
D
Fraction of Foxp3+ Tregs
P = 0.0085
P < 0.0001
P = 0.0147
P = 0.0004
P = 0.0026
P < 0.0001
P < 0.0001
P < 0.0001
Blood WT
Blood Blimp1fl/fl
Lymphoid WT
Lymphoid Blimp1fl/fl
Tissue WT
Tissue Blimp1fl/fl
GALT WT
GALT Blimp1fl/fl
Naïve
CD69+
CD69+CD103+Nrp1-
KLRG1+ST2+
Ki67+
Helios-
CD69+KLRG1+CD103+
CCR7+CD62L+Helios-
CD69+Helios-CCR9+
ST2+CD103+Ki67+

**Supplementary Figure 11**. **S1PR2 deficiency does not alter Treg fitness or differentiation in tissues.** Mixed bone-marrow chimeras were generated with 50% CD45.1 wildtype bone-marrow and 50% CD45.2 *S1pr2*$^{-/-}$ bone-marrow, transplanted into Rag-deficient recipients. Recipient mice were assessed at 8 weeks post-transplantation by flow cytometry with intravenous CD45 labeling, allowing comparative assessment of the CD45.1 wt and the CD45.2 KO Tregs. **A)** Frequency of Tregs among WT or *S1pr2*$^{-/-}$ CD4$^{+}$ T cells, across the tissue set assessed. **B)** Absolute numbers of WT and *S1pr2*$^{-/-}$ Tregs. Statistical analysis by Šídák's multiple comparisons test on 2-way ANOVA. **C)** Frequency of Tregs when normalized to blood (100 being equal). **D)** UMAP plot of flow cytometry phenotype, for WT and *S1pr2*$^{-/-}$ Tregs, based on the tissue grouping of blood, lymphoid tissues, non-lymphoid tissues and gut-associated tissues, built on the markers CD103, CTLA-4, CD62L, Helios, CCR2, CD44, ICOS, RORγT, PD-1, CXCR3, KLRG1, CCR9, CD95, Neuropilin, Ly-6c, TNFRII, T-bet, GATA-3, CD69, CD25, ST2, IRF4 and Ki67. Statistical analysis by Kolmogorov-Smirnov test (UMAP-diff) comparing WT vs. *S1pr2*$^{-/-}$ Tregs. **E)** FlowSOM cluster distribution. Statistical analysis by Tukey’s multiple comparisons test on 2-way ANOVA.

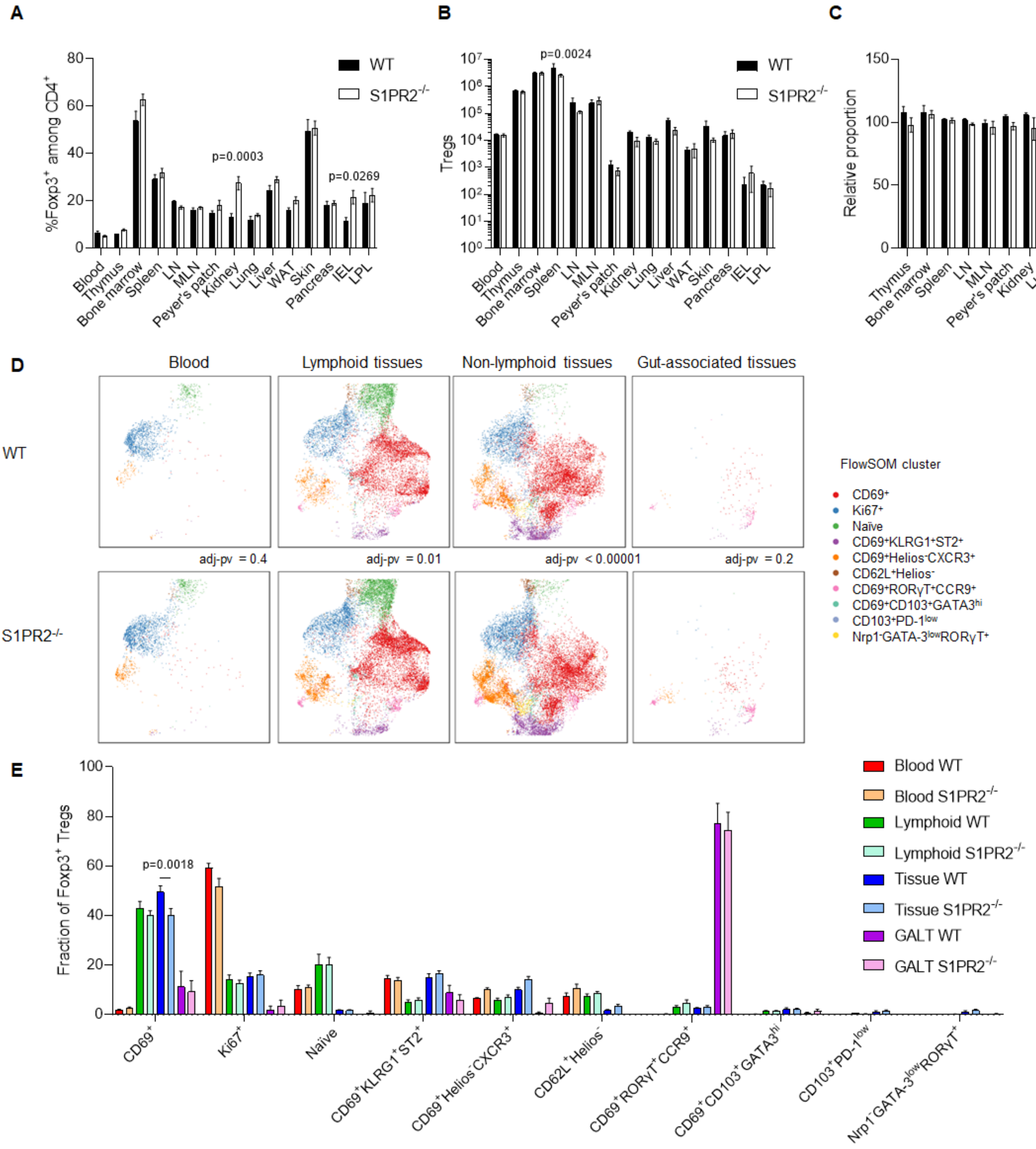

A
%Foxp3+ among CD4+
WT
S1PR2-/-
p=0.0003
p=0.0269
Blood
Thymus
Bone marrow
Spleen
LN
MLN
Peyer's patch
Kidney
Lung
Liver
WAT
Skin
Pancreas
IEL
LPL
B
Tregs
p=0.0024
C
Relative proportion
D
Blood
Lymphoid tissues
Non-lymphoid tissues
Gut-associated tissues
WT
S1PR2-/-
adj-pv = 0.4
adj-pv = 0.01
adj-pv < 0.00001
adj-pv = 0.2
FlowSOM cluster
CD69+
Ki67+
Naïve
CD69+KLRG1+ST2+
CD69+Helios-CXCR3+
CD62L+Helios-
CD69+RORγT+CCR9+
CD69+CD103+GATA3hi
CD103+PD-1low
Nrp1-GATA-3lowRORγT+
E
Fraction of Foxp3+ Tregs
p=0.0018
Blood WT
Blood S1PR2-/-
Lymphoid WT
Lymphoid S1PR2-/-
Tissue WT
Tissue S1PR2-/-
GALT WT
GALT S1PR2-/-

**Supplementary Figure 12**. **BATF deficiency impedes Treg entry into the tissues.** Mixed bone-marrow chimeras were generated with 50% CD45.1 wildtype bone-marrow and 50% CD45.2 *Batf*$^{-/-}$ bone-marrow, transplanted into Rag-deficient recipients. Recipient mice were assessed at 10 weeks post-transplantation by flow cytometry, allowing comparative assessment of the CD45.1 wt and the CD45.2 KO Tregs. **A)** Frequency of Tregs among WT or *Batf*$^{-/-}$ CD4$^{+}$ T cells, across the tissue set assessed. **B)** Absolute numbers of WT and *Batf*$^{-/-}$ Tregs. Statistical analysis by Šídák's multiple comparisons test on 2-way ANOVA. **C)** Frequency of Tregs when normalized to blood (100 being equal). **D)** UMAP plot of flow cytometry phenotype, for WT and *Batf*$^{-/-}$ Tregs, based on the tissue grouping of blood, lymphoid tissues, non-lymphoid tissues and gut-associated tissues, built on the markers CD103, CTLA-4, CD62L, Helios, CCR2, CD44, ICOS, RORγT, PD-1, CXCR3, KLRG1, CCR9, CD95, Neuropilin, Ly-6c, TNFRII, T-bet, GATA-3, CD69, CD25, ST2, IRF4 and Ki67. Statistical analysis by Kolmogorov-Smirnov test (UMAP-diff) comparing WT vs. *Batf*$^{-/-}$ Tregs. **E)** FlowSOM cluster distribution. Statistical analysis by Tukey's multiple comparisons test on 2-way ANOVA.

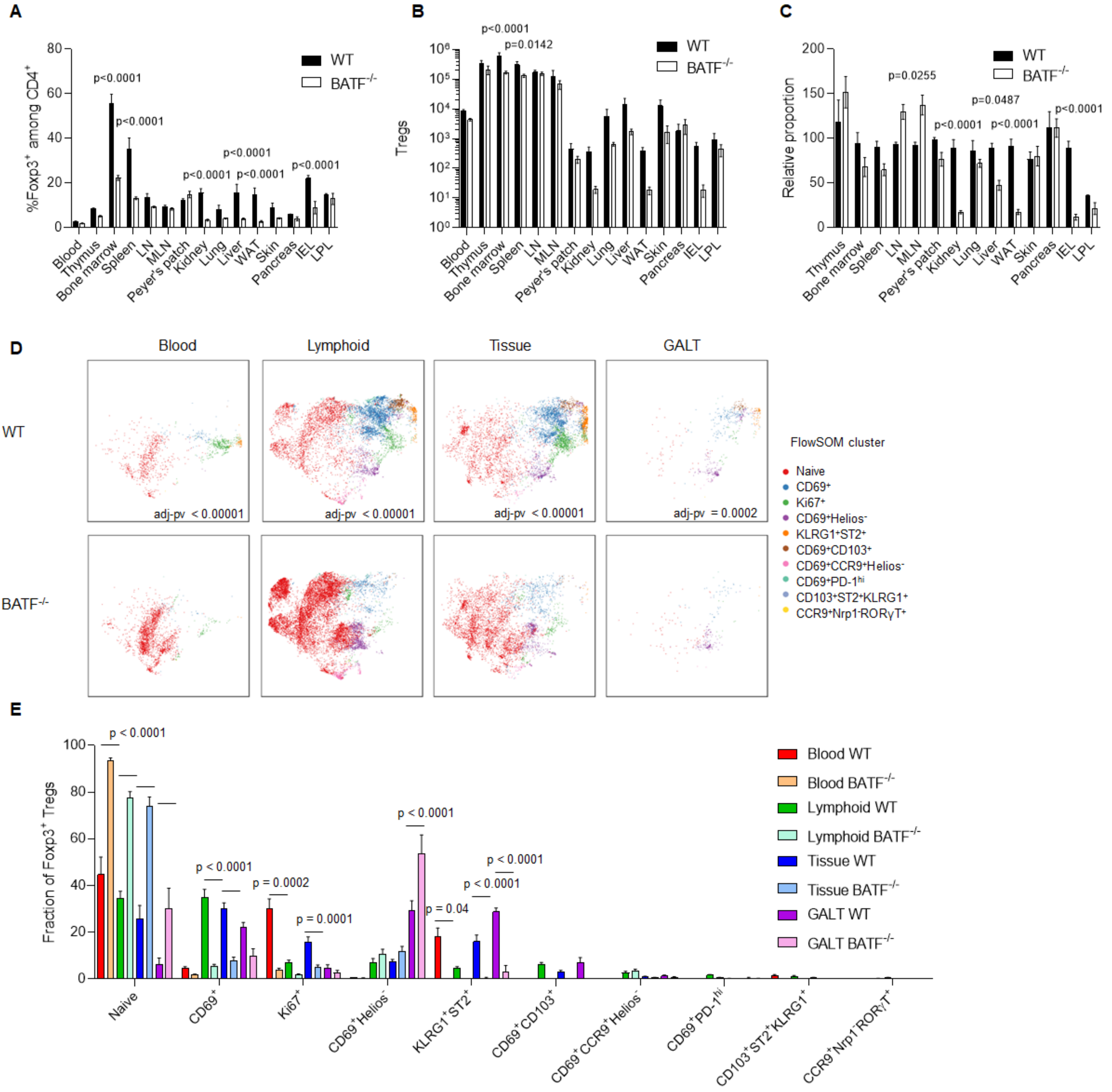
A
%Foxp3+ among CD4+
WT
BATF-/-
p<0.0001
p<0.0001
p<0.0001
p<0.0001
p<0.0001
p<0.0001
Blood
Thymus
Bone marrow
Spleen
LN
MLN
Peyer's patch
Kidney
Lung
Liver
WAT
Skin
Pancreas
IEL
LPL
B
Tregs
p<0.0001
p=0.0142
C
Relative proportion
p=0.0255
p=0.0487
p<0.0001
p<0.0001
p<0.0001
D
Blood
Lymphoid
Tissue
GALT
WT
BATF-/-
adj-pv < 0.00001
adj-pv < 0.00001
adj-pv < 0.00001
adj-pv = 0.0002
FlowSOM cluster
Naive
CD69+
Ki67+
CD69+Helios-
KLRG1+ST2+
CD69+CD103+
CD69+CCR9+Helios-
CD69+PD-1hi
CD103+ST2+KLRG1+
CCR9+Nrp1-RORγT+
E
Fraction of Foxp3+ Tregs
p < 0.0001
p < 0.0001
p = 0.0002
p = 0.0001
p < 0.0001
p = 0.04
p < 0.0001
p < 0.0001
Blood WT
Blood BATF-/-
Lymphoid WT
Lymphoid BATF-/-
Tissue WT
Tissue BATF-/-
GALT WT
GALT BATF-/-
Naive
CD69+
Ki67+
CD69+Helios-
KLRG1+ST2+
CD69+CD103+
CD69+CCR9+Helios-
CD69+PD-1hi
CD103+ST2+KLRG1+
CCR9+Nrp1-RORγT+

**Supplementary Figure 13**. **HIF1α deficiency does not alter Treg fitness or differentiation in tissues.** Mixed bone-marrow chimeras were generated with 50% CD45.1 wildtype bone-marrow and 50% CD45.2 CD4-Cre *Hif1a*$^{fl/fl}$ bone-marrow, transplanted CD45.1/2 recipients. Recipient mice were assessed at 8 weeks post-transplantation by flow cytometry with intravenous CD45 labeling, allowing comparative assessment of the CD45.1 wt and the CD45.2 KO Tregs. **A)** Frequency of Tregs among WT or CD4-Cre *Hif1a*$^{fl/fl}$ CD4$^{+}$ T cells, across the tissue set assessed. **B)** Absolute numbers of WT and CD4-Cre *Hif1a*$^{fl/fl}$ Tregs. Statistical analysis by Šídák's multiple comparisons test on 2-way ANOVA. **C)** Frequency of Tregs when normalized to blood (100 being equal). **D)** tSNE plot of flow cytometry phenotype, for WT and CD4-Cre *Hif1a*$^{fl/fl}$ Tregs, based on the tissue grouping of blood, lymphoid tissues, non-lymphoid tissues and gut-associated tissues, built on the markers CD103, CTLA-4, CD62L, GITR, CXCR6, Helios, CCR2, CD44, ICOS, RORγT, PD-1, CD127, LAG3, CD5, Tim-3, CXCR3, KLRG1, CCR9, CD95, Neuropilin, Ly-6c, Blimp1, T-bet, CD69, CD25, CD137, CD38, ST2, CCR8, CCR7 and Ki67. Statistical analysis by Kolmogorov-Smirnov test with Holm correction (tSNE-diff) comparing WT vs. CD4-Cre *Hif1a*$^{fl/fl}$ Tregs. **E)** FlowSOM cluster distribution. Statistical analysis by Tukey's multiple comparisons test on 2-way ANOVA.

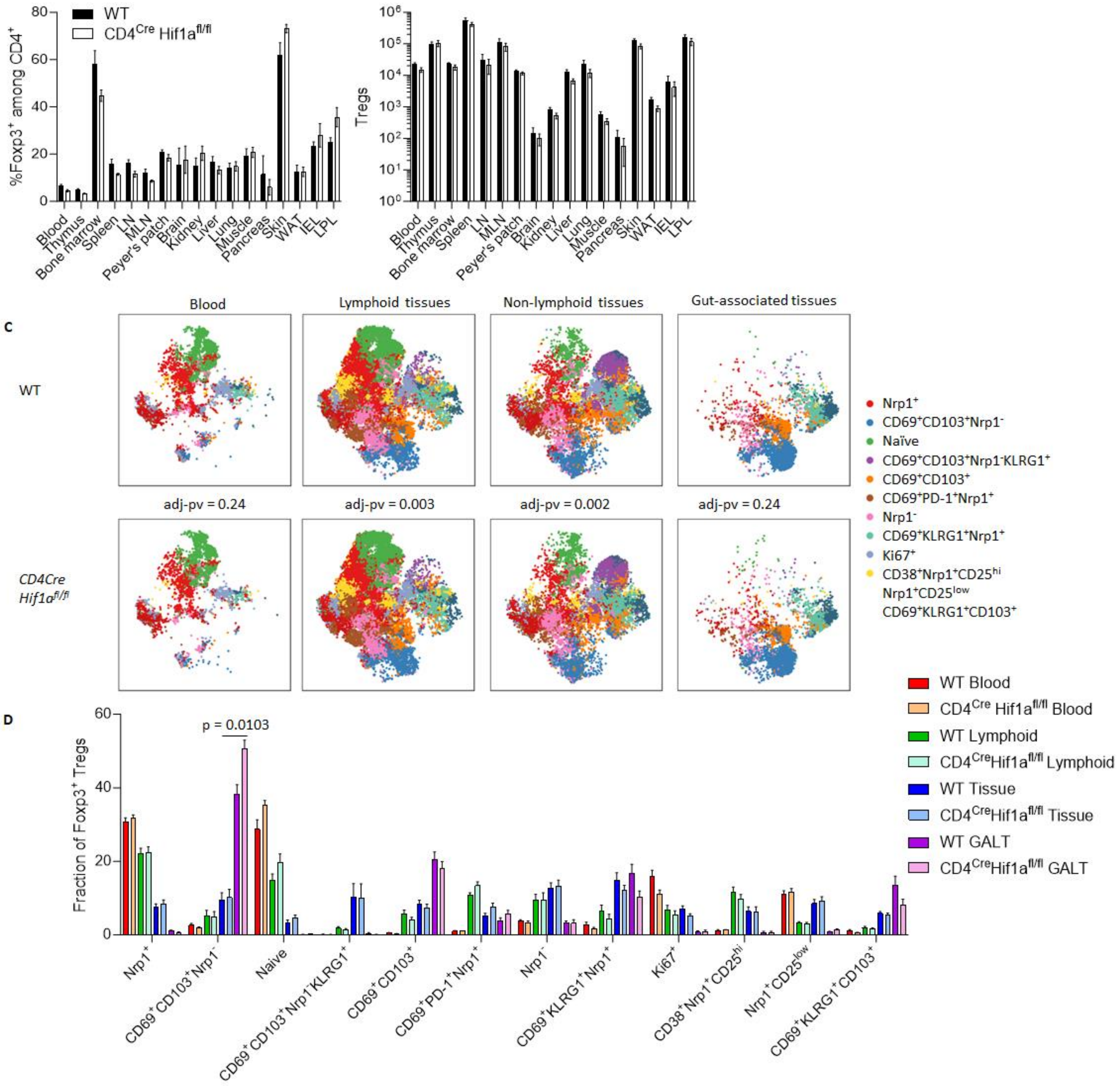
WT
CD4Cre Hif1afl/fl
%Foxp3+ among CD4+
Tregs
Blood
Thymus
Bone marrow
Spleen
LN
MLN
Peyer's patch
Brain
Kidney
Liver
Lung
Muscle
Pancreas
Skin
WAT
IEL
LPL
C
Blood
Lymphoid tissues
Non-lymphoid tissues
Gut-associated tissues
WT
CD4Cre Hif1afl/fl
adj-pv = 0.24
adj-pv = 0.003
adj-pv = 0.002
adj-pv = 0.24
Nrp1+
CD69+CD103+Nrp1-
Naïve
CD69+CD103+Nrp1-KLRG1+
CD69+CD103+
CD69+PD-1+Nrp1+
Nrp1-
CD69+KLRG1+Nrp1+
Ki67+
CD38+Nrp1+CD25hi
Nrp1+CD25low
CD69+KLRG1+CD103+
D
Fraction of Foxp3+ Tregs
p = 0.0103
WT Blood
CD4Cre Hif1afl/fl Blood
WT Lymphoid
CD4Cre Hif1afl/fl Lymphoid
WT Tissue
CD4Cre Hif1afl/fl Tissue
WT GALT
CD4Cre Hif1afl/fl GALT

**Supplementary Figure 14**. **Areg deficiency alters skin and gut tissue Tregs, with minor impact on other tissue Treg populations.** Areg KO and wt mice were injected with intravenous anti-CD45 antibody label and assessed for flow cytometry at 11 weeks of age. **A)** Frequency of Tregs among WT or *Areg*$^{-/-}$ CD4$^{+}$ T cells, across the tissue set assessed. **B)** Absolute numbers of WT and *Areg*$^{-/-}$ Tregs. Statistical analysis by Šídák's multiple comparisons test on 2-way ANOVA. **C)** UMAP plot of flow cytometry phenotype, for WT and *Areg*$^{-/-}$ Tregs, based on the tissue grouping of blood, lymphoid tissues, non-lymphoid tissues and gut-associated tissues, built on the markers CD103, CTLA-4, CD62L, CXCR6, Helios, CCR2, CD44, ICOS, RORγT, PD-1, CXCR3, KLRG1, CCR9, CD95, Neuropilin, Ly-6C, TNFRII, T-bet, GATA-3, CD69, CD25, ST2, IRF4 and Ki67. No statistical difference by Kolmogorov-Smirnov test with Holm correction (UMAP-diff) comparing WT vs. *Areg*$^{-/-}$ Tregs. **D)** FlowSOM cluster distribution. No statistical difference by Tukey's multiple comparisons test on 2-way ANOVA. **E)** UMAP plot of flow cytometry phenotype, for WT and *Areg*$^{-/-}$ Tregs in the skin, with **F)** FlowSOM cluster distribution. **G)** UMAP plot of flow cytometry phenotype, for WT and *Areg*$^{-/-}$ Tregs for IEL and LPL in the small intestine, with **H)** FlowSOM cluster distribution. Statistical analysis by Tukey's multiple comparison test on 2-way ANOVA.

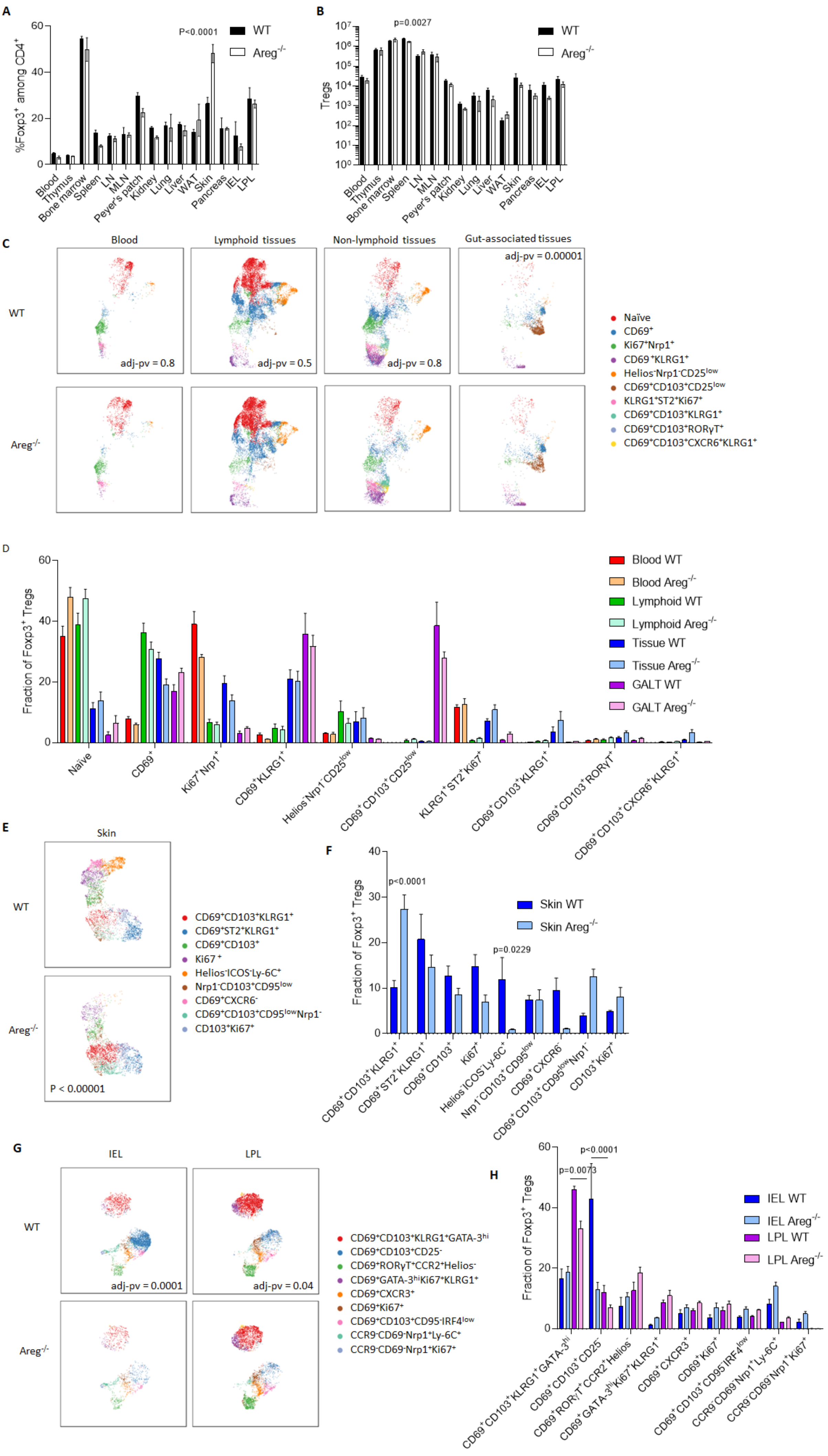
A
%Foxp3+ among CD4+
P<0.0001
WT
Areg-/-
Blood
Thymus
Bone marrow
Spleen
LN
MLN
Peyer's patch
Kidney
Lung
Liver
WAT
Skin
Pancreas
IEL
LPL
B
Tregs
p=0.0027
C
Blood
Lymphoid tissues
Non-lymphoid tissues
Gut-associated tissues
adj-pv = 0.8
adj-pv = 0.5
adj-pv = 0.8
adj-pv = 0.00001
Naïve
CD69+
Ki67+Nrp1+
CD69+KLRG1+
Helios-Nrp1-CD25low
CD69+CD103+CD25low
KLRG1+ST2+Ki67+
CD69+CD103+KLRG1+
CD69+CD103+RORγT+
CD69+CD103+CXCR6+KLRG1+
D
Fraction of Foxp3+ Tregs
Blood WT
Blood Areg-/-
Lymphoid WT
Lymphoid Areg-/-
Tissue WT
Tissue Areg-/-
GALT WT
GALT Areg-/-
E
Skin
P < 0.00001
CD69+CD103+KLRG1+
CD69+ST2+KLRG1+
CD69+CD103+
Ki67+
Helios-ICOS-Ly-6C+
Nrp1-CD103+CD95low
CD69+CXCR6-
CD69+CD103+CD95lowNrp1-
CD103+Ki67+
F
p<0.0001
p=0.0229
Skin WT
Skin Areg-/-
G
IEL
LPL
adj-pv = 0.0001
adj-pv = 0.04
CD69+CD103+KLRG1+GATA-3hi
CD69+CD103+CD25-
CD69+RORγT+CCR2+Helios-
CD69+GATA-3hiKi67+KLRG1+
CD69+CXCR3+
CD69+Ki67+
CD69+CD103+CD95-IRF4low
CCR9-CD69-Nrp1+Ly-6C+
CCR9-CD69-Nrp1+Ki67+
H
p=0.0073
p<0.0001
IEL WT
IEL Areg-/-
LPL WT
LPL Areg-/-

**Supplementary Figure 15. CD103 is not required for Treg tissue residency. A)** Sanger sequence traces showing the deletion of 10bp of *CD103* locus in exon 5. **B)** Histogram showing CD103 expression levels in WT and $CD103^{KO}$ splenocytes (n=4-6). **C)** Percentage of $CD45^{+}$ $CD4^{+}$ cells in different tissues from WT and $CD103^{KO}$ mice aged 13 weeks (n=4-6). **D)** Percentage of $CD45^{+}$ $CD8^{+}$ cells in different tissues from WT and $CD103^{KO}$ mice (n=4-6). **E)** Percentage of $CD4^{+}$ $Foxp3^{+}$ Treg cells in different tissues from WT and $CD103^{KO}$ mice (n=4-6). **F)** Percentage of $CD4^{+}$ $Foxp3^{+}$ $CD62L^{+}$ $CD44^{low}$ Treg cells in different tissues from WT and $CD103^{KO}$ mice (n=4-6). **G)** Percentage of $CD4^{+}$ $Foxp3^{+}$ $CD44^{high}$ $CD62L^{-}$ Treg cells in different tissues from WT and $CD103^{KO}$ mice (n=4-6). **H)** Percentage of $CD4^{+}$ $Foxp3^{+}$ $CD69^{+}$ Treg cells in different tissues from WT and $CD103^{KO}$ mice (n=4-6). **I)** Percentage of $CD4^{+}$ $Foxp3^{+}$ $KLRG1^{+}$ Treg cells in different tissues from WT and $CD103^{KO}$ mice (n=4-6). **J**) tSNE plots of flow cytometry phenotype, for WT and CD103 KO Tregs from perfused mice, based on the tissue grouping of blood, lymphoid tissues, non-lymphoid tissues and gut-associated tissues, built on the markers CD103, CTLA-4, CD62L, Helios, CD44, ICOS, PD-1, KLRG1, Neuropilin, T-bet, CD69, CD25, ST2 and Ki67. Statistical analysis by Kolmogorov-Smirnov test (tSNE-diff) comparing WT vs. *Cd103*$^{-/-}$ Tregs. K) Dendrogram built on tSNE crossentropy of flow cytometry phenotype. Abbreviations: BM = Bone marrow; TH = Thymus; LN = Lymph nodes; SP = Spleen; MLN = Mesenteric lymph nodes; PP = Payer’s patches; BL = Blood; ADR = Adrenals; IEL = Intraepithelial lymphocytes; LPL = lamina propria T lymphocytes; KID = Kidney; LIV = Liver; PAN = Pancreas; WAT = White adipose tissue. All data are means ± SEM. *p < 0.05.

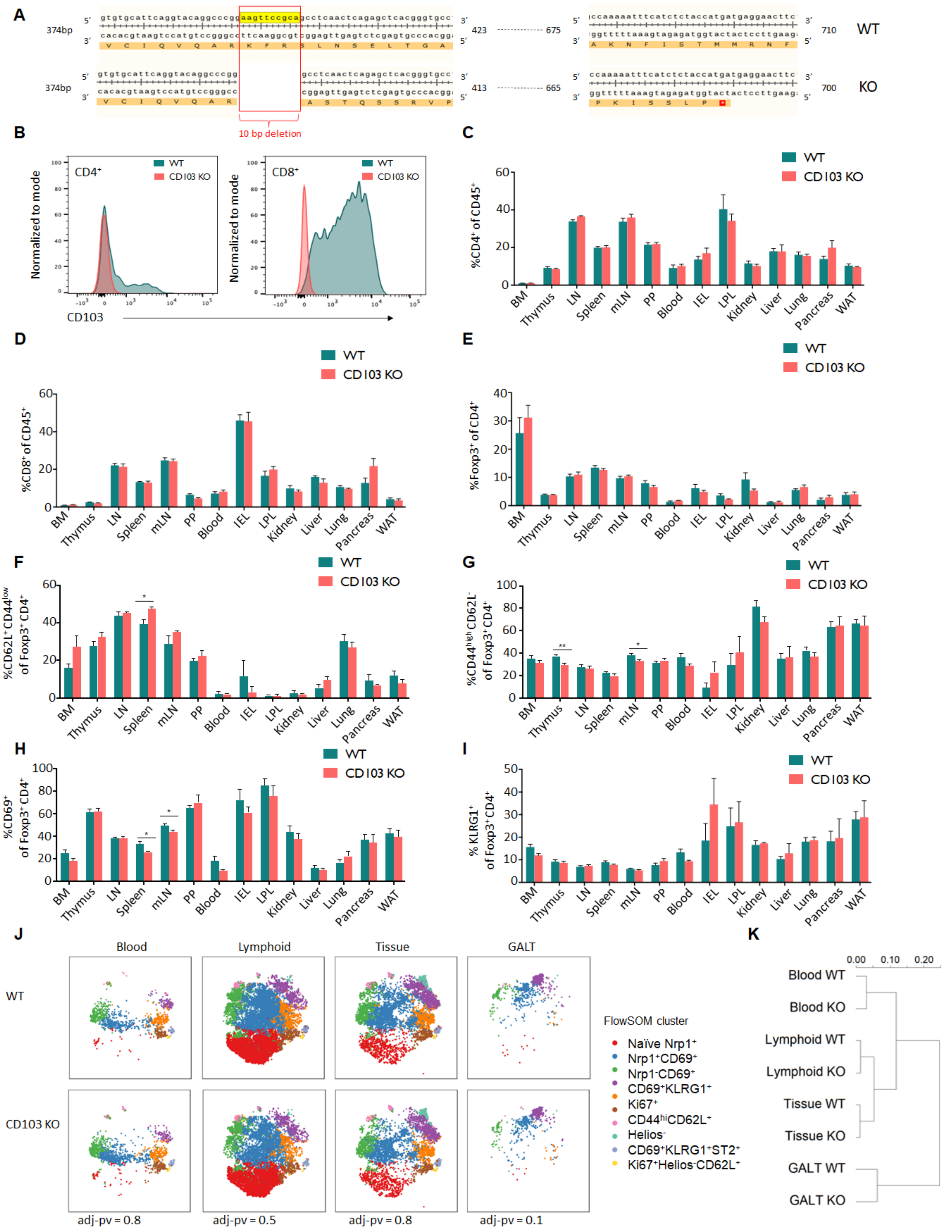

A
B
C
D
E
F
G
H
I
J
K
WT
KO
374bp
10 bp deletion
CD4+
CD8+
WT
CD103 KO
Normalized to mode
CD103
%CD4+ of CD45+
%CD8+ of CD45+
%Foxp3+ of CD4+
%CD62L+CD44low of Foxp3+CD4+
%CD44highCD62L- of Foxp3+CD4+
%CD69+ of Foxp3+CD4+
% KLRG1+ of Foxp3+CD4+
BM
Thymus
LN
Spleen
mLN
PP
Blood
IEL
LPL
Kidney
Liver
Lung
Pancreas
WAT
Blood
Lymphoid
Tissue
GALT
CD103 KO
FlowSOM cluster
Naïve Nrp1+
Nrp1+CD69+
Nrp1-CD69+
CD69+KLRG1+
Ki67+
CD44hiCD62L+
Helios-
CD69+KLRG1+ST2+
Ki67+Helios-CD62L+
adj-pv = 0.8
adj-pv = 0.5
adj-pv = 0.8
adj-pv = 0.1
0.00
0.10
0.20
Blood WT
Blood KO
Lymphoid WT
Lymphoid KO
Tissue WT
Tissue KO
GALT WT
GALT KO

**Supplementary Figure 16. Induced deficiency in CD69 does not substantially alter tissue Treg number or phenotype.** *Foxp3*$^{ERT2Cre}$ *CD69*$^{fl/fl}$ mice and *Foxp3*$^{ERT2Cre}$ *CD69*$^{wt/wt}$ mice were treated with tamoxifen (0.2mg/g) by oral gavage weekly for four weeks. Mice were assessed by flow cytometry with intravenous labeling of CD45. **A)** Frequency of Tregs among *Foxp3*$^{ERT2Cre}$ *CD69*$^{+/+}$ or *CD69*$^{fl/fl}$ CD4$^{+}$ T cells, across the tissue set assessed. **B)** Absolute numbers of *Foxp3*$^{ERT2Cre}$ *CD69*$^{+/+}$ or *CD69*$^{fl/fl}$ Tregs. Statistical analysis by Šídák's multiple comparisons test on 2-way ANOVA. **C)** Frequency of Tregs when normalized to blood (100 being equal). **D)** tSNE plot of flow cytometry phenotype, for *Foxp3*$^{ERT2Cre}$ *CD69*$^{wt/wt}$ or *CD69*$^{fl/fl}$ Tregs, based on the tissue grouping of blood, lymphoid tissues, non-lymphoid tissues and gut-associated tissues, built on the markers CD103, CTLA-4, CD62L, Helios, CCR2, CD44, ICOS, RORγT, PD-1, LAG3, GITR, CXCR3, KLRG1, CCR9, CD95, Neuropilin, Ly-6c, Nur77 , T-bet, CD69, CD25, GATA-3, , ST2, CCR7 and Ki67. Statistical analysis by Kolmogorov-Smirnov test with Holm correction (tSNE-diff) comparing *Foxp3*$^{ERT2Cre}$ *CD69*$^{+/+}$ vs *CD69*$^{fl/fl}$ Tregs. **E)** FlowSOM cluster distribution. Statistical analysis by Tukey's multiple comparisons test on 2-way ANOVA.

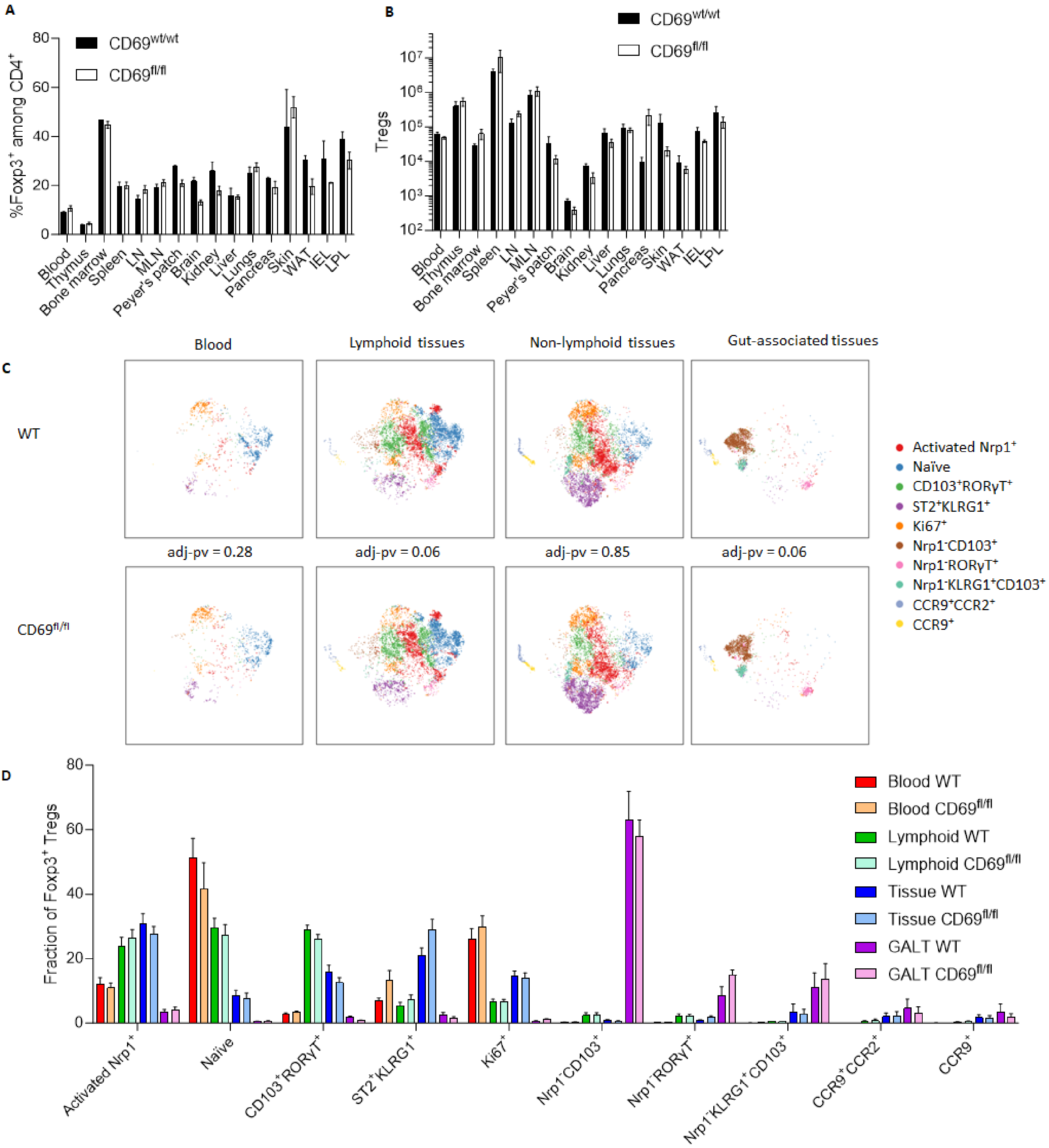
A
%Foxp3+ among CD4+
CD69wt/wt
CD69fl/fl
80
60
40
20
0
Blood
Thymus
Bone marrow
Spleen
LN
MLN
Peyer's patch
Brain
Kidney
Liver
Lungs
Pancreas
Skin
WAT
IEL
LPL
B
Tregs
CD69wt/wt
CD69fl/fl
10^7
10^6
10^5
10^4
10^3
10^2
Blood
Thymus
Bone marrow
Spleen
LN
MLN
Peyer's patch
Brain
Kidney
Liver
Lungs
Pancreas
Skin
WAT
IEL
LPL
C
Blood
Lymphoid tissues
Non-lymphoid tissues
Gut-associated tissues
WT
CD69fl/fl
adj-pv = 0.28
adj-pv = 0.06
adj-pv = 0.85
adj-pv = 0.06
Activated Nrp1+
Naïve
CD103+RORγT+
ST2+KLRG1+
Ki67+
Nrp1-CD103+
Nrp1-RORγT+
Nrp1-KLRG1+CD103+
CCR9+CCR2+
CCR9+
D
Fraction of Foxp3+ Tregs
80
60
40
20
0
Blood WT
Blood CD69fl/fl
Lymphoid WT
Lymphoid CD69fl/fl
Tissue WT
Tissue CD69fl/fl
GALT WT
GALT CD69fl/fl
Activated Nrp1+
Naïve
CD103+RORγT+
ST2+KLRG1+
Ki67+
Nrp1-CD103+
Nrp1-RORγT+
Nrp1-KLRG1+CD103+
CCR9+CCR2+
CCR9+

**Supplementary Figure 17. Markov chain models for tissue Treg cellular kinetics.** CD45.1 mice were parabiosed to CD45.2 mice. Pairs of parabiotic animals were sacrificed at weeks 1, 2, 4, 8, and 12 for tissue analysis by flow cytometry (n=11,12,18,16,14). Markov chains were built to model the changes in cell state and tissue exchange, with each tissue built using a model containing the tissue, blood, and combined other tissues. Displayed are the original data points superimposed on the model predictions for resting Treg (left), activated Treg (middle) and $CD69^+$ Treg (right) from **A.** bone-marrow, **B.** LN, **C.** mLN, **D.** PP, **E.** spleen, **F.** adrenals, **G.** brain, **H.** kidney, **I.** liver, **J.** lung, **K.** muscle, **L.** pancreas, **M.** skin, **N.** WAT, **O.** IEL, **P.** LPL, **Q.** and blood. The blood model results displayed is from the spleen, blood and other tissue model, while all other tissues are displayed from their own model.

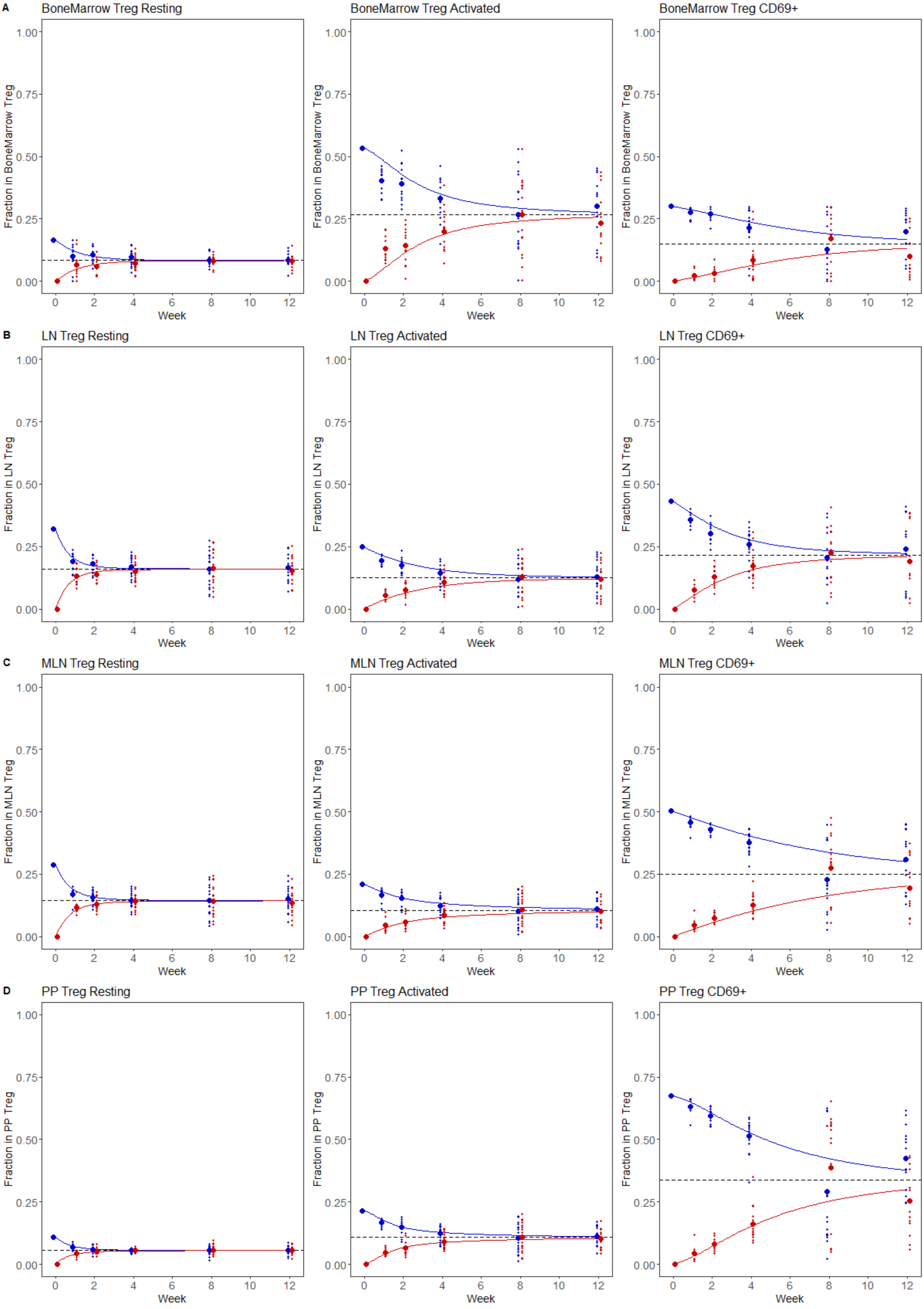
A
BoneMarrow Treg Resting
BoneMarrow Treg Activated
BoneMarrow Treg CD69+
Fraction in BoneMarrow Treg
B
LN Treg Resting
LN Treg Activated
LN Treg CD69+
Fraction in LN Treg
C
MLN Treg Resting
MLN Treg Activated
MLN Treg CD69+
Fraction in MLN Treg
D
PP Treg Resting
PP Treg Activated
PP Treg CD69+
Fraction in PP Treg
Week
0.00
0.25
0.50
0.75
1.00
0
2
4
6
8
10
12

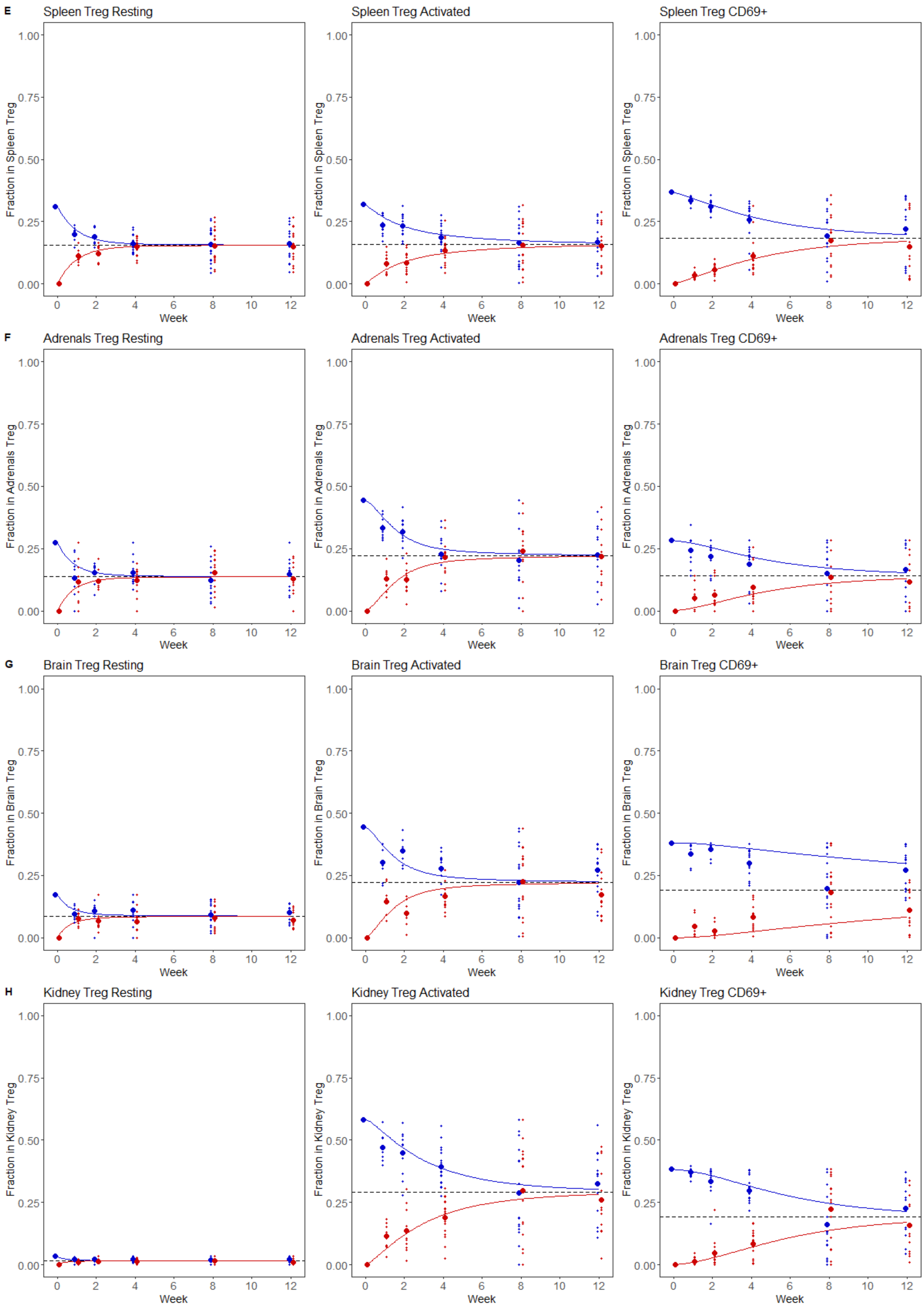
E
Spleen Treg Resting
Spleen Treg Activated
Spleen Treg CD69+
Fraction in Spleen Treg
F
Adrenals Treg Resting
Adrenals Treg Activated
Adrenals Treg CD69+
Fraction in Adrenals Treg
G
Brain Treg Resting
Brain Treg Activated
Brain Treg CD69+
Fraction in Brain Treg
H
Kidney Treg Resting
Kidney Treg Activated
Kidney Treg CD69+
Fraction in Kidney Treg
0.00
0.25
0.50
0.75
1.00
0
2
4
6
8
10
12
Week

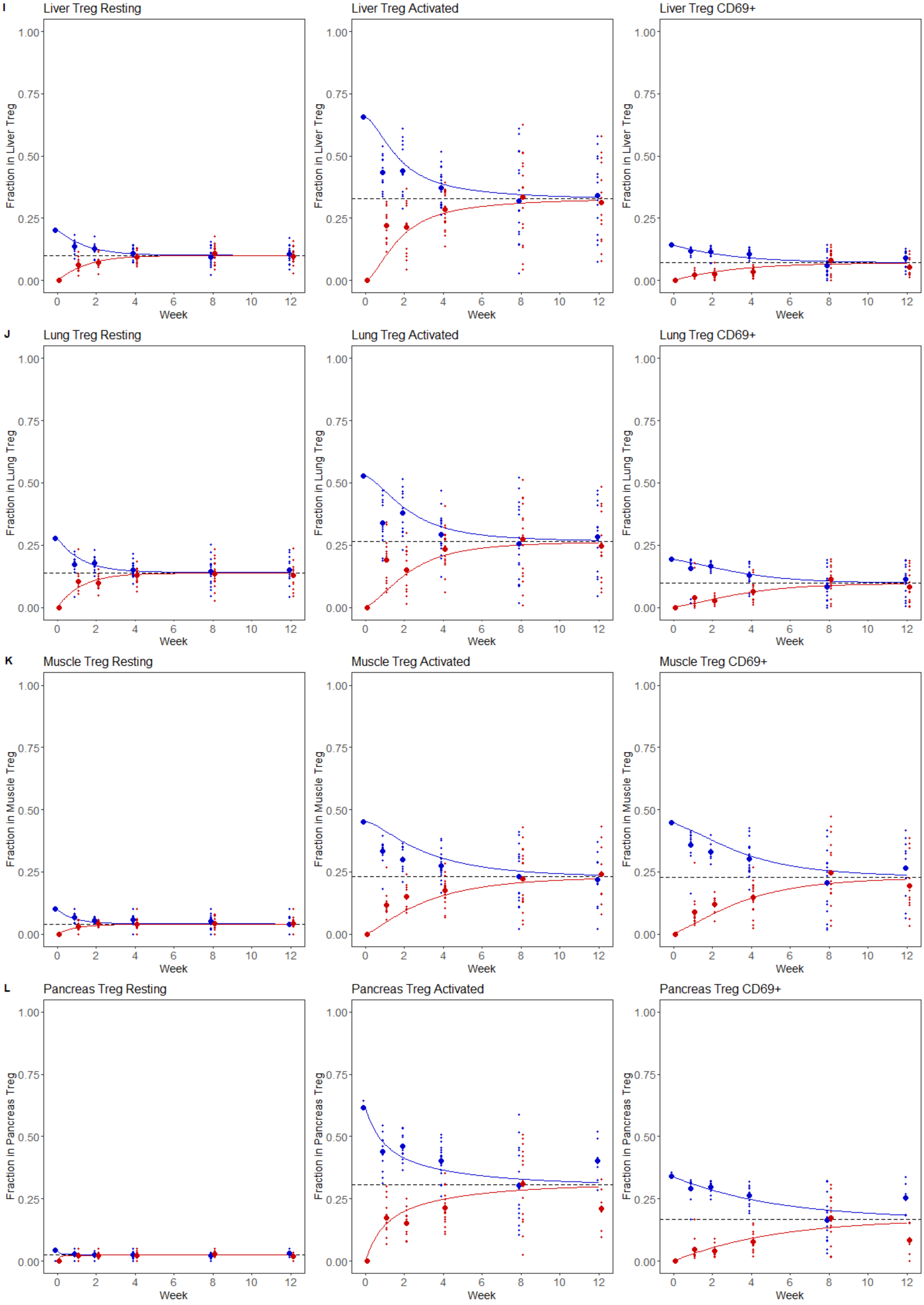

I
Liver Treg Resting
Liver Treg Activated
Liver Treg CD69+
Fraction in Liver Treg
J
Lung Treg Resting
Lung Treg Activated
Lung Treg CD69+
Fraction in Lung Treg
K
Muscle Treg Resting
Muscle Treg Activated
Muscle Treg CD69+
Fraction in Muscle Treg
L
Pancreas Treg Resting
Pancreas Treg Activated
Pancreas Treg CD69+
Fraction in Pancreas Treg
0.00
0.25
0.50
0.75
1.00
0
2
4
6
8
10
12
Week

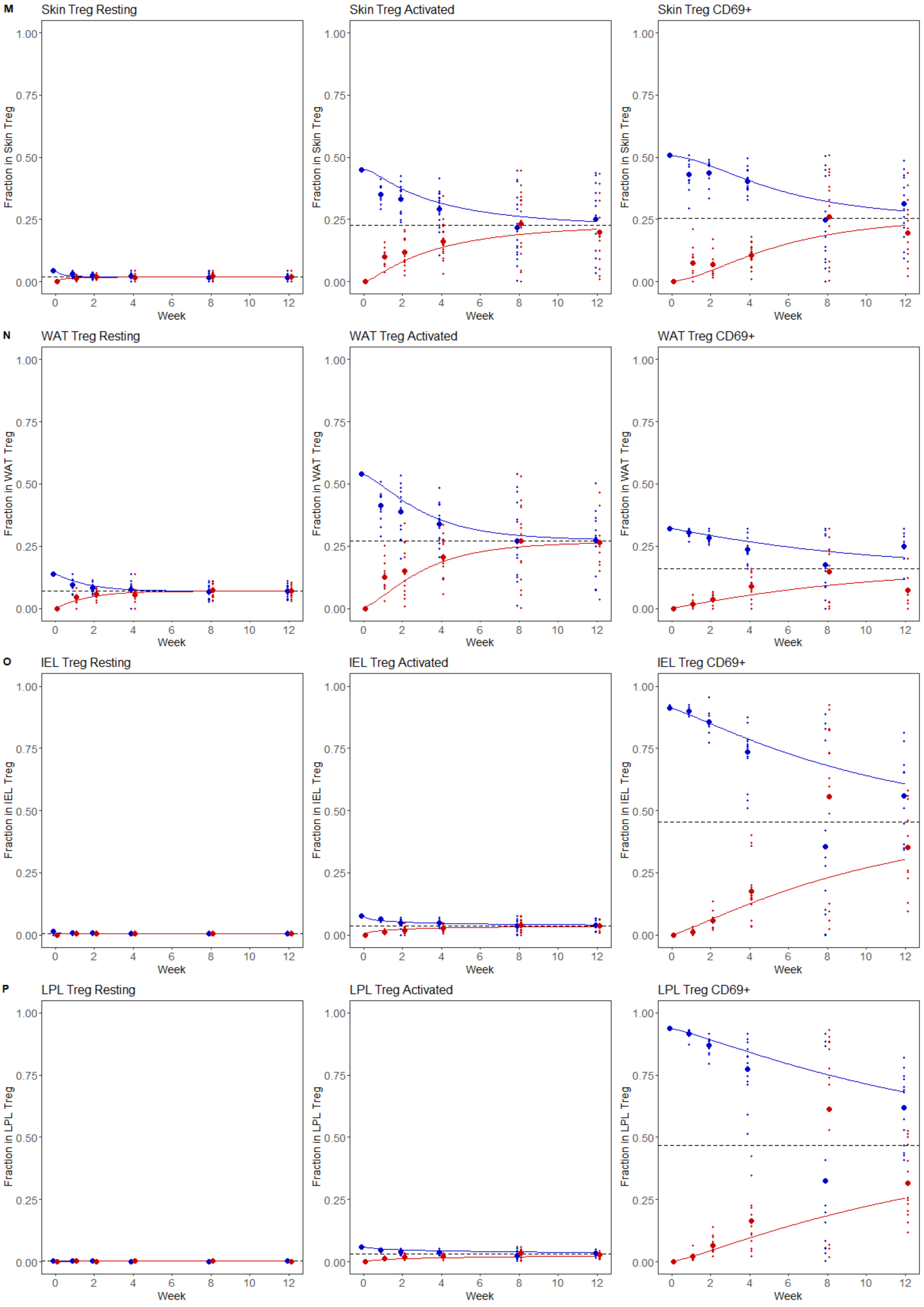
M
Skin Treg Resting
Skin Treg Activated
Skin Treg CD69+
Fraction in Skin Treg
N
WAT Treg Resting
WAT Treg Activated
WAT Treg CD69+
Fraction in WAT Treg
O
IEL Treg Resting
IEL Treg Activated
IEL Treg CD69+
Fraction in IEL Treg
P
LPL Treg Resting
LPL Treg Activated
LPL Treg CD69+
Fraction in LPL Treg
0.00
0.25
0.50
0.75
1.00
0
2
4
6
8
10
12
Week

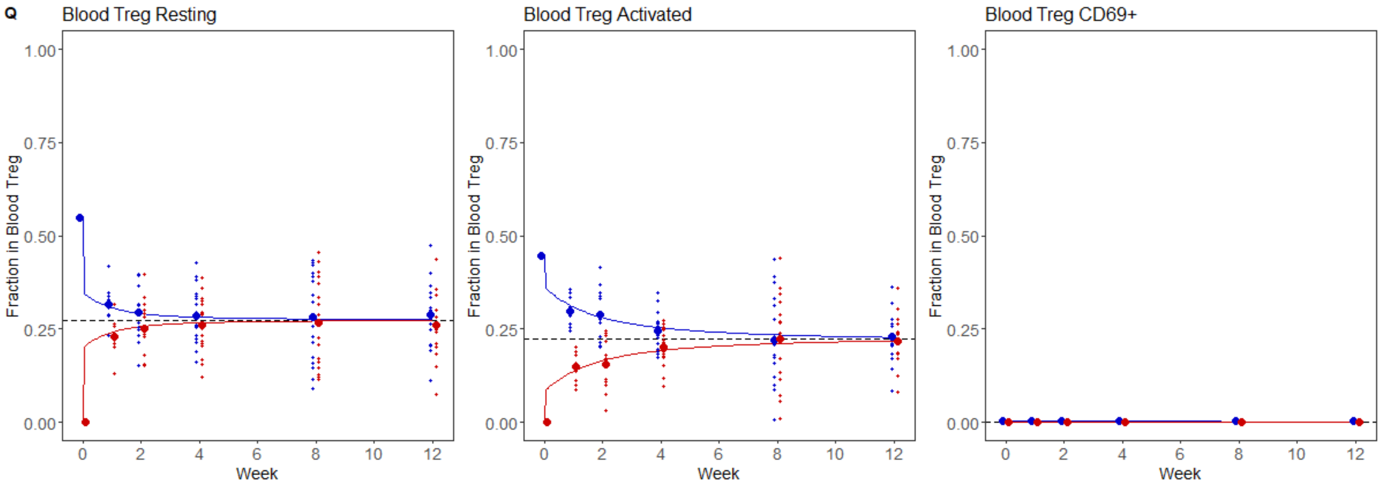

Q
Blood Treg Resting
Blood Treg Activated
Blood Treg CD69+
Fraction in Blood Treg
1.00
0.75
0.50
0.25
0.00
0
2
4
6
8
10
12
Week

**Supplementary Figure 18. Distribution of dwell times within CD69+ tissue Tregs based on Markov chain models.** CD45.1 mice were parabiosed to CD45.2 mice. Pairs of parabiotic animals were sacrificed at weeks 1, 2, 4, 8, and 12 for tissue analysis by flow cytometry (n =11,12,18,16,14}). Markov chains were built to model the changes in cell state and tissue exchange, with each tissue built using a model containing the tissue, blood, and combined other tissues. Displayed are the probabilistic distributions of dwell times for CD69$^{+}$ Tregs, including mean and percentiles (1%, 25%, 50%, 75%, 90%) for the population. Calculates for **A.** bone-marrow, **B.** lymph nodes, **C.** mLN, **D.** Peyer's Patches, **E.** spleen, **F.** adrenals, **G.** brain, **H.** kidney, **I.** liver, **J.** lung, **K.** muscle, **L.** pancreas, **M.** skin, **N.** white adipose tissue, **O.** IEL, **P.** and LPL. The blood model results displayed is from the spleen, blood and other tissue model, while all other tissues are displayed from their own model.

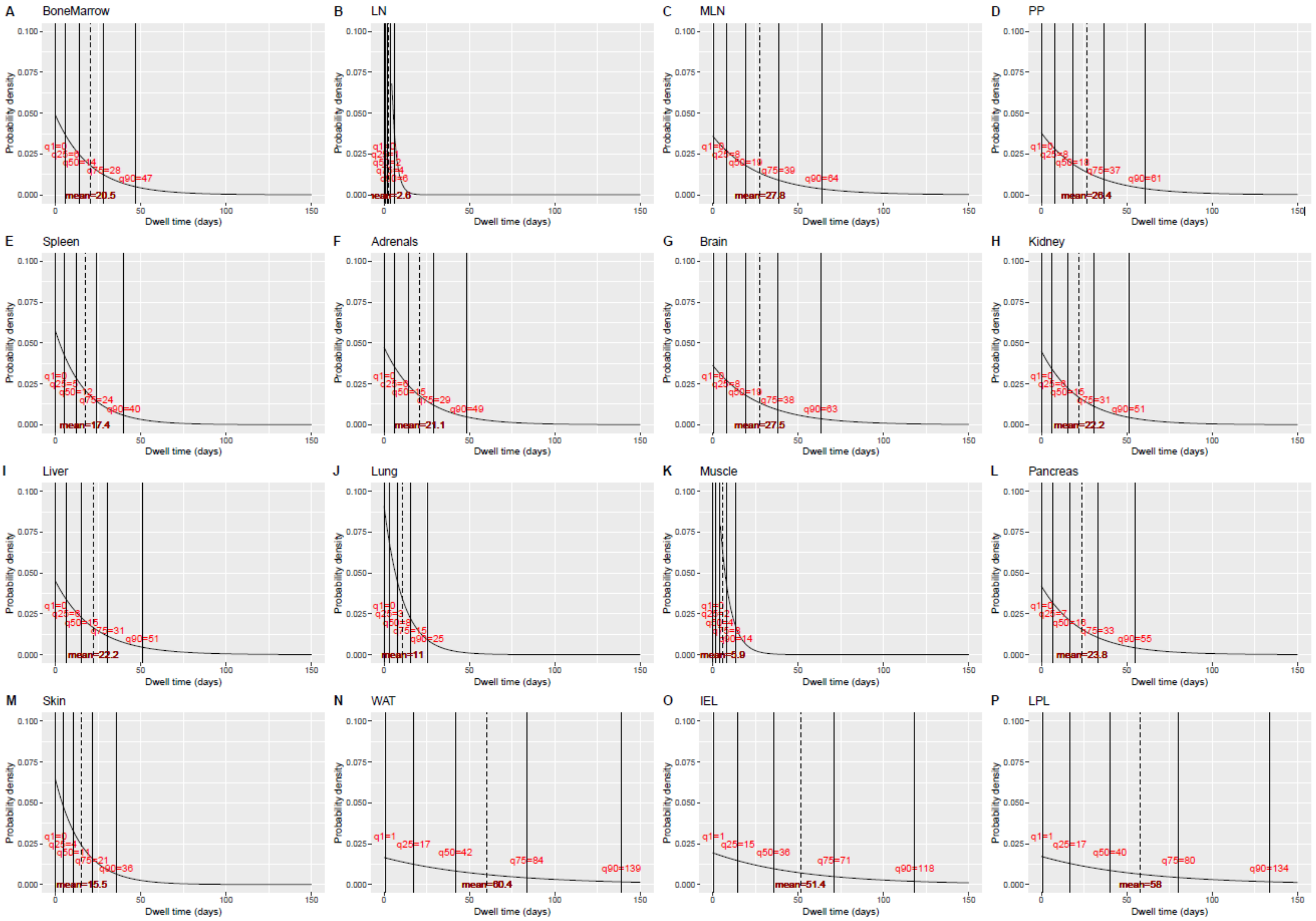

**Supplementary Figure 19. KLRG1 expression does not represent a terminal tissue-resident fate in Tregs.** *Klrg1^Cre^ RosaAi14* mice aged 15 weeks were assessed by high dimensional flow cytometry with intravenous CD45 labeling across lymphoid tissues, non-lymphoid tissues and gut-associated tissues (n=5). A) Fraction of Tregs expressing KLRG1 or ex-KLRG1-expressing ($KLRG1^-$ $tdTomato^+$), across the tested tissues. **B)** UMAP of neverKLRG1 ($KLRG1^-$ $tdTomato^-$), $KLRG1^+$ and exKLRG1 ($KLRG1^-$ $tdTomato^+$) Tregs, amalgamated from all tissues, showing that KLRG1 and exKLRG1 cells are enriched for activated, tissue-resident Treg phenotypes. UMAP built on the markers CTLA-4, Helios, CCR2, CD44, ICOS, RORγT, PD-1, CXCR3, CD95, Neuropilin, Ly-6c, CD69, CD25, ST2, CD103, CD127, CD62L, GITR, IRF4 and Ki67. **C)** Frequency of flowSOM clusters in neverKLRG1, $KLRG1^+$ and exKLRG1 Treg subsets.

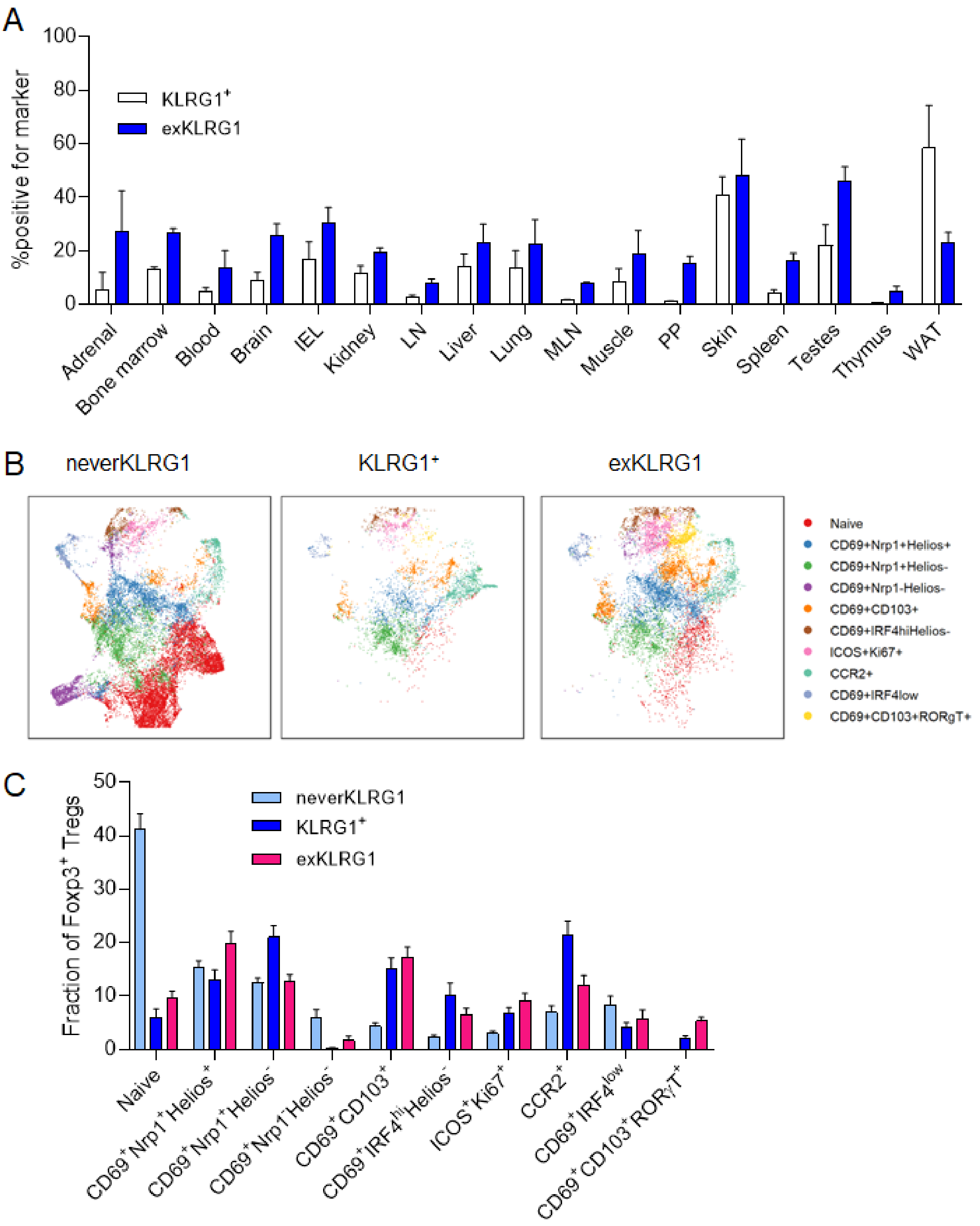

**Supplementary Figure 20. Immunopeptidome overlap between tissues.** Reanalysis of a published H2D$^{d}$ immunopeptidome of 19 normal tissues from C57BL/6 mice mass spectrometry dataset [35]. **A)** The fraction of spectral count that correspond to peptides detected only in a single tissue (Unique), in 2-9 tissues (Some), in 10-18 tissue (Most) or in all tissues (All). **B)** Distribution of the spectral count frequency of each peptide depending on their sharing group.

A

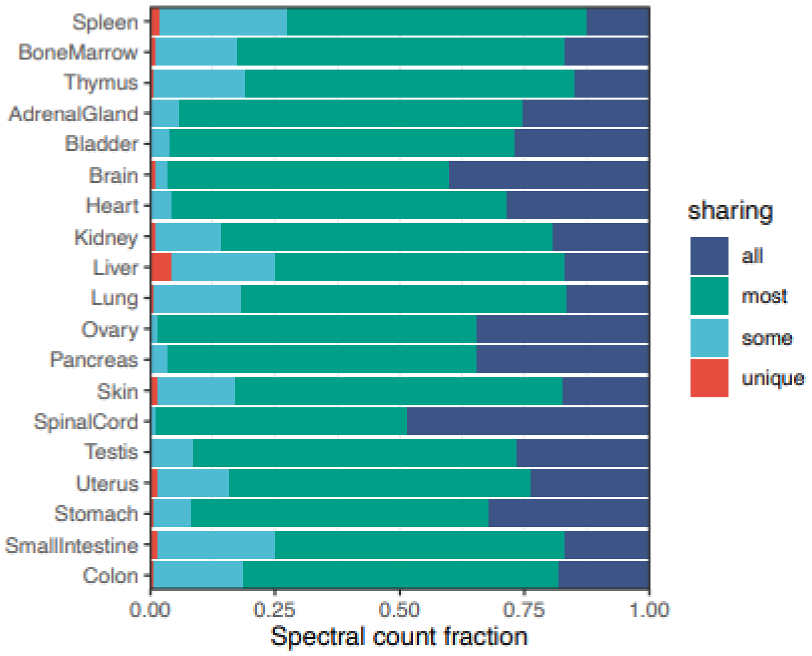


B

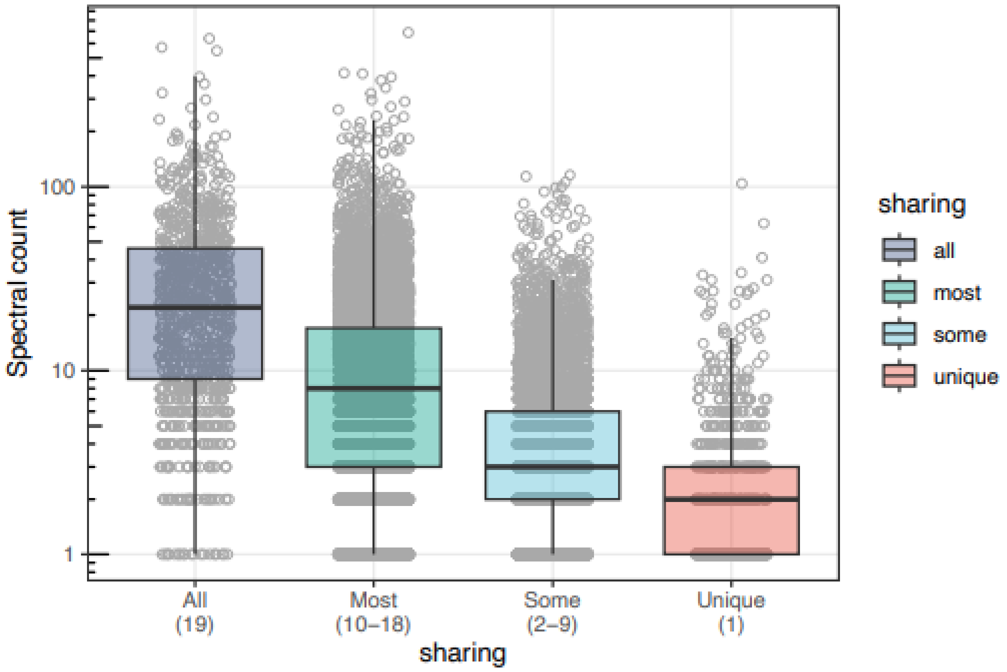

**Supplementary Figure 21. Selection of multiplexed tissue Treg TCR retrogenics to reflect the common properties of tissue Treg TCRs.** *Foxp3*$^{Thy1.1}$ male mice at 16 weeks of age were injected with intravenous anti-CD45 antibody label prior to FACS sorting of Tregs from blood, kidney, liver, pancreas and LPL (n=4) for analysis by scTCRseq. **A)** Top 20 TCRs from the scSeq database, ranked based on the total number of samples the clonotype was detected in. Heatmaps represent the frequency of detection of each clone, across the 5 tested tissues and 4 replicate mice. TCR clones 01-10, selected for retrogenic analysis, are marked. **B)** Top 10 TCRs from the scSeq database, ranked based on the frequency with which the clonotypes were detected in mouse 1 kidney, liver, pancreas and LPL samples. Heatmaps represent the frequency of detection of each clone, across the 5 tested tissues and 4 replicate mice. TCR clones 11-21, constituting the top three clones from each tissue, were selected for retrogenic analysis. **C)** Plot showing tissue Treg TCRs, based on the total number of cells in which the clone was detected and the number of samples it was detected in (including both replicate mice and different tissue samples, not including the blood). Selected clones are indicated and named. **D)** Bar graph based on public tissue Treg clones (identified in at least two mice, with at least one count per mouse), indicating the fraction that were observed in only a single tissue across multiple mice ("tissue-restricted"), a single tissue plus blood ("tissue/blood-restricted") or across multiple tissues ("multi-tissue). Parallel analysis made on the 20 selected TCRs. **E)** Rag-deficient bone-marrow stem cells were individually transduced with the 20 retroviruses and pooled for reconstitution of irradiated mice. After 10 weeks, tissue samples were prepared for analysis by flow cytometry. FlowCodeDecoder was used to assign each T cell to the appropriate TCR clone, based on expression of the 7 epitopes. Individual results from T cells derived from each of the FlowCode retrogenic TCRs, across the assessed tissues of spleen, LN, kidney, lung, liver, pancreas and LPLs. For each clone and tissue is shown the absolute cell count of detected CD4 T cells (size) together with the frequency of Tregs within the population (indicate by colour, with percentage listed on each sample).

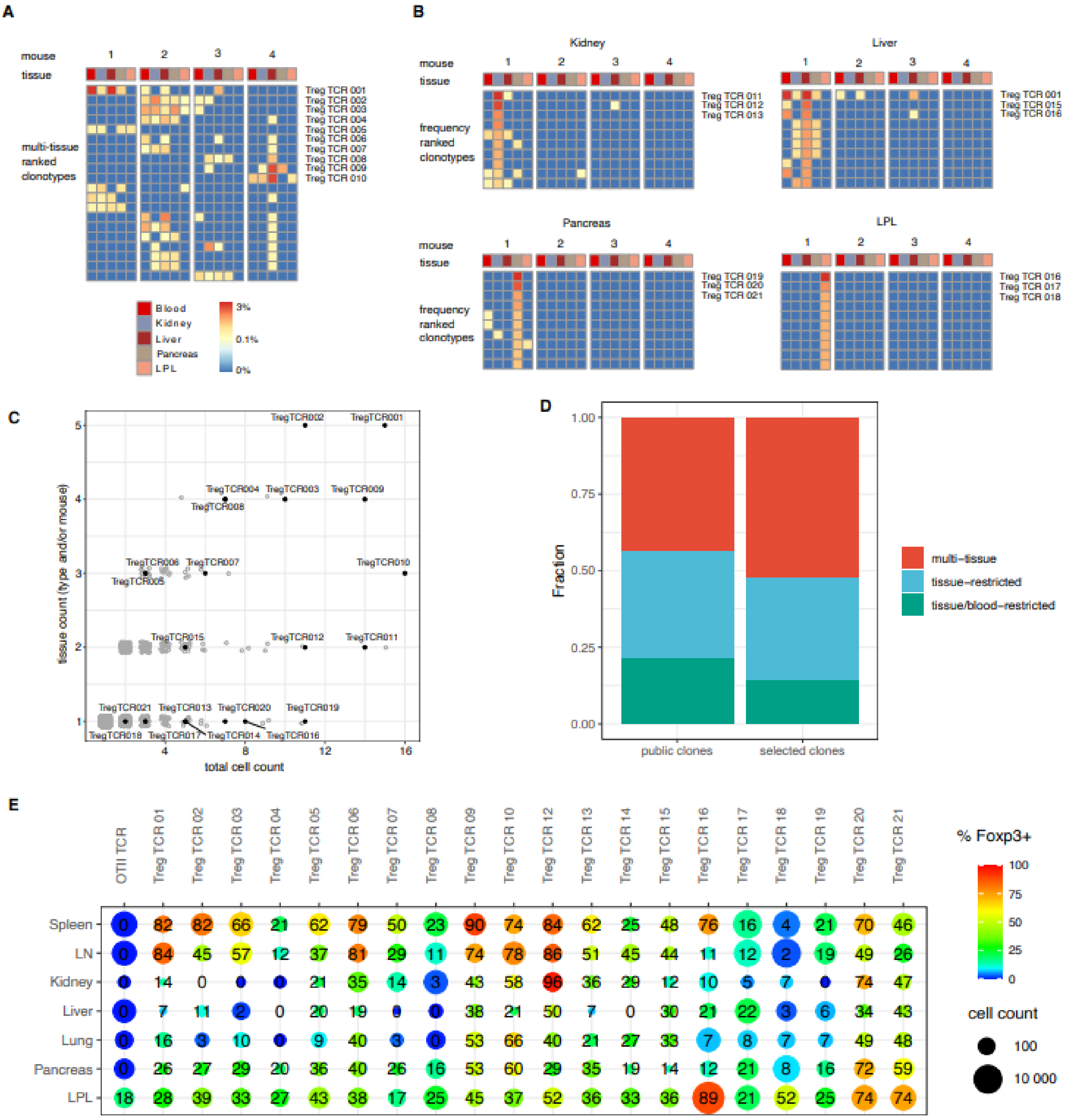
A
mouse
tissue
1
2
3
4
multi-tissue ranked clonotypes
Treg TCR 001
Treg TCR 002
Treg TCR 003
Treg TCR 004
Treg TCR 005
Treg TCR 006
Treg TCR 007
Treg TCR 008
Treg TCR 009
Treg TCR 010
Blood
Kidney
Liver
Pancreas
LPL
3%
0.1%
0%
B
Kidney
Liver
Pancreas
LPL
frequency ranked clonotypes
Treg TCR 011
Treg TCR 012
Treg TCR 013
Treg TCR 001
Treg TCR 015
Treg TCR 016
Treg TCR 019
Treg TCR 020
Treg TCR 021
Treg TCR 016
Treg TCR 017
Treg TCR 018
C
tissue count (type and/or mouse)
total cell count
TregTCR002
TregTCR001
TregTCR004
TregTCR003
TregTCR009
TregTCR008
TregTCR006
TregTCR007
TregTCR010
TregTCR005
TregTCR015
TregTCR012
TregTCR011
TregTCR021
TregTCR013
TregTCR020
TregTCR019
TregTCR018
TregTCR017
TregTCR014
TregTCR016
D
Fraction
public clones
selected clones
multi–tissue
tissue–restricted
tissue/blood–restricted
E
OTII TCR
Treg TCR 01
Treg TCR 02
Treg TCR 03
Treg TCR 04
Treg TCR 05
Treg TCR 06
Treg TCR 07
Treg TCR 08
Treg TCR 09
Treg TCR 10
Treg TCR 12
Treg TCR 13
Treg TCR 14
Treg TCR 15
Treg TCR 16
Treg TCR 17
Treg TCR 18
Treg TCR 19
Treg TCR 20
Treg TCR 21
Spleen 0 82 82 66 21 62 79 50 23 90 74 84 62 25 48 76 16 4 21 70 46
LN 0 84 45 57 12 37 81 29 11 74 78 86 51 45 44 11 12 2 19 49 26
Kidney 0 14 0 0 0 21 35 14 3 43 58 96 36 29 12 10 5 7 0 74 47
Liver 0 7 11 2 0 20 19 0 0 38 21 50 7 0 30 21 22 3 6 34 43
Lung 0 16 3 10 0 9 40 3 0 53 66 40 21 27 33 7 8 7 7 49 48
Pancreas 0 26 27 29 20 36 40 26 16 53 60 29 35 19 14 12 21 8 16 72 59
LPL 18 28 39 33 27 43 38 17 25 45 37 52 36 33 36 89 21 52 25 74 74
% Foxp3+
cell count
100
10 000
